# Systematic evaluation of the impact of promoter proximal short tandem repeats on expression

**DOI:** 10.1101/2025.09.14.676153

**Authors:** Xuan Zhang, Lingzhi Zhang, Ellice Wang, Susan Benton, Eduardo Modolo, Mikhail Maksimov, Sharona Shleizer-Burko, Qingqing Gong, Catherine Wang, Michael Lamkin, Eric Mendenhall, Melissa Gymrek, Alon Goren

## Abstract

Genetic variation at thousands of short tandem repeats (STRs), which consist of consecutive repeated sequences of 1-6bp, has been statistically associated with gene expression and other molecular phenotypes in humans. However, the causality and regulatory mechanisms for most of these STRs remains unknown. Massively parallel reporter assays (MPRA) enable testing the regulatory activity of a large number of synthesized variants, but have not been applied to STRs due to experimental and computational challenges. Here, we optimized an MPRA framework based on random barcoding to study the impact of variation in repeat copy number on expression. We first performed an MPRA on sequences derived from 30,516 promoter-proximal STR loci along with up to 152bp of genomic context, testing 3-4 variants with differing repeat copy numbers for each locus in HEK293T cells. We identified 1,366 loci with significant associations between repeat copy number and expression, which were enriched for positive effect sizes (P=2.08e-110). We then designed a second MPRA in which we performed deeper perturbations, including systematic manipulation of the repeat unit sequence, orientation, and copy number, with 200-300 perturbations for each of the 300 loci with the strongest signals. Our results revealed that the repeat unit sequence is the primary driver of differences in the relationship between copy number and expression across loci, whereas orientation and flanking sequence have weaker effects, primarily for AT-rich repeat units. The high resolution of these perturbations enabled us to detect non-linear effects, most notably for AAAC/GTTT repeats, which emerge only beyond a certain copy number threshold. Finally, we observed that a subset of STRs in our library show expression levels that are tightly linked with predicted DNA secondary structure formation. We repeated our perturbation MPRA in HeLa S3 cells under wildtype and RNase H1 knockdown conditions, which, via reduction in RNase H1 activity, are expected to hinder resolution of R-loops. This demonstrated that associations between copy number and expression at G-quadruplex-forming CCCCG/CGGGG repeats are particularly sensitive to loss of RNase H1, providing support for an R-loop mediated mechanism for these repeats. Altogether, we establish STRs as a critical component of the non-coding regulatory grammar and provide a framework for understanding how this dynamic form of genetic variation shapes gene expression.

## Introduction

Short tandem repeats (STRs), consisting of sequences of 1-6bp in tandem, represent one of the most abundant sources of genetic variation in humans^1^ and have been proposed as an important source of phenotypic variation both in humans and other species^1–4^. STRs have long been known to contribute to human disease – expansions at STRs cause dozens of diseases including Huntington’s Disease, Fragile X Syndrome, and others (reviewed in^5^) and we and others have implicated modest variation in STR copy number in a variety of traits including autism^6,7^, thyroid disease^8^, and cancer risk^9^.

We previously estimated that variation in STR copy number accounts for 10-15% of the *cis* heritability of gene expression in humans^10^. More recently, population-scale studies have identified tens of thousands of STRs acting as quantitative trait loci (QTLs) for gene expression^11–13^ as well as other molecular phenotypes, including splicing^13,14^, DNA methylation^15^, and other epigenomic phenotypes^13^. Further, genome-wide association studies (GWAS) leveraging whole genome sequencing (WGS) datasets available from large biobanks have identified hundreds of STRs for which repeat copy number is associated with complex traits^16–18^ and suggested many of these signals are mediated by STR effects on transcription. While these observations suggest a widespread contribution of STRs to a variety of phenotypes, they are based on statistical associations and in most cases do not directly provide evidence that STRs are acting as the causal variants or give immediate insight into the underlying mechanisms.

The molecular mechanisms by which most STRs influence transcriptional regulation are not fully understood. Studies on individual genes or small sets of sequences have suggested STRs may regulate transcription through multiple mechanisms, including formation of DNA or RNA secondary structures^19–21^, impacting DNA methylation^15,22^, or modifying nucleosome positioning^23,24^. Additionally, it has been suggested that STRs can influence the affinity of nearby transcription factor (TF) binding sites^25^, or even act as binding sites themselves^26,27^, and a recent *in vitro* study showed that STRs have a major influence on TF-DNA interactions^28^. However, these studies have been performed only on limited sets of STRs, and thus it is unclear if these observations are representative of genome-wide properties of different types of STRs and their functions.

Systematic interrogation of the impact of STRs on gene regulation is difficult for multiple reasons. First, synthesis and amplification of low complexity STR regions is error-prone and technically challenging. Second, certain types of STRs, including homopolymers and GC-rich repeats, face high dropout rates in next-generation sequencing (NGS) and are a large source of errors^29^. Finally, computational analysis of STR regions from NGS is challenging, largely due to the errors described above (reviewed in^30^). Further, STRs often require different analysis paradigms compared to other variants such as single nucleotide polymorphisms (SNPs) – whereas typical workflows compare the impact of single nucleotide substitutions, STR analyses often focus on associations of repeat copy number with a phenotype of interest^11,12,14–16^.

Massively parallel reporter assays (MPRAs), which involve testing the impact of thousands to millions of DNA sequences simultaneously on a reporter readout, have transformed the ability to measure regulatory activity at scale. MPRAs have proven highly successful in characterizing *cis*-regulatory regions and disease variants across diverse cellular contexts^31–33^, enabling systematic investigation of sequence-specific regulatory effects. Despite their clear promise, to our knowledge, MPRAs have not been employed to interrogate the regulatory impact of STRs, in part due to the challenges mentioned above, thus leaving a major gap in the ability to study STR-mediated gene regulatory mechanisms.

Here, we optimized an MPRA platform to overcome the challenges described above and established a framework for systematic interrogation of STR regulatory activity. We applied this platform to study the impact of copy number and repeat unit sequence on regulatory activity of 30,516 STRs occurring within 3.6kb of annotated transcription start sites. We then performed deep perturbation of 300 of these STRs to study the role of sequence composition, repeat copy number, and secondary structure formation such as G-quadruplex structures. Overall, our results provide important insights on the contribution of STRs to the regulatory non-coding grammar and provide a valuable framework to enable future high-throughput studies of the impact of STRs.

## Results

### A framework for systematically studying the regulatory effects of STRs

To assess the effects of genetic variation at STRs on gene expression, we optimized a previously described MPRA system^34^ (**Fig. 1a**). This approach amplifies an oligonucleotide pool containing sequences of interest, adding random barcode sequences to each molecule, effectively resulting in dozens to hundreds of replicate measurements per sequence. Given the observed enrichment of eSTRs (STRs for which repeat number is associated with expression of nearby genes in a population) in promoter regions^11^, we focused on STRs in close proximity to transcription start sites (TSS). We designed a pool of 100,000 oligonucleotides, each of them 230nt long, based on 30,516 unique STRs within 3.6kb of a TSS of a human protein-coding gene (**Supplementary Tables 1-2)**. Each oligonucleotide comprises an STR of interest plus up to 152nt of upstream and downstream genomic context. For each STR locus, we included variant sequences containing either the reference (hg38) STR length, 5 fewer repeat copies than hg38, or 5 more copies than hg38. For a random subset of STRs (8,452), we tested a fourth variant of 3 more copies than hg38. To maintain consistent oligonucleotide lengths during synthesis and amplification while using identical genomic sequence context for variants with different lengths at each locus, we introduced a “filler sequence” to each oligonucleotide, which is removed during the final cloning step to generate the reporter library. In total, our library consisted of 45% homopolymer, 15% dinucleotide, and 12% trinucleotide repeats, with the remainder consisting of longer repeat units (**Fig. 1**).

**Figure 1.**
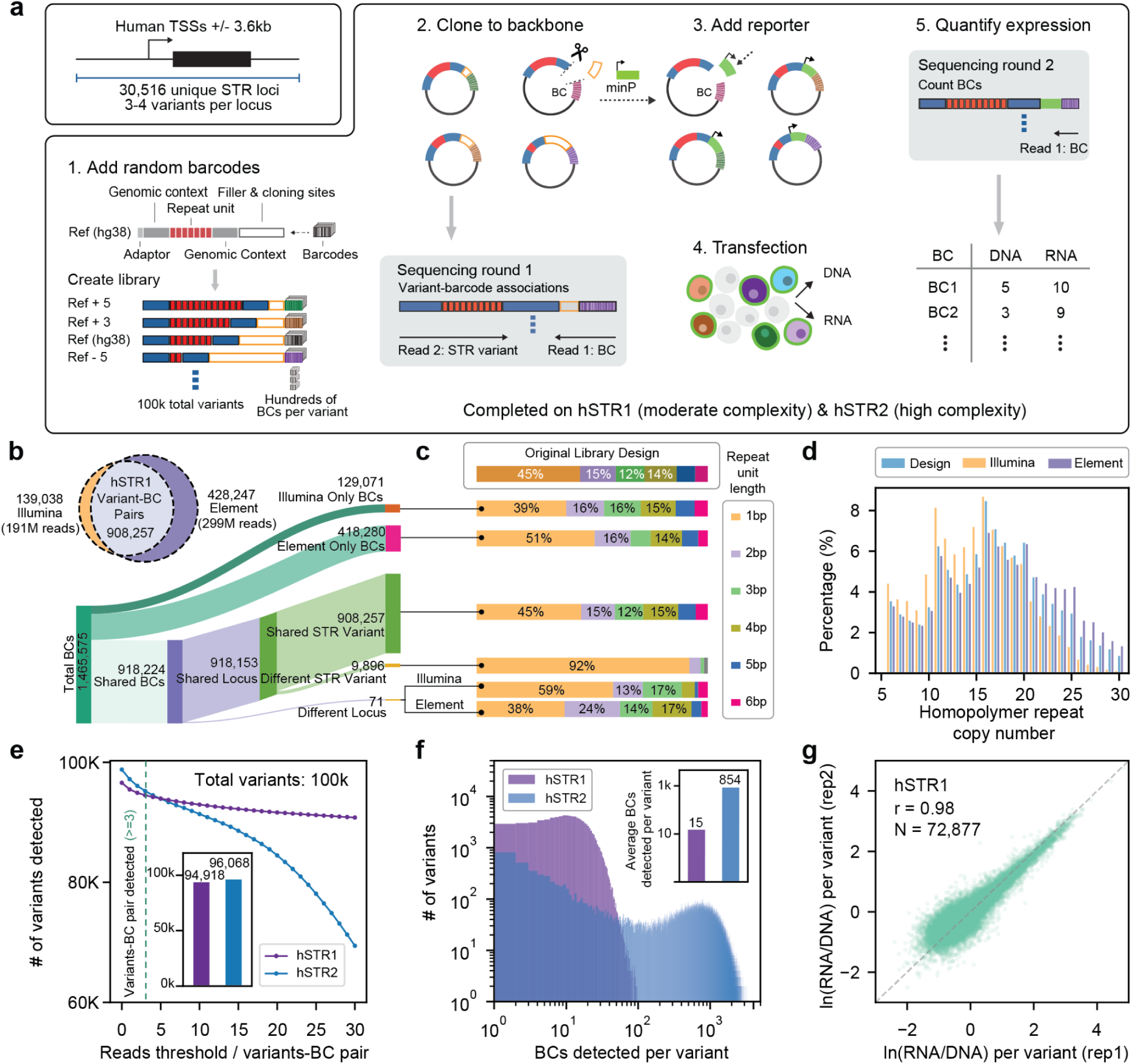
An MPRA framework to evaluate the effect of STRs on gene expression. **(a) Schematic overview of MPRA library construction and experimental workflow**. The upper left box illustrates how STR-containing loci were chosen within 3.6kb of TSSs for human genes. A total of 3-4 variants with different STR copy numbers (each red box=1 repeat unit) plus flanking genomic regions (blue) at each locus were synthesized, with an additional filler region (orange) to ensure equal length oligonucleotides. The workflow includes (1) synthesis of STR-containing oligonucleotides plus addition of random barcodes (BCs; colored boxes) via emulsion PCR, (2) cloning of the oligonucleotide library into the pGL4.23 backbone plasmid, which is sequenced (*Sequencing round 1*) to associate barcodes with variants, (3) replacing the filler region from the plasmid pool with a minimal promoter (minP; 90° arrow) and GFP reporter (green), (4) transfection of the resulting plasmid library into the target cell line followed by DNA and RNA extraction, and (5) quantification of expression based on RNA/DNA barcode counts (*Sequencing round 2*). This process was conducted separately for moderate (hSTR1) and high barcode complexity (hSTR2) libraries. **(b) Comparison of variant-BC pairs detected by NGS with Illumina and Element.** The Venn diagram depicts captured variant-BC associations using Illumina (orange) and Element (purple) for hSTR1. The Sankey diagram expands the Venn diagram to show the concordance of BC assignments by each platform at the STR locus or variant level, since in some cases we observed agreement on the STR locus but disagreement on the copy number of the variant. **(c) Proportion of detected variant-BC pairs classified by repeat unit length.** The percentages of all variants by repeat unit length in the original oligonucleotide pool design is shown in the top box. Subsequent rows show the breakdown by categories in **b.** **(d) Percentage of homopolymers by copy number detected by each sequencing platform.** The *x*-axis shows the homopolymer copy number and the *y*-axis shows the percentage of all homopolymer variants with that length in the original design (light blue), vs. those captured by Illumina (orange) or Element (blue) in hSTR1. Similar analyses for hSTR2 are shown in **Supplementary Fig. 1**. **(e) Number of variants identified after merging Illumina and Element reads.** The line graph depicts the number of variants detected (*y*-axis) for the high (blue) and moderate-complexity (purple) association libraries as a function of the threshold used for the minimum number of supporting reads required for each variant-BC association (*x*-axis). The inset bar graph shows the number of variants detected (out of 100k synthesized) using our default threshold of ≥3 reads (dashed line in main panel). **(f) Distribution of the number of BCs detected per variant in the high-complexity (blue) and moderate-complexity (purple) libraries.** The inset bar plot shows the average per library. **(g) Correlation of RNA/DNA expression ratios between replicates.** The plot compares two replicates of the hSTR1 library. Plots comparing all replicate pairs for hSTR1-2 are shown in **Supplementary Figs. 4-5**.

We amplified our oligonucleotide pool and added 20bp random barcodes to each fragment under two protocols in parallel: one with moderate barcode complexity (“hSTR1”) and a second designed to maximize barcode complexity (“hSTR2”) (**Methods**). To more accurately maintain STR lengths, emulsion PCR was used for all amplification steps. After amplification, each library was cloned into a backbone plasmid. To determine the set of barcodes associated with each unique oligonucleotide, we performed paired-end sequencing of the plasmids for each library. Due to challenges in sequencing STRs, we employed two NGS approaches: Illumina and the Element avidity system^35^, which was shown to outperform alternative technologies in challenging regions, particularly homopolymer repeats. We designed a custom pipeline to model PCR stutter errors at reads overlapping the STR sequences (**Methods**). Variant-barcode associations identified by each platform were highly concordant. Of the 1.04M variant-barcode pairs captured by Illumina, 87% were captured by the higher coverage Element dataset (**Fig. 1b**). Of barcodes identified by both technologies for hSTR1, >99% supported the same STR locus and 98.9% supported identical variants. Discordant barcodes assigned to different variants, as well as those identified only by Element, were strongly enriched at homopolymer loci (**Fig. 1c**). More detailed inspection of homopolymer sequences showed that those identified by Illumina were depleted for long variants (>20bp) whereas those sequences were well-captured by Element (**Fig. 1d**). Overall, these results suggest Element sequencing is able to detect long homopolymer sequences that could not be captured by Illumina. Similar trends were observed for hSTR2 (**Supplementary Fig. 1a-b**), but with reduced overlap between detected variant-barcode pairs driven by the higher complexity and reduced sequencing saturation of that library (**Supplementary Fig. 1c**).

Sequencing reads from each platform (total 191M/494M by Illumina and 299M/839M by Element for hSTR1/hSTR2) were merged to determine a final set of variant-barcode associations for each library. After filtering (**Methods; Supplementary Fig. 2**), we identified 1,376,241 (hSTR1) and 82,095,183 (hSTR2) unique variant-barcode pairs which captured 94,918 (hSTR1) and 96,068 (hSTR2) of the original 100,000 variant sequences (**Fig. 1e**). Each variant was associated with an average of 15 (hSTR1) and 854 (hSTR2) unique barcodes (**Fig. 1f**). Undetected variants in both libraries were enriched for repeats containing stretches of G or C homopolymers (**Supplementary Fig. 3)**.

After inserting a minimal promoter (minP) and green fluorescent protein (GFP) reporter, we transfected the hSTR1 and hSTR2 libraries into HEK293T cells in triplicate (three separate dishes). Following transfection, we extracted DNA and RNA. RNA was reverse transcribed into cDNA, but for simplicity we refer to it as RNA below. We performed targeted amplification of the barcode region in both the RNA and DNA prior to NGS. Read counts for each barcode for both DNA and RNA were highly concordant across replicates within each library (Pearson *r* range 0.98-0.99 for DNA, all 1.00 for RNA; 0.98-0.98 for RNA/DNA ratios; **Fig. 1g; Supplementary Figs. 4-5**). Resulting read counts were used for downstream analyses of expression performed below.

### STRs modulate gene expression in a sequence-dependent manner

Before testing for impacts of repeat copy number, we first performed an initial assessment of the overall ability of our STR containing variants to impact GFP expression (**Fig. 2a**). We filtered the data to include only variants with at least three unique barcodes and a minimum of 10 reads in both the RNA and DNA libraries (**Methods**). We observed strong expression signals for most variants, as evidenced by enrichment of RNA vs. DNA counts (**Supplementary Fig. 6**), with consistent RNA/DNA ratios between the hSTR1 and hSTR2 libraries (**Fig. 2b**; Pearson r=0.90; two-sided P<10e-200). We used the distribution of Z-normalized log_10_(RNA/DNA) values to characterize the overall expression level of each variant. Altogether, we found that variants with GC-rich repeat units tended to have higher expression compared to variants with AT-rich repeats (**Fig. 2c**). The median Z-scores across repeat units are highly concordant across hSTR1 and hSTR2 (**Fig. 2d**; Pearson r=0.92; two-sided P=2.78e-167). Overall, these findings demonstrate that the synthesized STR variants are impacting transcription activity, and suggest a strong relationship between the GC content of STRs and their impact on gene expression, with GC-rich repeat units generally associated with increased expression and AT-rich units associated with decreased expression.

**Figure 2.**
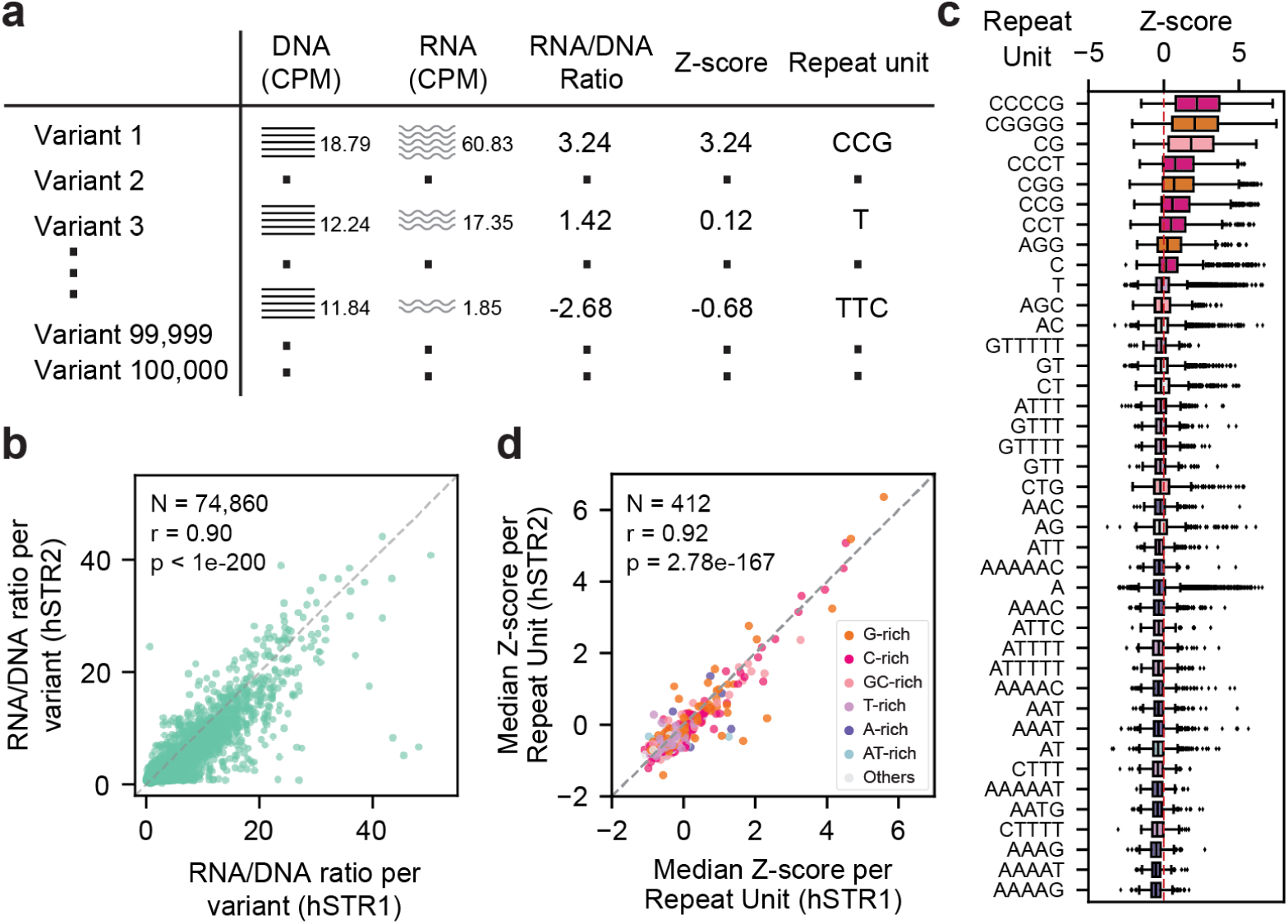
Characterization of STR variants with high vs. low expression in our MPRA framework. **(a) Schematic overview of expression results.** For each STR variant, we quantified DNA and RNA abundances (counts per million; CPM), the RNA/DNA ratio, Z-score of the ratio relative to all other variants, and repeat unit sequence. RNA abundances are represented as wavy lines. **(b) Comparison of RNA/DNA measurements across moderate- and high-complexity association libraries.** Each dot represents a single variant and is based on summing RNA and DNA read counts across each barcode associated with each variant merged across all replicates. The dashed line indicates the *x*=y diagonal. **(c) Distribution of expression values for each repeat.** The boxplot shows the distribution of expression Z-scores for all variants with each repeat unit. Pink and orange represent G/C rich repeat units while blue and purple represent A/T rich repeat units. **(d)** Concordance of expression values for each repeat unit between the moderate complexity hSTR1 (*x*-axis) vs. high complexity hSTR2 (*y*-axis) libraries. Expression is summarized for each repeat unit as the median Z-score across variants containing that unit. Color denotes the category of repeat unit. The dashed line indicates the *x*=y diagonal. Only repeat units with at least 200 variants included in the analysis are shown in **c-d**.

### The impact of variation in STR copy number on reporter expression

After demonstrating above that STR-containing variants can drive GFP expression, we next tested for association between the number of copies of the repeat unit (copy number) for the variants from each locus and reporter expression (RNA/DNA ratio) using a linear model accounting for experimental replicate as a covariate (**Fig. 3a**). After filtering variants with low barcode counts and loci with data for fewer than three unique variants (**Methods**), 19,818 loci from the hSTR1 library remained for analysis. Of these, 1,366 loci showed a significant association between copy number and expression at a false discovery rate (FDR) of 10% (**Fig. 3b**). Notably, variants from loci with significant associations tended to have higher overall expression compared to variants from loci without significant associations (**Supplementary Fig. 7**; Mann-Whitney one-sided P=7.84e-222), which likely reflects higher power to detect significant associations when at least one variant is highly expressed. We observed a strong bias toward positive correlation between repeat length and expression (n=1,083 loci) compared to a negative correlation (n=283 loci) (binomial test two-sided P=2.08e-110). We repeated the regression analysis for the hSTR2 library, and found a strong correlation of effect sizes between the hSTR1 and hSTR2 libraries (Pearson r=0.92; two-sided P=1.90e-123; n=292 loci identified as significant in both) and a similar bias toward positive effect sizes (binomial test two-sided P=3.54e-98; **Supplementary Fig. 8**). Effect sizes measured by MPRA are not significantly correlated with eSTR effect sizes we previously measured in humans^11,36^ (**Supplementary Fig. 9**). We hypothesize this low correlation is largely driven by the non-overlapping repeat ranges tested in each, different cell types, and dropout in our library of many GC-rich repeats with strongest effects in humans (**Discussion**).

**Figure 3.**
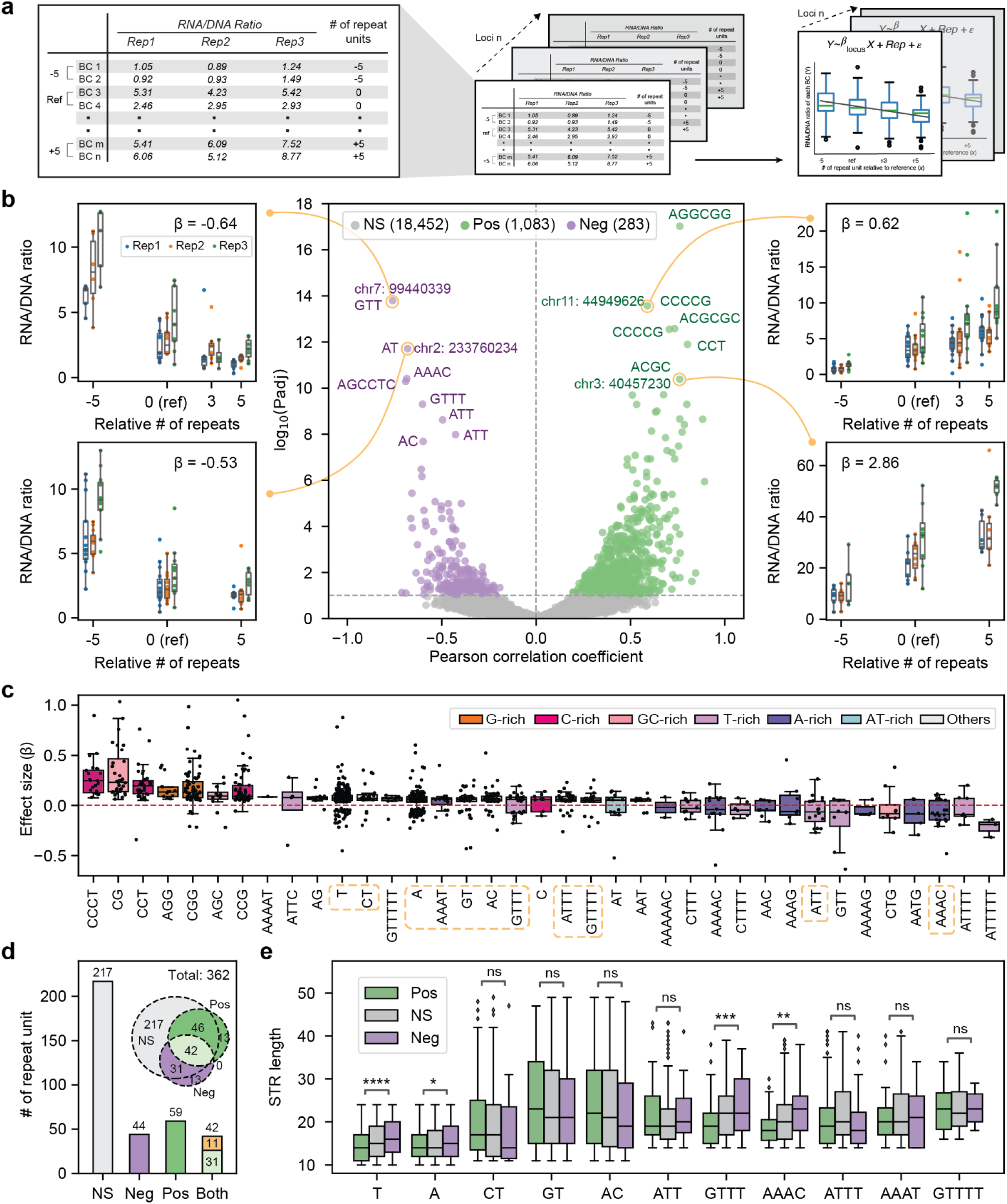
Relationship between STR copy number and MPRA reporter activity across loci. **(a) Schematic representation of expression library barcode count analysis.** For each locus, we compute RNA/DNA ratios as well as the repeat copy number for the variant associated with each barcode across all three replicates. The left panel shows example data for a single locus. We perform a separate regression analysis between repeat copy number and RNA/DNA ratios for each locus, accounting for replicate as a covariate. **(b) Volcano plot summarizing associations between repeat copy number and reporter expression at each locus in hSTR1**. The *x*-axis shows the Pearson correlation between copy number and expression. The *y*-axis shows the log10 of the adjusted P-value. Selected loci are annotated with the hg38 locus coordinates and repeat unit. Purple=loci with a negative correlation between copy number and expression, green=loci with a positive association, grey=not significant. Representative plots are shown for two loci with positive and two with negative associations. For these, repeat copy number on the *x*-axes is shown relative to the reference genome (reference denoted by “0”). The *y*-axes denote RNA/DNA ratios. Each data point represents one barcode in one replicate and boxplots summarize distributions within each replicate. Regression slopes (β) are annotated for each locus. Blue=replicate 1, orange=replicate 2, green=replicate 3. **(c) Distribution of regression effect sizes (β) for significant loci, grouped by repeat unit type.** Only repeat units for which more than 50 loci were tested in the regression analysis are shown. Each dot represents a single locus. Boxplots summarize the distribution of values across loci for each repeat unit. Dashed orange boxes on the *x*-axis indicate repeat units with more than 15 loci with positive and more than 15 with negative effects. **(d) Summary of effect directions for each repeat unit.** Bars show the number of unique repeat units for loci with no significant effect (grey), negative effects (purple), positive effects (green), or both positive and negative effects (light green and orange). The inset Venn diagram indicates the overlap of repeat units across categories. Repeat units highlighted in orange in **(c)** are also emphasized in orange within the “both” bar category. **(e) Distribution of reference repeat length for selected repeat units that display both positive and negative associations (bivalent).** Boxplots show the total repeat length (in bp) in hg38 for loci with positive (green), not significant (grey), or negative (purple) effects on expression. Only loci for which at least 15 loci each with positive and negative effects were identified are shown. ns=not significant; *=1.00e-02 < p ≤ 5.00e-02; **=1.00e-03 < p ≤ 1.00e-02; ***=1.00e-04 < p ≤ 1.00e-03; ****=p ≤ 1.00e-04.

We next examined whether effect sizes and directions differed across STRs with different repeat unit sequences (**Fig. 3c-d**). Due to the reduced power resulting from lower read counts for each barcode in hSTR2 (**Supplementary Fig. 2b**), the results below focus on the hSTR1 library. Overall, we found that STRs with GC-rich repeat units tended to have a strong bias toward positive effects. In contrast, effect sizes tended to shift toward negative values for more AT-rich units, a trend illustrated by four representative examples in **Fig. 3b**. The observation that GC-rich and AT-rich repeat lengths have positive and negative correlations with expression, respectively, adds to our findings above, in which GC-rich repeat units tend to show higher overall expression compared to AT-rich units (**Fig. 2**).

Of 135 unique repeat units included in our library for which at least one locus showed a significant effect of copy number on expression, 59 units showed only positive effects and 44 showed only negative effects (**Fig. 3d**). Notably, 42 repeat units exhibit a bivalent trend, with some loci showing positive and others showing negative effects (**Fig. 3c-d**). To further investigate this phenomenon, we selected 11 repeat units (highlighted in **Fig. 3c-d**) from this set with a sufficient number of loci (n ≥ 15 positive and n ≥ 15 negative) and compared the STR length of the reference variants between loci with positive vs. negative effects (**Fig. 3e**). For a subset of repeats with significant differences, loci with positive effects have consistently shorter STR regions compared to those with no effect, whereas those with negative effect sizes tend to have longer STR regions (**Fig. 3e**). This finding is consistent with a non-linear effect of some repeat units on expression, in which up to an optimal repeat length, increasing copy number increases expression, after which increasing copy number decreases expression. We investigate these and other trends identified in our initial library below by incorporating additional variants spanning a broader range of copy numbers for a subset of loci.

### Deep perturbation of candidate regulatory STRs

To gain deeper insights into the STRs with strongest regulatory effects, we designed a second array, focusing on the most significant loci from our hSTR datasets. We selected 200 and 100 loci with strong positive and negative effects, respectively, between copy number and expression (**Fig. 4a**). For each selected locus, we implemented three distinct types of perturbations. First, we varied the number of repeat units, creating sequences containing 0 to up to 23 repeats for homopolymers and up to 15 repeats for other unit types (**Fig. 4b**; left). Second, we systematically substituted the original repeat unit with various sequence types, including alternative repeat units (up to one alternative unit each from homopolymers, dinucleotides, trinucleotides, or tetranucleotides, and including the reverse complement of the original unit), testing all length variants considered for each locus (**Fig. 4b**; middle). We additionally tested non-STR random sequences, and non-STR random sequences with matched GC-content to the original repeat unit, considering lengths equal to multiples of three copies of the original repeat unit (**Methods**; **Supplementary Table 3**). Third, we included both the forward and reverse orientations for each region tested (**Fig. 4b**; right). Finally, we additionally included any original hSTR variants from all 300 selected loci that were not already included as part of this design (see **Methods**) as internal controls. This comprehensive design generated a total of 59,842 variants with a median of 205 (range 90-289) variant sequences per locus (**Fig. 4a-c**). Below we refer to this oligonucleotide pool and its corresponding libraries as dpSTR (deep perturbation of STRs). Similar to procedures established for hSTR1 and hSTR2, we performed emulsion PCR to amplify the oligonucleotide pool and add 20bp random barcodes, and cloned the library to a backbone plasmid. We then used NGS to determine associations between random barcodes and synthesized variants.

**Figure 4.**
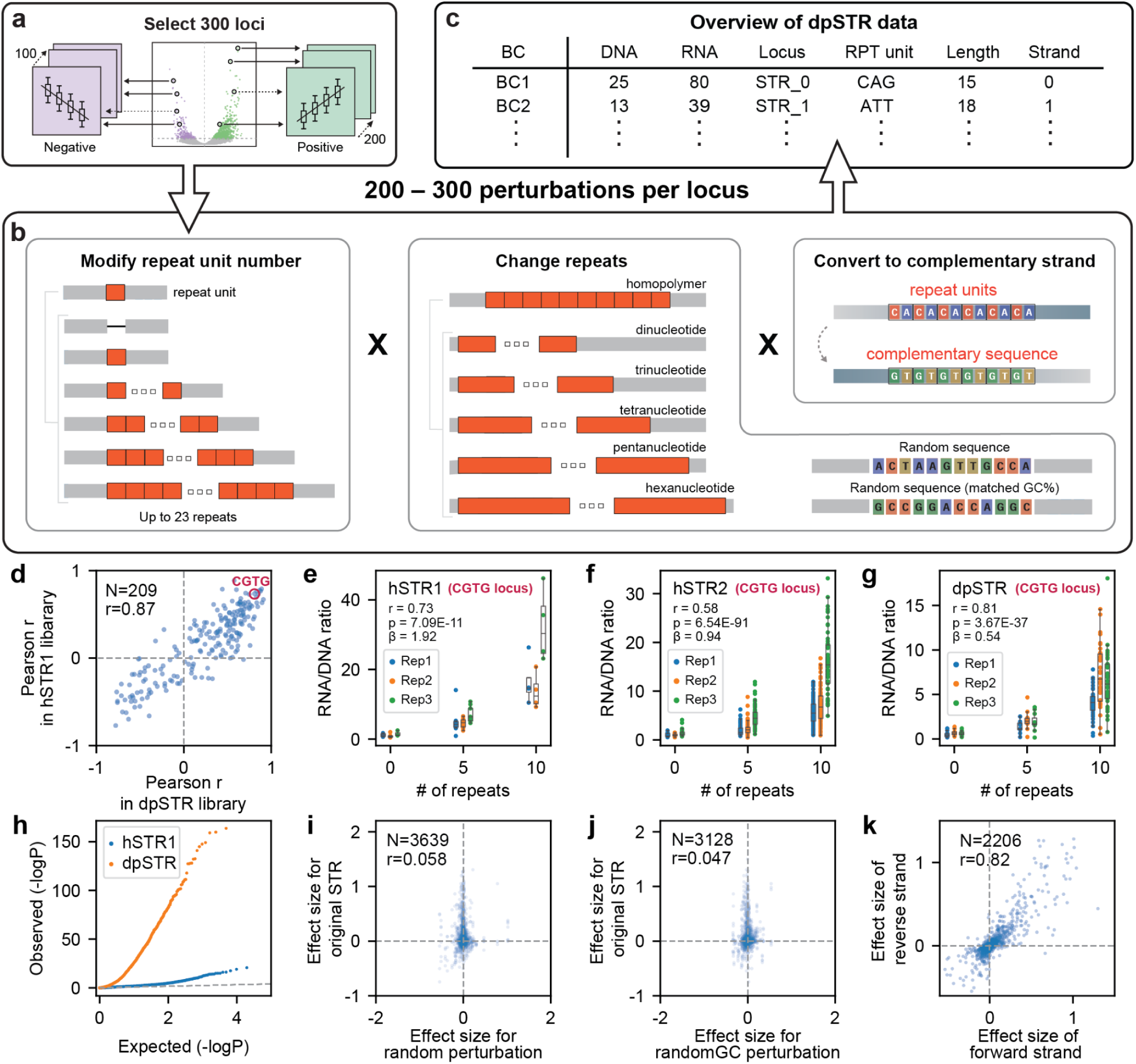
Deep perturbation analysis of candidate regulatory STR loci. **(a) Overview of locus selection for the dpSTR (deep perturbation of STRs) library.** We selected 300 loci with strong effect sizes (200 positive, 100 negative) from the analysis of the moderate-complexity hSTR1 library. **(b) Overview of perturbation design.** For each locus, we modified repeat number (left), repeat unit sequence (middle), and strand orientation (top right). We also replaced each repeat with variable length random sequences (bottom right). For each locus, the repeat unit is represented in orange and flanking genomic context in grey. A list of all designed oligonucleotides for dpSTR is given in **Supplementary Table 3**. **(c) Overview of data derived from the dpSTR library.** After cloning and transfection, similar to hSTR1 and hSTR2 a barcode count table is generated with the associated locus and perturbation and its corresponding RNA/DNA ratio. **(d) Comparison of correlation coefficients of original and perturbation libraries.** Correlations are measured between copy number and expression for each variant using Pearson r. The plot includes loci passing QC steps in both libraries. The locus circled in dark pink with reference repeat unit CGTG is shown in detail in **e-g**. **(e–g) Representative plots for one locus illustrating repeat length versus RNA/DNA ratio across moderate-complexity hSTR1 (e), high-complexity hSTR2 (f), and deep perturbation libraries (g).** Repeat copy number on the *x*-axes is shown relative to the reference genome (reference denoted by “0”). The *y*-axes denote RNA/DNA ratios. Each data point represents one barcode in one replicate and boxplots summarize distributions within each replicate. Regression effect sizes (β), nominal P-values, and Pearson r are annotated for each locus. Blue=replicate 1, orange=replicate 2, green=replicate 3. **(h) Quantile-quantile plot comparing the distribution of regression P-values between hSTR1 and the dpSTR libraries.** Blue=hSTR1, orange=dpSTR. **(i–k) Comparison of regression effect sizes for paired perturbations at each locus.** Plots compare effect sizes when replacing the repeat sequence of each locus with random sequences (**i**), random sequences with matched GC content (**j**), or flipping the orientation (**k**) of each locus. Each dot represents a single locus in the dpSTR library.

The high sequence similarity between different variants included in dpSTR presented substantial technical challenges. For example, dpSTR includes both the forward and reverse complement of each tested sequence as separate variants which are not disambiguated by standard aligners and high-resolution of the range of copy numbers tested, with many variants differing from the next closest sequence by a single repeat unit copy. In addition to experimental optimization detailed for both oligonucleotide pools in **Methods**, this second pool necessitated development of a novel analysis pipeline for assigning barcodes to variant sequences (**Methods**). First, we implemented a multi-stage alignment strategy to disambiguate between sequences from the forward vs. reverse design of each variant. Second, we implemented a maximum likelihood estimation procedure to model PCR stutter errors, which can result in addition or deletion of one or more repeat units. These errors can falsely lead to the same barcode being associated with multiple distinct variants, particularly when synthesized variants differ in length by a single repeat, which was not the case for hSTR1 and hSTR2 for which all variants differed by at least two repeat units. Following correction, barcodes which had been associated with two or more variants (30% of total barcodes; **Supplementary Fig. 10a-c**) were uniquely assigned. Our refined pipeline identified 57,735 out of 59,842 synthesized variants with an average of 59 barcodes per variant (**Supplementary Fig. 10d**).

In a similar manner to hSTR1 and hSTR2, we inserted a minimal promoter and GFP reporter, transfected the resulting plasmid library into HEK293T cells in triplicate, and extracted and sequenced RNA and DNA from the barcode region (**Fig. 4c**). We first tested whether RNA/DNA ratios obtained from dpSTR recapitulated previously observed associations between repeat copy number and reporter expression at each internal control locus. Correlations between copy number and expression at each locus were highly consistent across the hSTR1 and dpSTR (Pearson r=0.87; n=209 loci; two-sided P=1.75e-64 **Fig. 4d**; example locus shown in **Fig. 4e-g**), confirming the robustness of expression patterns across both libraries. Association statistics were far stronger in dpSTR compared to hSTR1 (**Fig. 4h**) when considering all loci in each library, which is expected since this second library focuses on loci with the strongest signals in hSTR1.

We next performed separate association tests for each perturbation (locus/sequence type pair, where sequence type indicates either different repeat unit or strand) and evaluated the impact of each one on the associations of copy number vs. expression. First, we tested whether the impact of copy number on expression can be explained by sequence length alone vs. by the specific STR sequence. We replaced STR regions with random sequences of identical length while maintaining the same flanking regions. This replacement eliminated most expression effects (**Fig. 4i**), even when controlling for GC content (**Fig. 4j**), demonstrating that the STR sequence itself, rather than just its length, drives the observed expression changes. In particular, for the majority of loci tested here, replacing the STR with random sequence resulted in effect sizes close to 0, suggesting that the presence of a repeat sequence is required to drive observed associations between copy number and expression. Next, we tested the impact of strand orientation. We observed that effect sizes at each locus are highly concordant between forward and reverse strand variants (Pearson r=0.82; n=2,206 locus/sequence type pairs; two-sided P<10e-200; **Fig. 4k**), indicating that most promoter STRs function symmetrically in our system, consistent with findings from another MPRA study showing that orientation has a modest impact on reporter expression^37^. We then compared variants for which the repeat unit was deleted entirely (copy number=0) to the reference variant at each locus (**Supplementary Fig. 11**) and found that this recapitulated trends consistent with those observed in hSTR1 and 2: deletion of GC-rich repeats or other repeats which showed generally positive effect sizes (**Fig. 2c**) tended to decrease expression, whereas deletion of repeats that showed primarily negative associations between copy number and expression (AAT and AAAG repeats) resulted in increased expression. Finally, we repeated the dpSTR MPRA experiment in HeLa S3 cells, and found that effect sizes for each locus/repeat unit pair were highly concordant with those observed in HEK293T (Pearson r=0.83; n=4,772; P<10e-200; **Supplementary Fig. 12**).

### STR regulatory effects are driven by a combination of repeat unit and flanking sequence

We next evaluated the impact of specific repeat units on associations between repeat copy number and expression. By design, at each locus our array includes both the reference repeat unit as well as alternative repeat units with varying length, with a total of 163 unique repeat units considered. For each unique repeat unit, we captured the majority of loci which either contained that unit originally or in the perturbation design (**Fig. 5a**, which includes the 33 unique units with more than 20 related variants). For each repeat unit, for each locus where the unit was included (which we define as a repeat unit/locus pair), we performed separate linear association tests between copy number and expression. When inspecting individual examples, we found that loci with the same repeat unit tended to show similar association trends (**Supplementary Fig. 13**), regardless of the flanking region and of the trend originally observed at each locus when considering the reference repeat unit sequence. This suggests that repeat unit sequence is a major driver of the associations observed in our MPRA setup.

**Figure 5.**
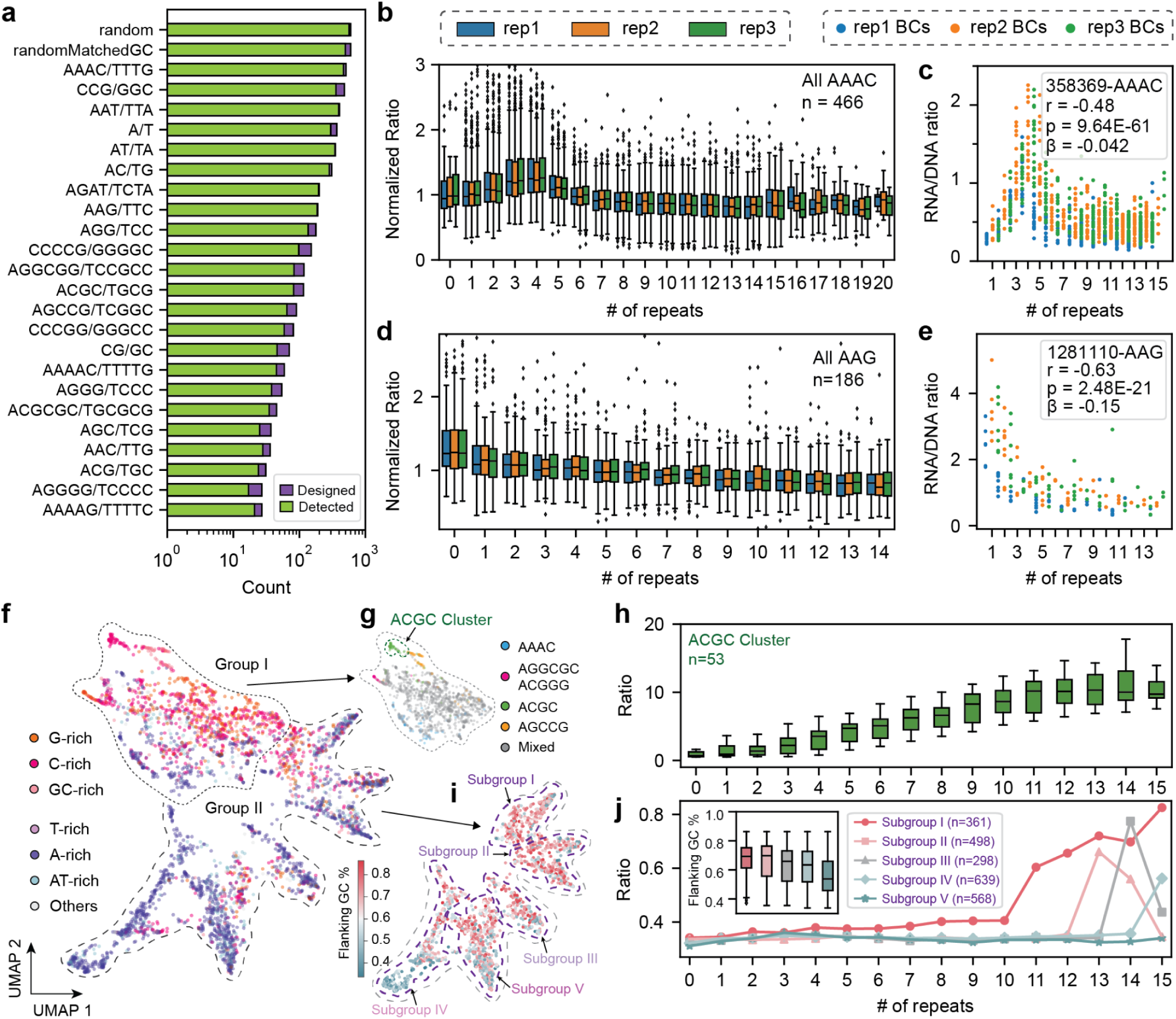
Repeat unit and flanking GC-content impact expression dynamics. **(a) Overview of repeat unit representation in the dpSTR library.** Purple bars show the number of variants in the dpSTR library design with each repeat unit. Green bars, which are overlaid on the purple bars, represent the number of variants that were detected after quality filtering. The *x*-axis is shown on a log scale. **(b) Distribution of normalized expression (RNA/DNA ratios) by copy number across AAAC repeats.** All loci for which AAAC repeats were included as either the reference repeat unit or one of the perturbations are included. Color indicates replicate number (blue=replicate 1, orange=replicate 2, green=replicate 3). The normalization procedure is described in **Methods.** Note, some loci might be tested with the same repeat unit twice if the reference contains that locus with imperfections, since we tested with and without the imperfection (**Methods**). **(c) Representative relationship between copy number and expression for an example AAAC repeat.** The *x*-axis gives the total repeat copy number. The *y*-axis denotes RNA/DNA ratio. Each data point represents one barcode in one replicate. Locus id, regression slope (β), nominal P-value, and Pearson r are annotated. Blue=replicate 1, orange=replicate 2, green=replicate 3. The locus originates at chr13:27450197-27450212 (hg38). **(d-e**) are the same as **b-c** but for AAG repeats. The example locus is from chr6:143450810-143450837 (hg38). **(f) UMAP visualization of expression patterns in the dpSTR library.** Each dot represents a single repeat unit/locus pair. Points are colored by repeat unit category. The UMAP projection was manually partitioned into <u>Group I</u> (primarily GC-rich repeats) and <u>Group II</u> (primarily AT-rich repeats). **(g) Visualization of Group I by repeat unit.** Keeping the UMAP pattern of Group I from (**f**), the colors indicate the repeat unit sequence of each repeat unit/locus pair. Green=ACGC; blue=AAAC; pink=AGGCGC and ACGGG; yellow=AGCCG; other=grey. Points in the other category did not group by repeat unit. **(h) Expression distribution by copy number.** RNA/DNA ratios are plotted by copy number across the ACGC cluster. The boxplot shows un-normalized barcode-level ratios for variants highlighted in green in panel **g.** **(i) Visualization of Group II by flanking GC content.** Keeping the UMAP pattern of Group II from (**f**), the colors indicate the GC % of the flanking sequence. Higher flanking GC content is represented by red and lower flanking GC % is represented by blue. Subgroups are based on manual inspection of the UMAP projection. **(j) Median RNA/DNA expression ratio across repeat unit/locus pairs for each subgroup of Group II.** For each subgroup identified by flanking GC % (panel **i**), the plot shows the median RNA/DNA ratio for the repeat unit/locus pairs within that subgroup. Colors correspond to subgroup classification as in panel **i**. The inset boxplot depicts the distribution of flanking GC % for repeat unit/locus pairs within each subgroup.

To systematically investigate the impact of the repeat unit sequence, we aggregated data for each unit across all loci analyzed and visualized normalized expression patterns as a function of repeat copy number (**Methods**). The expanded range of lengths considered in the dpSTR design, which were identical across all loci tested for each repeat unit, revealed strong non-linear trends for multiple repeat units (**Fig. 5b**; **Supplementary Fig. 14**). For example, for the most abundant repeat unit perturbation, AAAC/GTTT, we observed a positive effect of copy number on expression for up to 4 copies, a negative effect for 4-7 copies, and no trend for 8+ copies (**Fig. 5b-c**). This trend, which we hypothesize is related to binding of FOX-family transcription factors^27,38^ (see **Discussion**), is consistent across nearly all loci for which the repeat unit was originally or substituted to be AAAC (examples in **Supplementary Figs. 13c,14**). This pattern was not observable at individual loci in the original hSTR1 library, which contained only 3-4 unique repeat copy numbers per locus and for which lengths could vary across loci. However, the finding of a non-linear effect here is consistent with our observations in aggregate from that library (**Fig. 3e**), in which AAAC repeats showed a bivalent direction of effect. In that library, loci showing positive associations tended to have shorter average length (aligning with the range of increasing trend in **Fig. 5b**) whereas those showing negative associations tended to be longer (aligning with the range of decreasing trend). We observed that most other repeat units with strong consistent trends, in particular those with high GC percentages, tended to exhibit positive associations across the full range of repeat numbers tested. However, some units, including AAG (**Fig. 5d-e**) and AAT (**Supplementary Fig. 14**) showed consistent negative associations.

To further identify sequence properties driving changes in observed expression patterns across variants in dpSTR, we performed dimensionality reduction on the matrix of expression values for each repeat unit/locus pair vs. copy number using Uniform Manifold Approximation and Projection (UMAP) to embed each pair in two-dimensional space (**Fig. 5f**). Because not all copy numbers were designed or detected for all repeat units included in dpSTR, prior to downstream analysis we first repeated the UMAP procedure restricting it to a maximum of either 15, 18, or 21 repeat copies to ensure that the overall patterns observed are robust (**Supplementary Fig. 15**). This analysis revealed two major groups primarily containing GC-rich repeat units (Group I) vs. AT-rich units (Group II), consistent with our initial observations in hSTR1, which showed a strong impact of GC content of the STR on observed expression patterns. Within the GC-rich group, the repeat unit strongly influenced clustering. For example, the units AGGCGC, ACGGG, and AAAC formed distinct subclusters, with many exhibiting generally positive associations (e.g., ACGC shown in **Fig. 5h**).

By contrast, Group II, which consists mostly of AT-rich repeats, largely did not cluster by repeat unit (**Supplementary Fig. 16**). Rather, the GC content of the surrounding genomic context of each repeat unit/locus pair was more strongly associated with observed subgroups defined by manual inspection. We identified a gradient from higher to lower flanking GC content across subgroups I-V (**Fig. 5i-j**). Further, we observed that an increase in the flanking GC content corresponded to upregulation of expression at shorter repeat copy numbers (**Fig. 5j**; **Supplementary Fig. 17**). Together, these findings reveal distinct patterns of STR copy number vs. expression associations that are driven by an interplay of repeat unit composition, length, and genomic context. Specifically, the majority of GC-rich repeats show trends primarily driven by the repeat unit, whereas trends at AT-rich repeats are better explained by differences in flanking GC content.

### STRs predicted to form G-quadruplexes are uniquely sensitive to RNAse H1 knockdown

Motivated by previous studies linking the ability of STRs to form unique DNA or RNA secondary structures such as G-quadruplexes (G4)^19^, Z-DNA^21^, or other stable conformations^39^, we investigated whether our MPRA observations could be explained, at least in part, by such structural or conformational changes. To this end, we applied an array of computational tools to predict G4 and Z-DNA formation and DNA/RNA folding energy (**Methods**) for each variant, and assessed the correlation of each of these scores with observed expression (RNA/DNA ratio) values (**Supplementary Fig. 18**).

While Z-DNA scores and folding energies showed correlations with multiple repeat units, they lacked specificity to any particular repeat unit types. In contrast, G4 structures demonstrated a significant and specific correlation with expression levels for the repeat units CCCCG/CGGGG and AGGGG/TCCCC (e.g. Pearson r=0.34 and 0.18, two-sided P=2.43e-25 and P=0.032 between these two repeat units and the sum of G4 scores, **Supplementary Fig. 18**). In our MPRA system, these repeats can both potentially form G4 structures on either the forward or reverse strand depending on which strand the G-rich orientation of the repeat unit is (e.g. CGGGG rather than CCCCG; **Fig. 6a**). Given existing evidence linking G4 structures to promoter regulation^40^, and our previous work demonstrating STRs predicted to form G4s are strongly enriched among fine-mapped eSTRs^11^, we focused our investigation on G4-mediated regulation. Notably, in both our eSTR and MPRA data, STRs predicted to form G4s predominantly show positive associations between copy number and expression.

**Figure 6.**
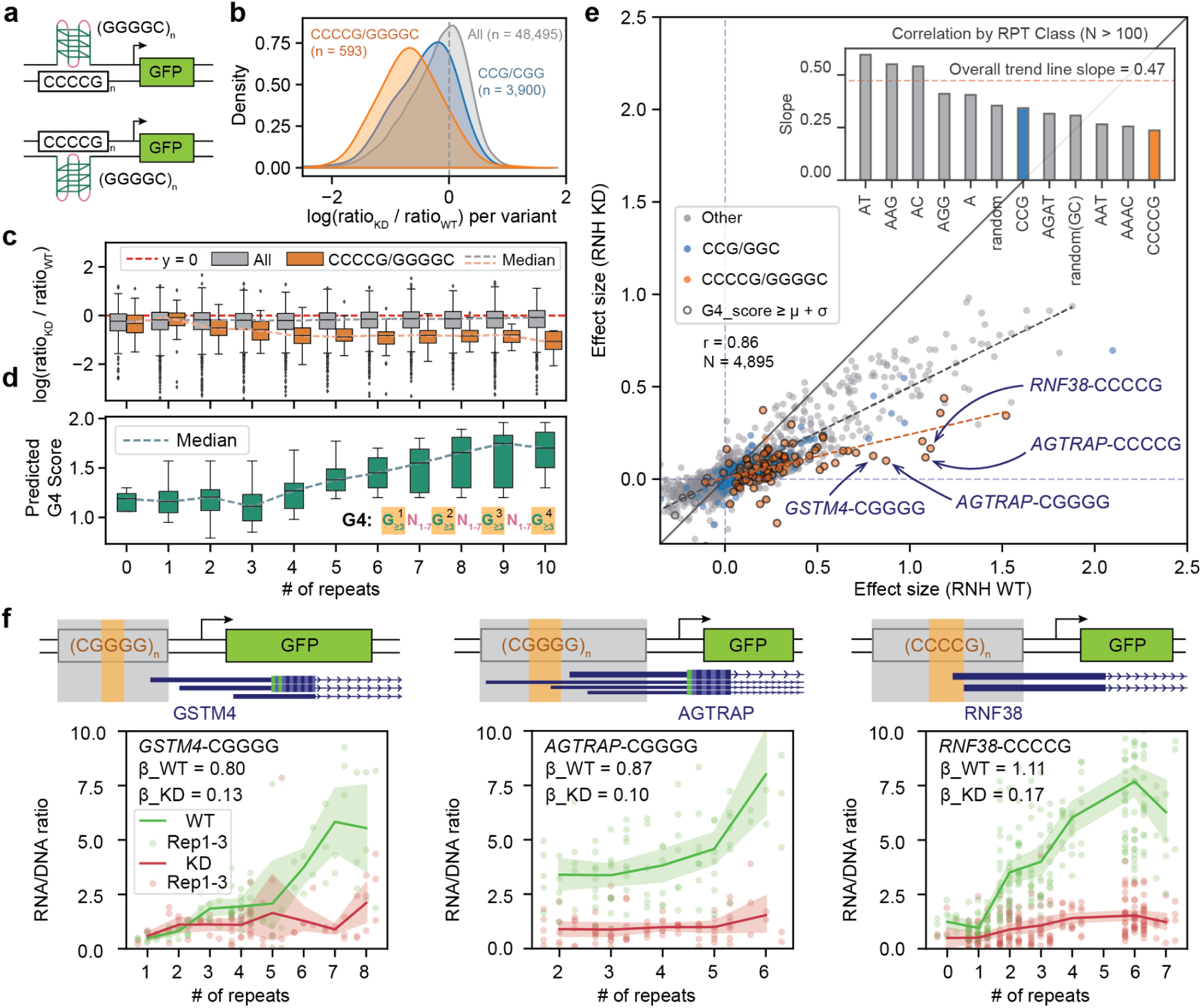
G-quadruplex formation drives R-loop sensitivity in STR regulation. **(a) Schematic illustrating the stranded G-quadruplex (G4) structure formed by CCCCG/CGGGG repeats in the STR region upstream of the reporter.** Green boxes denote GFP and arrows represent the promoter and transcription direction. **(b) Distribution of variant-level log(KD/WT) expression ratios.** Distributions are shown for all variants (grey), repeat unit/loci pairs with CCG/CGG repeats (blue) and repeat unit/loci pairs with CCCCG/CGGGG (orange) repeats. KD=knockdown; WT=wild type. **(c) Distribution of variant-level log(KD/WT) expression ratios stratified by repeat copy number.** Orange=CCCCG/CGGGG repeat variants; grey=all detected variants. The y=0 line is represented in black, the median for all detected variants in dashed grey, and the median for CCCCG/CGGGG repeat variants in dashed orange. **(d) Distribution of predicted G4 scores for detected variants containing the CCCCG/CGGGG repeat unit, stratified by repeat copy number.** The G4 consensus sequence is annotated in the bottom right. The dashed line shows the median predicted sum of G4 scores for each repeat copy number. **(e) Comparison of regression effect sizes for association tests between repeat copy number and expression for the WT vs. RNase H1 KD (RNH) knockdown conditions.** Each dot represents one repeat unit/locus pair. Blue=CCG/CGG repeats; orange=CCCCG/CGGGG repeats; grey=other repeats. Data points with G4 score ≥ µ + σ are highlighted by a black circle. The *y=x* line is represented as a solid black line, the best fit line for all data points is represented as a dashed black line, and the best fit line for all data points with CCCCG/GGGGC repeat units is represented as a dashed orange line. Selected loci with the CCCCG/GGGGC repeat unit (also depicted in **f**) are annotated according to the proximal gene. The inset bar plot shows the Pearson correlation between effect sizes stratified by repeat unit. The red dashed line shows the correlation when considering all data points. **(f) Relationship between copy number and expression for examples with outlier changes in effect size in WT vs. KD conditions.** Outliers were identified by inspection of panel **e**. Schematics above each plot illustrate the position of the region that was studied in the MPRA (top; orange=STR, grey=flanking, the minP and GFP are annotated as in Fig. 1) and dark blue rectangles (bottom) show gene annotations at the corresponding locus. Each data point represents a single barcode/replicate pair. Green=WT, red=RNase H1 KD. Shaded regions indicate 95% confidence intervals.

G4 structures in promoters have been linked to formation of R-loops (reviewed in^41^), which consist of a three-stranded DNA-RNA hybrid structure formed during transcription. If these structures are not resolved, transcription may be stalled^42^. Thus, we hypothesized that G4-forming STRs would be particularly sensitive to impaired R-loop resolution. To test this, we performed MPRA experiments under both wild-type conditions and RNase H1 knockdown via CRISPRi^43^ in HeLa S3 cells (**Methods**). Building on evidence that RNase H1 resolves R loops, RNase H1 depletion is expected to stabilize these structures^44^ and therefore should result in reduced expression. This is particularly expected for variants with the potential to form R loops, including those predicted to form G4s.

Analysis of normalized expression (RNA/DNA ratios) across all variants revealed that CCCCG/CGGGG repeats, the most prevalent repeat unit in our data predicted to form G4, showed the most pronounced reduction in reporter expression relative to other repeat units. On the other hand, CCG/CGG repeats, which have the same GC content but are not predicted to form G4s, showed only a modest overall reduction in expression (**Fig. 6b**). Further analysis of CCCCG/CGGGG variants stratified by copy number showed increasing sensitivity to RNase H1 knockdown up to 4 repeats, after which the effect plateaus (**Fig. 6c**). This aligns with the predicted G4 score and is consistent with the known G4 consensus sequence (G_3–5_N_1–7_G_3–5_N_1–7_G_3–5_N_1–7_G_3–5_) which requires a minimum of 4 guanine tracts to form the quadruplex structure (**Fig. 6d**). Additional guanine tracts increase the predicted G4 score but in our system has no additional impact on expression beyond formation of a single G4. In contrast, other repeat classes showed relatively flat expression ratios independent of repeat length (**Fig. 6c**; **Supplementary Fig. 19**).

To assess how RNase H1 knockdown affects the relationship between repeat copy number and expression at individual loci, we compared effect sizes under WT vs. knockdown conditions for each repeat unit/locus pair. Effect sizes showed strong overall correlation between conditions, but with a global reduction in the knockdown (Pearson’s *r*=0.47; **Fig. 6e**). Among repeat units for which at least 100 repeat unit/locus pairs were included in the analysis, CCCCG/CGGGG showed the lowest correlation, indicating the greatest disruption of length-dependent effects under RNase H1 knockdown (**Fig. 6e**; **Supplementary Fig. 20**). On the other hand, CCG/CGG repeats showed similar correlation compared to non-G4 forming repeats. Inspection of repeat unit/locus pairs with the strongest reduction in effect sizes revealed that most consisted of CCCCG/CGGGG repeats (**Fig. 6e**). The CCCCG/CGGGG outliers with the most extreme reduction originated from promoter regions for three specific genes (*GSTM4*, *AGTRAP*, and *RNF38*; **Fig. 6e-f**). Importantly, all outlier loci examined contained native TSSs of the original nearby genes within the reporter construct. Analysis of both orientations of these outlier regions (forward and reverse strand G4 formation) showed that regardless of which strand was predicted to have the G4 structure, these loci remained similarly sensitive to RNase H1 knockdown (**Supplementary Fig. 21**).

## Discussion

We present here the first adaptation, to our knowledge, of massively parallel reporter assays (MPRAs) to systematically study the involvement of STRs in regulation of expression. We identified optimal conditions for performing MPRA for STRs, and built on previous observations by us and others showing that STRs with impacts on expression (eSTRs) are most strongly enriched in promoter regions^3,10,11,45,46^. We used MPRAs to perform deep perturbations for a subset of STRs that enabled us to demonstrate that STRs with the same or similar repeat units show largely consistent length-dependent regulatory effects, with additional, but more modest, impact from the composition of the flanking sequences, with consistent effects across different cell lines tested. Using a combination of computational secondary structure analysis and repeating our MPRA under RNAse H1 knockdown conditions demonstrated that associations between copy number and expression at G-quadruplex-forming CCCCG/CGGGG repeats are particularly sensitive to loss of RNase H1, providing support for an R-loop mediated mechanism for these repeats.

The development of our STR MPRA platform required extensive optimization to overcome the inherent challenges of working with repetitive sequences. STRs present unique technical difficulties for library construction due to their low-complexity nature, which can cause synthesis errors, cloning instabilities, sequencing artifacts, and analysis challenges. Through systematic optimization—from transitioning to high-fidelity oligonucleotide synthesis approaches to refining PCR and cloning procedures and implementing dual-platform NGS (Illumina and Element) technologies—we established robust experimental conditions for STR interrogation. This first successful adaptation of the MPRAs for comprehensive STR studies opens new avenues for systematic studies of the roles of STRs which are localized in other genomic regions such as enhancers.

Overall, we observed a strong relationship between GC content, of both the repeat unit and flanking regions, with expression. We found that variants with GC-rich repeat units tended to be more highly expressed (**Fig. 2d**), and that increasing repeat copy number at these loci resulted in increased expression. By contrast, AT-rich repeats tended to show lower overall expression and weaker, or in some cases negative, relationships between copy number and expression (**Fig. 3c**). For those repeats, our deep perturbation library demonstrated that differences across loci with the same unit could be partially explained by GC content of the flanking regions (**Fig. 5j**), again with higher flanking GC content leading to higher overall expression across the same repeats with the same copy number. This result aligns with previously observed increased GC content and CpG islands in active promoter regions^47,48^. It is also consistent with our previous observations from population-scale analysis in the Genotype-Tissue Expression (GTEx) cohort^11^, that GC-rich repeats tended to have strong positive effect sizes and were most strongly enriched near transcription start sites. Although we could not identify additional characteristics of flanking regions that influence the *trend* between repeat copy number and expression for particular repeat units, we note that flanking regions have a substantial impact on the *overall expression* of variants from different loci, suggesting the flanking region and STR contribute additively to the total observed expression for each variant (**Supplementary Fig. 13**). Further, while we expect that some STRs may act in a tissue-specific manner, effects we observed are largely consistent across the cell types studied here (HEK293T and HeLa S3), consistent with work by us and others showing that the majority of detected expression quantitative trait loci (including eSTRs) are shared across tissues^11,49^.

Despite the strong influence of GC content, we note that the observed STR effects here and in our previous study in humans cannot be explained by GC content alone. For example, AAGG and AC repeats have the same GC content (50%) but strikingly different patterns between repeat copy number and expression (**Supplementary Fig. 14**). Given the diverse repertoire of STRs (e.g. ranging from homopolymers to hexanucleotides and 0-100% GC content), and broad range of trends (e.g. increasing, decreasing, or non-linear) observed between copy number and expression across different repeat units, we hypothesize that the regulatory impacts of STRs are driven by multiple underlying mechanisms beyond GC content. One potential mechanism that has been previously proposed is the ability of STRs to form non-canonical DNA or RNA secondary structures that may influence transcription through multiple pathways, including effects on DNA accessibility^50^, polymerase processivity^51,52^, and recruitment of G4-binding proteins^41^. The correspondence between observed expression trends in the dpSTR library and computationally generated G4 predictions (**Supplementary Fig. 18**) provides support for a G4-mediated regulatory mechanism. Our observation that the most pronounced G4 effects occur at loci containing endogenous transcription start sites suggests biological relevance, where STR length polymorphism could directly impact transcriptional efficiency in native gene regulatory contexts. The length-dependent sensitivity of some STRs to RNase H1 knockdown, particularly of CGGGG/CCCCG repeats which have high predicted potential to form G4s, further supports this suggested mechanism. Knockdown of RNase H1 is expected to hinder R-loop resolution, and we thus hypothesize that longer G4-forming repeats may increasingly interfere with transcriptional processes when positioned near active transcriptional elements if the R-loop is not resolved.

Although many repeats showed linear trends between copy number and expression in hSTR1-2, expanding the lengths of copy numbers tested in our dpSTR library revealed multiple striking non-linear patterns. The most pronounced non-linear trend observed was for AAAC/TTTG repeats, which show an increasing trend between copy number and expression up to a peak at 3-4 repeat copies, which then decreases at longer copy numbers. Multiple recent studies have demonstrated that these repeats can form binding sites for *FOX*-related transcription factors (TFs). For example, recent structural work by Zhang et al.^27^ demonstrated that *FOXP3* binds most strongly to TTTG repeats compared to other T_n_G repeats, with binding affinity increasing as repeat number increases *in vitro*. Their crystallographic analysis revealed that *FOXP3* can form extensive multimeric complexes on longer repeat sequences, providing a potential mechanistic framework for STR-mediated gene regulation. Our observation of a peak in expression at 3-4 copies is in line with the recent discovery of a TTTG repeat in a primate-specific enhancer implicated in thyrotropin resistance^8^. While their study establishes four repeats as the non-pathogenic length associated with enhancer function, our data indicate that maximal reporter expression can be reached around three or four copies of the AAAC repeat, with flanking sequence context influencing which repeat number yields peak activity. This suggests that the precise copy number threshold for optimal transcription activation may be flexible and modulated by the surrounding genomic environment.

Beyond the binding of Fox-like TFs to AAAC/TTTG repeats, additional repeat types have been shown to impact TF binding and could potentially explain some of the signals observed in our MPRA. For example, AAGG repeats, which have been shown to be bound by the fusion protein EWS-FLI1 in Ewing Sarcoma^26^. Other repeats in our library have sequences that are similar to known consensus motifs for other TFs, such as CGGGGG repeats which could form binding sites for *SP1* (motif 5’-(G/T)GGGCGG(G/A)(G/A)(C/T)-3’^53^). Intriguingly, these or similar repeat units (AAAC, AAGG, and CGGGG) all show non-linear trends (**Supplementary Fig. 14**) suggesting certain optimal copy numbers could provide the ideal spacing and binding architecture for productive protein-DNA interactions for different repeat unit/TF pairs. In addition to forming binding sites themselves, Horton and colleagues demonstrated that STRs can dramatically alter the affinity of nearby TF binding sites^28^. Overall, our findings, in combination with these previous observations, suggest the impact of STRs on TF binding as a strong candidate mechanism driving the detected expression effects for a subset of STRs we studied here that should be investigated in future studies.

Our study has several limitations that should guide future research directions. First, our analysis focused on promoter-associated STRs, representing only a subset of the repetitive elements that may influence gene expression. Expansion to include enhancer or other STRs will be essential for comprehensive understanding of STR regulatory biology. Second, our episomal MPRA system with short (up to 152bp) flanking regions, while enabling high-throughput analysis, may not fully recapitulate the chromatin context and three-dimensional regulatory interactions present in native genomic settings. The increasing adoption of lentiviral MPRA approaches^54^ that integrate reporter constructs into chromatin may provide more biologically relevant insights into STR function. Further, as synthesis techniques improve it will be possible to consider additional flanking context^55^. Additionally, our current platform does not directly assess transcription factor binding or chromatin accessibility changes, limiting our ability to definitively establish the molecular mechanisms underlying observed regulatory effects.

Effect sizes in our MPRA did not show strong correlation with eSTR effect sizes observed in human studies (**Supplementary Fig. 9**). We suspect this is due to a variety of factors. First, the ranges of repeat lengths tested at most loci differed across MPRA vs. eSTR studies. Whereas we tested a wide length range in our hSTR libraries (-5, 0, and +5 repeat copies compared to hg38), the range of repeat lengths in humans is often much narrower and does not necessarily overlap with this range. Second, many of the GC-rich repeats with strongest signals in humans dropped out of our analysis due to challenges in synthesizing, cloning, and sequencing of long regions with 100% GC content (**Supplementary Fig. 3**). Third, eSTR studies were in different cell types (mostly tissues for GTEx eSTRs^11^, LCLs for GEUVADIS eSTRs^36^), compared to the HEK293T and HeLa S3 cell lines (and episomal context) used here. Finally, direct comparison of effect sizes from MPRA vs. eSTR data is challenging, since we observed that a single eSTR can be associated with expression of multiple different nearby genes with different effect sizes or directions, making the comparison of effect sizes across platforms ambiguous. In future studies we aim to design additional MPRA experiments explicitly capturing variants observed in the human population.

Overall, the observations in our study align with the broader conceptualization of STRs as evolutionary “tuning knobs”^3,5,46,56^ that can provide fine-scale regulation of gene expression or other molecular phenotypes via stepwise changes in the number of repeat units. The mutational properties of STRs—namely their tendency to expand and contract at high rates^57^—creates a unique evolutionary substrate where changes in repeat number provide extensive variation with minimal genetic load and the potential for rapid and reversible adjustment of complex traits.

In summary, our work establishes STR MPRA as a powerful platform for systematic investigation of the impact of repeat elements on regulatory activity. Our findings, including repeat and flanking sequence composition effects, non-linear effects with optimal repeat lengths, and the impact of STR-mediated DNA or RNA secondary structure—establish STRs as a critical component of the non-coding regulatory grammar. Future studies expanding our framework to additional STR classes and cellular contexts will further refine our understanding of how repetitive elements shape gene expression programs and contribute to phenotypic diversity.

## Methods

### STR selection and oligonucleotides design

All STRs within 3.6kb of transcription start sites (TSSs) of human genes were identified. TSSs for protein coding autosomal genes were obtained from Ensembl version 104 (hg38)^58^ using the BiomaRt R package^59^. For genes with multiple transcripts, transcripts were prioritized by transcript support level (highest support preferred) and then by length (longest transcript preferred) to identify a total of 18,762 unique transcripts and their TSSs. STR coordinates were obtained from the hg38 HipSTR^60^ reference. STRs on non-autosomal chromosomes, with complex sequences consisting of more than a single repeat unit, and with a reference length >60bp for non-homopolymers or >30bp for homopolymers were filtered. This left a total of 30,516 unique STRs within 3.6kb of a TSS. For each STR locus, 3 variants were constructed, corresponding to the reference (hg38) repeat copy number, reference -5 repeat copies, and reference +5 repeat copies. For a random subset of 8,452 loci, a 4th variant corresponding to the reference +3 repeat copies was included, to reach a total of 100,000 unique variants. All repeat regions consisted of perfect copies of the repeat unit for the locus, even if sequence imperfections existed in the reference sequence. For each unique STR locus, identical flanking genomic context centered on the STR was included. The length of the context (flanking) was chosen according to the longest variant at each locus, computed as: *oligonucleotide length - length of required oligonucleotide elements - length of the largest repeat variant*. The maximum length of the variant plus context in each region was 168bp.

The final oligonucleotide consisted of the following elements: (1) 5’ adapter (5’-ACTGGCCGCTTGACG-3’), (2) the repeat + context sequence described above, (3) Gibson and AsiSI site (5’-CACTGCGGCTCCTGCGATCGC-3’), (4) a global filler sequence, (5) a BsaI recognition site (5’-GGTCTC-3’), and a Gibson and BsaI cut site (5’-TGTCGATCGCGTCGACGAAC-3’). The filler and other sequences are provided in **Supplementary Table 1**. The length of the filler sequence used was chosen such that the length of all variants for each locus had identical oligonucleotide lengths, computed as: *oligonucleotide length - length of elements 1,2,3 and 5 described above* for each variant. All final oligonucleotides containing more than one match to elements (1), (3), or (5) were filtered.

### MPRA experimental procedure

Detailed information of the molecular biology processes are provided in **Supplementary Experimental Procedures**. We provide here a brief overview with a focus on the modifications for adjusting the procedures for STRs.

#### Oligonucleotide synthesis, random barcoding and generation of the STR-barcode association libraries

Oligonucleotides of length 230bp were synthesized using Agilent Technologies HiFi Oligo Pools (100k total sequences for hSTR1/2 and 59,842 for dpSTR). Random barcodes of length 20bp were added to STR oligonucleotides by emulsion PCR (ePCR) following a modified version of the procedure from Tewhey et. al.^34^. The amplified oligonucleotides were cloned into the pGL4.23 backbone plasmid using NEBuilder HiFi DNA Assembly (E2621S). Each 100 µL ligation reaction contained 0.135 µg of barcoded-oligonucleotides and 0.54 µg of the linearized plasmid. For the hSTR1 and dpSTR libraries, 4 µL of the ligated vector was transformed into 50µL of NEB10-beta competent cells (NEB C3019H) using heat shock transformation. For hSTR2, 1.5 µL of the purified ligated vector was electroporated into 50 µL of 10-beta *E. coli* (NEB, C3020K), using the Gene Pulser Xcell Microbial System (Bio-rad, 1652662). Each vial was recovered in 2 mL of NEB10-beta out-growth medium at 37°C for 1 hour; the recovered cells were then combined. The plasmid library was purified using the NucleoBond Xtra Midi kit (Item number: 740412.50) and validated by Plasmidsaurus whole plasmid sequencing. Libraries for NGS analysis of the STR-barcode association were prepared using emulsion PCR (ePCR). The products were sequenced by Illumina NextSeq 500/550 (High Output Kit v2.5 150-cycle, Catalog ID 20024907) or Element Aviti (2x75 Sequencing Kit Cloudbreak FS High Output and Element Bioscience 860-00015, 184 cycles and 1 billion reads) with two indices of 8bp, a read 1 length of 25bp and a read 2 length of 135bp to obtain barcode/oligo pairings (“*Sequencing round 1*” in **Fig. 1**).

#### Generating the MPRA expression plasmids

The MPRA expression plasmids were generated by inserting a minimal promoter attached to GFP (minP-GFP) amplicon into linearized plasmids using the NEBuilder® HiFi DNA assembly reaction protocol. For the hSTR1 and dpSTR libraries, the resulting plasmid libraries were transformed by heat shock to NEB 10-beta competent *E. coli* cells in six parallel transformations. For the hSTR2 library, the resulting plasmid library was electroporated as above into NEB 10-beta electrocompetent *E. coli* in six parallel transformations. The plasmid libraries were QCed by Plasmidsaurus sequencing and digestion with restriction enzymes.

#### Cell culture, transfection, RNA/DNA extraction and cDNA synthesis and NGS for expression libraries

HEK293T cells and HeLa S3 cells transfections were conducted in three replicates using either PDL (Poly-D-Lysine) coated 10 cm dishes or PDL-coated T225 flasks with cells at about 80% confluence using Lipofectamine 3000. Each replicate used 0.22-0.25 μg of plasmids per cm^2^. At 40 hours post transfection, the cells were dissociated and collected in RLT buffer. Total RNA and DNA were extracted from cells using an AllPrep DNA/RNA mini Kit (Qiagen, 80204); cDNA was made following the protocol from Gordon et al.^54^. MPRA expression libraries (“*Sequencing round 2*” in **Fig. 1**) were prepared from cDNA (RNA) and DNA by performing two rounds of PCR. Three replicates of indexed cDNA (RNA) and DNA libraries were pooled and sequenced to a depth of about 6.25 billion reads (via a combination of NextSeq 2000 P3 50 Cycles (Illumina ref# 20046810), single read, 1.2 billion reads; NextSeq 2000 P4 50 Cycles (Illumina, ref# 2101842) single read, 1.8 billion reads and NovaSeq 6000, 3.25 billion reads by The UCSD IGM Genomics Center.

### Associating barcodes with STR variants in hSTR1 and hSTR2

We analyzed the STR-barcode association NGS libraries (“*Sequencing round 1”*), merging reads from Illumina and Element) to determine which variant was associated with each barcode. Reads were aligned to a reference built from our oligonucleotide sequences using BWA-MEM v0.7.18^61^ with option -M. We then implemented a stringent CIGAR string-based filtering approach for aligned read 1 (STR element) sequences. Specifically, we accepted sequences meeting any of the following criteria: (1) perfect matches to the reference, (2) sequences containing a single insertion or deletion smaller than 2bp, or (3) sequences with terminal soft-clipping less than 10bp in length. Additionally, all accepted sequences were required to contain fewer than 4 total CIGAR string elements. For sequences passing these filters, we extracted the corresponding barcode from read 2 to generate a preliminary barcode-variant association table.

In instances where a single barcode was associated with STR variants from two or more different loci, we removed that barcode since the assignment was ambiguous. For cases where a single barcode was associated with variants from the same locus, we applied a simplified model of PCR stutter based on that used in HipSTR^60^ to determine the most likely variant for each barcode. Our model consists of two parameters: *u* and *d*, which denote the probability that a PCR error introduces an insertion or deletion error, respectively compared to the true repeat copy number in each read. Unlike in the full HipSTR model, which additionally models the size of errors as being drawn from a geometric distribution, we did not directly model step size since these libraries are constrained to have a small number of copy numbers (-5, 0, +3, +5) and possible step sizes that are not represented well by that model.

Let 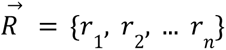 denote the vector of all reads observed for a given barcode, where *r_i_* denotes the repeat copy number of observed in read *i* (in this case, either -5, 0, +3, or +5). We define the likelihood of a particular repeat copy number *G* as 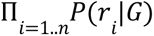 where *P*(*r_i_*|*G*) is equal to *u* if *r_i_* > *G*, *d* if *r_i_* < *G*, and 1 − *u* − *d* if *r_i_* == *G*. We applied an expectation-maximization approach based on that used in HipSTR to iteratively (1) infer maximum likelihood copy numbers for each barcode and (2) estimate *u* and *d* assuming the copy numbers from step 1 as ground truth, until convergence (each parameter changes by <0.001 compared to the previous iteration). Parameters *u* and *d* were learned separately for each repeat unit length (homopolymers, dinucleotides, trinucleotides, tetranucleotides, pentanucleotides, and hexanucleotides). After determining maximum likelihood genotypes using the final inferred values for *u* and *d*, we discarded remaining barcodes whose variant assignments were supported by less than 3 reads.

### Testing for association between repeat copy number and expression

We determined the expression level of each variant using the MPRA expression NGS libraries (“*Sequencing round 2”*). For each library type (RNA or DNA), we first extracted barcodes from each read and computed the total counts of each unique barcode. For each barcode, we quantified its expression as the ratio of the RNA/DNA counts. Using the previously generated association table, we mapped each barcode to its corresponding STR variant.

To assess the relationship between STR length and expression, we performed a separate linear regression analysis for each locus in which expression values for each barcode across all replicates were jointly analyzed in a single regression model: *Y_i_* = β*X_i_* + γ*R_i_*, where *Y_i_* is the RNA/DNA ratio of expression value *i*, *X_i_* is the length of the repeat relative to the hg38 reference for the variant associated with the barcode for expression value *i* (for hSTR1/2, *X_i_* ɛ {−5, −3, 0, +5}), *R* encodes the replicate (as two dummy variables) for expression value *i*, and ɛ is the error term. The regression coefficients β (STR effect size) and γ (covariate effect size) are learned. Regression models were fit using statsmodels v0.13.2. We required each analyzed locus to have data from a minimum of three distinct variants, with each variant represented by at least three unique barcodes to ensure robust measurements. To control for multiple hypothesis testing, we adjusted the resulting P-values for the β regression coefficient using the Benjamini-Hochberg false discovery rate (FDR) correction method^62^. We performed identical regression and filtering procedures separately for each locus in hSTR1 and hSTR2.

### Comparison of MPRA and eSTR effect sizes

We compared our MPRA effect sizes with our published population-based eSTR studies, which identified associations between average repeat copy number and expression of nearby genes in a cohort of individuals. eSTR summary statistics from Fotsing *et al.*^11^ computed based on 17 tissues from the Genotype Tissue Expression (GTEx) project cohort^49^ were obtained from Supplementary Dataset 1 of that study. eSTR summary statistics based on lymphoblastoid cell lines (LCLs) for individuals of European ancestry from the GEUVADIS cohort^63^ were obtained from Ziaei-Jam *et al.* ^36^ Supplementary Data 14.

### Designing the dpSTR (deep perturbation) array

We identified 200 loci and 100 loci with strong negative effects of repeat copy number on expression in the hSTR1/2 libraries as a basis for designing a second pool of 60k oligonucleotides (dpSTR). For each locus, we generated variant sequences with the following perturbations: (1) we varied the length of the STR region from 0 to *n* repeat copies, where the maximum copy number *n* depended on the length of the original repeat unit at the locus (*n*=23 for homopolymers, 14 for dinucleotides, 15 for tri- and tetra-nucleotides, 13 for pentanucleotides, and 11 for hexanucleotides). (2) For each copy number *c* ε {1.. *n*} considered, we varied the repeat unit to be either: the original repeat unit, and up to one alternative unit each from a set of candidate homopolymer (“A”), dinucleotide (“AC”, “AT’, “AG”, or “GC”), trinucleotide (“CCG”, “AAT”, “ACG”, or “AAG”), or tetranucleotide (“AAAC”, “AGAT”, “AAAT”, or “ACAT”) repeat units. For each of these, the variant region was replaced with *c* perfect copies of the repeat unit. We additionally included a set of variants with the original repeat unit but for which we did not remove repeat imperfections present in the reference genome sequence. Finally, we replaced the variant region with either random sequence, or random sequence with matched GC content to the original repeat unit. Note in order to enable space for a large number of perturbations in the 60k oligonucleotides we excluded random perturbations for which *c* mod 3!=0 (i.e. only lengths corresponding to 3, 6, 9, etc. copies of the original repeat unit were considered). We excluded any perturbations for which the total STR length exceeded 80bp. (3) We additionally included a sequence with the entire repeat unit removed (copy number 0). For each repeat length/repeat unit perturbation in 1-3, we included both the total variable region and its reverse complement sequence. Any variable regions containing the AsiSI or BsaI recognition sites were filtered. We removed duplicate sequences (e.g. if the original repeat was already perfect there in some cases will be two identical sequences). Finally, we manually added the exact variant sequences considered for the 300 candidate loci in the hSTR1/2 design to enable direct comparison to those libraries. For each sequence, we added the same elements (5’ adapter, variant sequence, Gibson and AsiSI site, filler, BsaI recognition site, and Gibson and BsaI cut site) as for hSTR1/2. The full set of oligonucleotides used is provided in **Supplementary Table 3.** Library construction, transfection, and generation of NGS libraries for identifying STR-barcode associations and for measuring expression from RNA and DNA barcode counts, were performed using the procedures described above.

### Analysis pipeline for the dpSTR array

The dpSTR library contains a large number of similar sequences compared to hSTR1/2 and thus required a new analysis pipeline to identify STR-barcode associations from NGS data (“*Sequencing round 1”*). We generated two separate references for alignment based on forward and reverse complement sequences of all variants. Reads from the STR-barcode association library were aligned separately to each reference using BWA-MEM v0.7.18^61^ with parameters -M -k 70 -t 8 -O 28 -L 10. The options -k (minimum seed length), -O (gap open penalty) and -L (clipping penalty) are set to stringent values to force precise and ungapped matches to the reference sequences. After alignment, we retained reads only if they were uniquely aligned with no insertions or deletions to the forward strand of a variant using one reference and the corresponding reverse complement sequence using the other reference, indicating concordant variant assignments using either reference.

In instances where a single barcode was associated with STR variants from two or more different variants from the same stranded locus/repeat unit pair, we applied a maximum likelihood estimation-based stutter error model. For the dpSTR library, which includes all possible repeat copy numbers in a certain range (compared to variants spaced by at least two units in length in hSTR1/2), we employed the full 3-parameter HipSTR stutter error model. For each read, in addition to modeling the probability to see an insertion (u) or deletion (d) error, we model the size of the error as being drawn from a geometric distribution with parameter ρ. Similar to hSTR1/2, we iteratively estimated maximum likelihood genotypes and inferred values of u, d, and ρ until convergence (each parameter changes by <0.001 compared to the previous iteration). This procedure was performed separately across all locus/repeat unit pairs for each repeat unit length to learn one set of parameters for homopolymers, dinucleotides, etc. In cases where a barcode was assigned to multiple stranded locus/repeat unit pair, if one pair accounted for more than 95% of the barcode counts and the total number of barcodes exceeded 100, the barcode was assigned to the dominant locus/repeat unit pair. Otherwise it was discarded. We additionally discarded remaining barcodes whose variant assignments were supported by less than 5 reads.

Similarly to hSTR1/2, we analyzed MPRA expression libraries (“*Sequencing round 2”*) by extracting barcodes from each read from each library type (RNA or DNA) and computing the total counts of each unique barcode. For each barcode, we quantified its expression as the ratio of the RNA/DNA counts. For downstream analyses, we excluded the top and bottom 5% expression outliers for each variant within each replicate. We required each analyzed locus to have data from a minimum of three distinct variants, with each variant represented by at least three unique barcodes to ensure robust measurements. Regression analyses to identify associations between copy number and expression were performed in the same manner as described for hSTR1/2. A separate regression analysis was performed for each locus/repeat unit/strand combination.

### Visualization of aggregate expression profiles

We generated visualizations (“meta-profiles”) of expression profiles in aggregate across different repeat units. To enable comparison across loci with different baseline expression levels, we first computed an aggregate measure of expression for each unique combination of locus, repeat unit, and strand orientation using the ratio of RNA to DNA counts summed across all barcodes. Expression values for each repeat length were normalized by this aggregate expression metric before visualization (e.g. **Fig. 5 b,d,h**). This process was performed separately for each replicate.

### Dimensionality reduction with UMAP

We performed dimensionality reduction analysis to visualize relationships among STR variants using the Uniform Manifold Approximation and Projection (UMAP) algorithm^64^. The input data consisted of a matrix of expression values for each repeat unit/locus pair (rows) and copy number (columns). Expression values were computed by averaging RNA/DNA values for each barcode across all replicates. These averaged expression values were then used as input features for UMAP analysis, implemented using the Python UMAP package with default parameters.

### Structural analysis for the deep perturbation array

Structural data predictions were performed on the STR region and flanking genomic sequences of the dpSTR oligonucleotide library. We used the mfold command-line tool from the UNAFold software package^65^ (v3.6) with default settings to predict RNA and DNA folding energies. Z-DNA forming potential was assessed using ZSeeker^66^ v1.0 with default thresholds, and G-quadruplex structures were predicted using the G4Hunter^67^ command-line tool (no version information available) with default parameters (window 25, threshold 1.2). Since ZSeeker and G4Hunter may output multiple scores per sequence, we considered both the mean and sum of the scores for each sequence. For each unique combination of locus, repeat unit type, and strand orientation, and structural prediction data, we retained both the raw scores and calculated the z-score across repeat lengths 1-15.

### Generating RNase H1 KD cells

RNase H1 KD HeLa S3 cells were generated using CRISPRi. A gRNA donor backbone vector, PB_gRNA_S.pyogenes_scaffold_BB (Addgene plasmid #226429) was generated using the plasmid PB-U6insert-EF1puro (Addgene Plasmid #104537). The PB-U6insert-EF1puro plasmid was digested with EcoRI-HF and AgeI-HF followed by ligation of annealed oligos from IDT containing a BbsI Golden Gate cloning site upstream of the *S. Pyogenes* gRNA scaffold sequence. The RNase H1 CRISPRi gRNA Donor Vector Library was generated by pooling together six different protospacer sequences targeting the RNase H1 promoter (selected from Human CRISPR Inhibition Pooled Library^68^). The HeLa S3 cell line expressing dCas9-tag-BFP-KRAB (gift from Dr. Komor’s lab at UCSD) was transfected with a human PiggyBac transposase expressing plasmid^69^ as well as the RNase H1 CRISPRi gRNA Donor Vector Library using Lipofectamine 3000 Transfection Reagent (L3000015), at a 1:2.5 molar ratio of transposase to transposon vector. Cells were selected with puromycin (Invitrogen cat. A1113803) for ∼5 days.

## Data availability

All raw and processed data relating to the MPRA experiment can be found on the Gene Expression Omnibus accession GSE306816.

## Code availability

Scripts for designing the original MPRA hSTR1/2 oligonucleotides are at https://github.com/gymreklab/str_mpra_design. Analysis scripts and scripts for designing the dpSTR oligonucleotides are at: https://github.com/gymreklab/str-mpra/.

## Supporting information

zhang-etal-SuppMaterial

zhang-etal-SupplementaryTables

## Author contributions

X.Z. performed and conceived computational analyses and wrote the manuscript. L.Z. supervised and performed MPRA experiments for all libraries. E.W. performed structure analysis and helped perform MPRA experiments and analysis. S.B. helped generate the hSTR1 and hSTR2 libraries and design the dpSTR array. Ed.M. generated the RNaseI KD cell line. M.M. designed the hSTR1/2 array. S.S-B. and C.W. performed initial MPRA experiments that guided design of the hSTR1/2 libraries. Q.G. designed initial computational analysis frameworks. M.L. performed computational analysis of pilot libraries. Er.M. helped supervise experiments and interpret analysis results. M.G. and A.G. conceived the project, obtained funding, wrote the manuscript, and co-supervised all components of this study.

## Acknowledgements

Research reported in this publication was supported in part by NIH/NHGRI grant R01HG010885 to M.G and A.G., Ed.M was supported in part by an institutional award to the UCSD Genetics Training Program from the National Institute for General Medical Sciences, T32 GM145427. This publication includes NGS data generated at the UC San Diego IGM Genomics Center utilizing an Illumina NovaSeq 6000 that was purchased with funding from a National Institutes of Health SIG grant (#S10 OD026929) as well as via sequencing services by the Stem Cell Genomics Core at the Sanford Stem Cell Institute. We thank J. Satovsky and T. Dishon for their assistance and input with Western blots and other experimental protocols, and R. Wachs for her help with the illustrations. We also thank the Applications team at Element for generating a subset of the sequencing data, R. Tewhey for advice on setting up MPRA experiments, and C. Benner for helpful feedback on the manuscript.

## References

1. Hannan, A. J. Tandem repeats mediating genetic plasticity in health and disease. Nat Rev Genet 19, 286–298 (2018).

2. Press, M. O., McCoy, R. C., Hall, A. N., Akey, J. M. & Queitsch, C. Massive variation of short tandem repeats with functional consequences across strains of Arabidopsis thaliana. Genome Res 28, 1169–1178 (2018).

3. Vinces, M. D., Legendre, M., Caldara, M., Hagihara, M. & Verstrepen, K. J. Unstable tandem repeats in promoters confer transcriptional evolvability. Science 324, 1213–1216 (2009).

4. Verbiest, M. et al. Mutation and selection processes regulating short tandem repeats give rise to genetic and phenotypic diversity across species. J Evol Biol 36, 321–336 (2023).

5. Mirkin, S. M. Expandable DNA repeats and human disease. Nature 447, 932–940 (2007).

6. Mitra, I. et al. Patterns of de novo tandem repeat mutations and their role in autism. Nature 589, 246–250 (2021).

7. Trost, B. et al. Genome-wide detection of tandem DNA repeats that are expanded in autism. Nature 586, 80–86 (2020).

8. Grasberger, H. et al. STR mutations on chromosome 15q cause thyrotropin resistance by activating a primate-specific enhancer of MIR7-2/MIR1179. Nat Genet 56, 877–888 (2024).

9. Grünewald, T. G. P. et al. Chimeric EWSR1-FLI1 regulates the Ewing sarcoma susceptibility gene EGR2 via a GGAA microsatellite. Nat Genet 47, 1073–1078 (2015).

10. Gymrek, M. et al. Abundant contribution of short tandem repeats to gene expression variation in humans. Nat Genet 48, 22–29 (2016).

11. Fotsing, S. F. et al. The impact of short tandem repeat variation on gene expression. Nat Genet 51, 1652–1659 (2019).

12. Tanudisastro, H. A. et al. Polymorphic tandem repeats influence cell type-specific gene expression across the human immune landscape. 2024.11.02.621562 Preprint at 10.1101/2024.11.02.621562 (2025).

13. Cui, Y. et al. Multi-omic quantitative trait loci link tandem repeat size variation to gene regulation in human brain. Nat Genet 57, 369–378 (2025).

14. Hamanaka, K. et al. Genome-wide identification of tandem repeats associated with splicing variation across 49 tissues in humans. Genome Res 33, 435–447 (2023).

15. Martin-Trujillo, A., Garg, P., Patel, N., Jadhav, B. & Sharp, A. J. Genome-wide evaluation of the effect of short tandem repeat variation on local DNA methylation. Genome Res 33, 184–196 (2023).

16. Margoliash, J. et al. Polymorphic short tandem repeats make widespread contributions to blood and serum traits. Cell Genom 3, 100458 (2023).

17. Manigbas, C. A. et al. A phenome-wide association study of tandem repeat variation in 168,554 individuals from the UK Biobank. Nat Commun 15, 10521 (2024).

18. Jadhav, B. et al. A phenome-wide association study of methylated GC-rich repeats identifies a GCC repeat expansion in AFF3 associated with intellectual disability. Nat Genet 56, 2322–2332 (2024).

19. Conlon, E. G. et al. The C9ORF72 GGGGCC expansion forms RNA G-quadruplex inclusions and sequesters hnRNP H to disrupt splicing in ALS brains. Elife 5, e17820 (2016).

20. Lin, Y., Dent, S. Y. R., Wilson, J. H., Wells, R. D. & Napierala, M. R loops stimulate genetic instability of CTG.CAG repeats. Proc Natl Acad Sci U S A 107, 692–697 (2010).

21. Rothenburg, S., Koch-Nolte, F., Rich, A. & Haag, F. A polymorphic dinucleotide repeat in the rat nucleolin gene forms Z-DNA and inhibits promoter activity. Proc Natl Acad Sci U S A 98, 8985–8990 (2001).

22. Liu, X. S. et al. Rescue of Fragile X Syndrome Neurons by DNA Methylation Editing of the FMR1 Gene. Cell 172, 979–992.e6 (2018).

23. Raveh-Sadka, T. et al. Manipulating nucleosome disfavoring sequences allows fine-tune regulation of gene expression in yeast. Nat Genet 44, 743–750 (2012).

24. Suter, B., Schnappauf, G. & Thoma, F. Poly(dA.dT) sequences exist as rigid DNA structures in nucleosome-free yeast promoters in vivo. Nucleic Acids Res 28, 4083–4089 (2000).

25. Sela, I. & Lukatsky, D. B. DNA Sequence Correlations Shape Nonspecific Transcription Factor-DNA Binding Affinity. Biophys J 101, 160–166 (2011).

26. Gangwal, K. et al. Microsatellites as EWS/FLI response elements in Ewing’s sarcoma. Proc Natl Acad Sci U S A 105, 10149–10154 (2008).

27. Zhang, W. et al. FOXP3 recognizes microsatellites and bridges DNA through multimerization. Nature 624, 433–441 (2023).

28. Horton, C. A. et al. Short tandem repeats bind transcription factors to tune eukaryotic gene expression. Science 381, eadd1250 (2023).

29. Heydari, M., Miclotte, G., Van de Peer, Y. & Fostier, J. Illumina error correction near highly repetitive DNA regions improves de novo genome assembly. BMC Bioinformatics 20, 298 (2019).

30. Tanudisastro, H. A., Deveson, I. W., Dashnow, H. & MacArthur, D. G. Sequencing and characterizing short tandem repeats in the human genome. Nat Rev Genet 25, 460–475 (2024).

31. Agarwal, V. et al. Massively parallel characterization of transcriptional regulatory elements. Nature 639, 411–420 (2025).

32. Melnikov, A. et al. Systematic dissection and optimization of inducible enhancers in human cells using a massively parallel reporter assay. Nat Biotechnol 30, 271–277 (2012).

33. Lee, S. et al. Massively parallel reporter assay investigates shared genetic variants of eight psychiatric disorders. Cell 188, 1409–1424.e21 (2025).

34. Tewhey, R. et al. Direct Identification of Hundreds of Expression-Modulating Variants using a Multiplexed Reporter Assay. Cell 165, 1519–1529 (2016).

35. Arslan, S. et al. Sequencing by avidity enables high accuracy with low reagent consumption. Nat Biotechnol 42, 132–138 (2024).

36. Ziaei Jam, H., et al. A deep population reference panel of tandem repeat variation. Nat Commun 14, 6711 (2023).

37. Klein, J. C. et al. A systematic evaluation of the design and context dependencies of massively parallel reporter assays. Nat Methods 17, 1083–1091 (2020).

38. Narumi, S. et al. Functional variants in a TTTG microsatellite on 15q26.1 cause familial nonautoimmune thyroid abnormalities. Nat Genet 56, 869–876 (2024).

39. Murat, P., Guilbaud, G. & Sale, J. E. DNA polymerase stalling at structured DNA constrains the expansion of short tandem repeats. Genome Biol 21, 209 (2020).

40. Esnault, C. et al. G-quadruplexes are promoter elements controlling nucleosome exclusion and RNA polymerase II pausing. Nat Genet 57, 1981–1993 (2025).

41. Wulfridge, P. & Sarma, K. Intertwining roles of R-loops and G-quadruplexes in DNA repair, transcription and genome organization. Nat Cell Biol 26, 1025–1036 (2024).

42. Tous, C. & Aguilera, A. Impairment of transcription elongation by R-loops in vitro. Biochem Biophys Res Commun 360, 428–432 (2007).

43. Qi, L. S. et al. Repurposing CRISPR as an RNA-guided platform for sequence-specific control of gene expression. Cell 152, 1173–1183 (2013).

44. Parajuli, S. et al. Human ribonuclease H1 resolves R-loops and thereby enables progression of the DNA replication fork. J Biol Chem 292, 15216–15224 (2017).

45. Quilez, J. et al. Polymorphic tandem repeats within gene promoters act as modifiers of gene expression and DNA methylation in humans. Nucleic Acids Res 44, 3750–3762 (2016).

46. Sawaya, S. M., Bagshaw, A. T., Buschiazzo, E. & Gemmell, N. J. Promoter microsatellites as modulators of human gene expression. Adv Exp Med Biol 769, 41–54 (2012).

47. Deaton, A. M. & Bird, A. CpG islands and the regulation of transcription. Genes Dev 25, 1010–1022 (2011).

48. Saxonov, S., Berg, P. & Brutlag, D. L. A genome-wide analysis of CpG dinucleotides in the human genome distinguishes two distinct classes of promoters. Proc Natl Acad Sci U S A 103, 1412–1417 (2006).

49. The GTEx Consortium. The GTEx Consortium atlas of genetic regulatory effects across human tissues. Science 369, 1318–1330 (2020).

50. Puget, N., Miller, K. M. & Legube, G. Non-canonical DNA/RNA structures during Transcription-Coupled Double-Strand Break Repair: Roadblocks or Bona fide repair intermediates? DNA Repair (Amst) 81, 102661 (2019).

51. Zhang, X. et al. Cockayne Syndrome Linked to Elevated R-Loops Induced by Stalled RNA Polymerase II during Transcription Elongation. Nat Commun 15, 6031 (2024).

52. Tateishi-Karimata, H. & Sugimoto, N. Roles of non-canonical structures of nucleic acids in cancer and neurodegenerative diseases. Nucleic Acids Res 49, 7839–7855 (2021).

53. Raiber, E.-A., Kranaster, R., Lam, E., Nikan, M. & Balasubramanian, S. A non-canonical DNA structure is a binding motif for the transcription factor SP1 in vitro. Nucleic Acids Res 40, 1499–1508 (2012).

54. Gordon, M. G. et al. lentiMPRA and MPRAflow for high-throughput functional characterization of gene regulatory elements. Nat Protoc 15, 2387–2412 (2020).

55. Arnould, C. et al. Capture-C MPRA: A high-throughput method to simultaneously characterize promoter interactions and regulatory activity. 2025.06.11.658967 Preprint at 10.1101/2025.06.11.658967 (2025).

56. Gemayel, R., Vinces, M. D., Legendre, M. & Verstrepen, K. J. Variable tandem repeats accelerate evolution of coding and regulatory sequences. Annu Rev Genet 44, 445–477 (2010).

57. Ellegren, H. Microsatellite mutations in the germline: implications for evolutionary inference. Trends Genet 16, 551–558 (2000).

58. Howe, K. L. et al. Ensembl 2021. Nucleic Acids Res 49, D884–D891 (2021).

59. Durinck, S., Spellman, P. T., Birney, E. & Huber, W. Mapping identifiers for the integration of genomic datasets with the R/Bioconductor package biomaRt. Nat Protoc 4, 1184–1191 (2009).

60. Willems, T. et al. Genome-wide profiling of heritable and de novo STR variations. Nat Methods 14, 590–592 (2017).

61. Li, H. Aligning sequence reads, clone sequences and assembly contigs with BWA-MEM. Preprint at 10.48550/arXiv.1303.3997 (2013).

62. Benjamini, Y. & Hochberg, Y. Controlling the False Discovery Rate: A Practical and Powerful Approach to Multiple Testing. Journal of the Royal Statistical Society. Series B (Methodological) 57, 289–300 (1995).

63. Lappalainen, T. et al. Transcriptome and genome sequencing uncovers functional variation in humans. Nature 501, 506–511 (2013).

64. McInnes, L., Healy, J. & Melville, J. UMAP: Uniform Manifold Approximation and Projection for Dimension Reduction. Preprint at 10.48550/arXiv.1802.03426 (2020).

65. Markham, N. R. & Zuker, M. UNAFold: software for nucleic acid folding and hybridization. Methods Mol Biol 453, 3–31 (2008).

66. Wang, G. et al. ZSeeker: an optimized algorithm for Z-DNA detection in genomic sequences. Brief Bioinform 26, bbaf240 (2025).

67. Bedrat, A., Lacroix, L. & Mergny, J.-L. Re-evaluation of G-quadruplex propensity with G4Hunter. Nucleic Acids Res 44, 1746–1759 (2016).

68. Sanson, K. R. et al. Optimized libraries for CRISPR-Cas9 genetic screens with multiple modalities. Nat Commun 9, 5416 (2018).

69. Farah, E. N. et al. Spatially organized cellular communities form the developing human heart. Nature 627, 854–864 (2024).

