## Supplementary material for "Systematic evaluation of the impact of promoter proximal short tandem repeats on expression": zhang-etal-SuppMaterial

Supplementary Material for Zhang *et al.* "Systematic evaluation of the impact of promoter proximal short tandem repeats on expression"

#### Contents:

Supplementary Experimental Procedures
Supplementary Figures 1-21
Supplementary Table 1-3 Legends
Supplementary References

#### Supplementary Experimental Procedures

##### *MPRA experimental procedure*

Oligonucleotide synthesis: Oligonucleotides of length 230nt were synthesized using Agilent Technologies SurePrint HiFi Oligo Pools (100k total sequences for hSTR1/2 and 60k for the dpSTR libraries).

Random barcoding: Random 20nt barcodes were added to STR oligonucleotides by emulsion PCR (ePCR) following a modified version of the procedure from Tewhey et. al[1]. Oligonucleotides (10 pmol) were dissolved in a 20 μL IDTE buffer (pH 8.0, IDT) to obtain a 500 nM solution. Random 20nt barcodes and Gibson homology sequences were added via 5 (hSTR1 and dpSTR) or 28 (hSTR2) separate reactions of ePCR. Each 50 μL reaction contained 0.2 ng (hSTR1 and dpSTR) or 1.86 ng (hSTR2) of the oligonucleotides, 1X PrimeSTAR Max Premix (PrimeSTAR® Max DNA Polymerase, R045A), 0.5 μM MPRA_v3_Fwd (5'-GCCAGAACATTTCTCTGGCCTAACTGGCCGCTTGACG-3') and MPRA_v2_20I_Rev primers (5'-AGCGGCCGGCCGCCCCGACTACGCTCTTCCGATCTNNNNNNNNNNNNNNNNNNNNTCTAGAGGTTCGTCGACGCGATC-3'), and 2 ng of BSA (NEB, B9000).

For ePCR reaction: PCR master mix was emulsified by vortexing with a 220 μL emulsion element 1, 20 μL emulsion element 2, and 60 μL emulsion element 3 (EURx, E3600) per 50 μL of PCR mix at 4ºC for 5 minutes at 3000 RPM. The emulsion mixture was plated (100 μL per well) across a 96 well plate with the following conditions: 95°C for 30 sec, 15 cycles of (95°C for 30 sec, 58°C for 10 sec, 72°C for 15 sec), 72°C for 5 min, hold at 4ºC. The amplified emulsion mixture was broken by adding 1 mL of isobutyl alcohol (ThermoFisher Scientific, A397-1) and purified (either by following Tewhey et al.[1] or Verma et al[2] with some modifications). In particular, the samples were purified either by Monarch PCR Purification Kit (NEB, Cat# T1030S) or by SpeedBeads (adding 130 µL of a *SpeedBeads master mix* (2 µL of SpeedBeads (Cytiva 65152105050250), 36 µL of 40% PEG 8000 (Sigma-Aldrich, p1458-50ml), 40 µL of 5M NaCl, and 52 µL of UltraPure Distilled Water (Invitrogen,10977-015)) per 350 µL of emulsion mix. The mixture was vortexed vigorously, then incubated for 10 minutes at room temperature. The broken emulsion/isobutyl alcohol mixture was spun at 2900 RCF for 5 min and the isobutyl alcohol phase was discarded. The aqueous phase was placed on a magnetic rack for 20 minutes prior to aspiration. The remaining beads were washed once with isobutyl alcohol, then three times with 80% ethanol and eluted in IDTE buffer (Integrated DNA Technologies; IDT). The resulting product was run on 1.5% agarose gel to check the PCR product size of 314bp.

Cloning the MPRA library to the pGL4.23 backbone plasmid: The linearized plasmid backbone was generated by digesting pGL4.23[luc/minP] (Promega E8411) with XbaI (NEB, Cat# R0145L) and incubating for 5 hours at 37°C, then adding SfiI (NEB, R0123L) and incubating overnight at 50°C. The product was then run on a 1% agarose gel and the correct band (2493bp) was cut and DNA was extracted by using the Zymoclean Gel DNA recovery kit (Cat# 11-301). The barcoded oligonucleotides (barcoded-oligos) were cloned using NEBuilder HiFi DNA Assembly (E2621S) Each 100 µL reaction contained 0.135 µg of barcoded-oligos and 0.54 µg of the linearized plasmid, and incubated for 60 min at 50ºC.

For the hSTR1 and dpSTR libraries, 4 µL of the ligated vector was transformed into 50 µL of NEB10-beta competent *E. coli* cells (NEB, C3019H) using heat shock transformation. For hSTR2, 1.5 µL of the purified ligated vector was electroporated into 50 µL of 10-beta electrocompetent *E. coli* (NEB, C3020K), by using the Gene Pulser Xcell Microbial System (Bio-rad, 1652662) following the company's electroporation protocol (2 kV, 200 ohm, 25 µF) in six parallel transformations. Each vial was independently recovered in 2 mL of NEB10-beta out-growth medium at 37°C in 12 mL bacterial culture tubes, shaking at 225 rpm for 1 hour; the recovered cells were then combined.

To determine CFU, 50 µL of the recovered cells were used for serial dilutions (1:10, 1:100, 1:1000 and 1:10,000). 100 µL of each dilution was then plated onto an LB Ampicillin Agar plate, grown overnight, and counted. We estimated CFUs of 10 million for hSTR1 and 100 million for hSTR2. The remaining recovered cells were divided into 3 flasks, with 4 mL of recovered cells added to 400 mL LB and Carbenicillin 100 ug/ml. These flasks were shaken at 225 rpm at 30ºC for 13h-14h until the checked OD600 was 0.9 to 1. Based on CFU count, all flasks were mixed then spun down by centrifuging at 6000-6500 RPM for 20 min at 4ºC. After removal of the liquid, the bacterial cell pellet was weighed. The plasmids were purified using the NucleoBond Xtra Midi kit (Item number: 740412.50) and validated by Plasmidsaurus (**URLs**) whole plasmid sequencing.

Generation of STR-barcode association libraries: Libraries for NGS analysis of the STR-barcode association were prepared from the plasmids containing these libraries (termed pGL4.23-hSTR1-BC; pGL4.23-dpSTR-BC and pGL4.23-hSTR2-BC) by performing two rounds of ePCRs. The same ePCR protocol as above was followed, using different primers and PCR conditions. The first round ePCR included four ePCR reactions, each 50 µL (50 ng of the plasmid STR-barcode libraries), 25 µL of PrimeSTAR Max Premix (2X) (PrimeSTAR Max DNA Polymerase from Clontech, R045A) 0.5µM MPRA_TruSeq_Universal_Adapter_Fwd (5'-GTGACTGGAGTTCAGACGTGTGCTCTTCCGATCTACTGGCCGCTTGACG-3') and 0.5µM MPRA_TruSeq_Universal_Adapter_Rev (5'-ACACTCTTTCCCTACACGACGCTCTTCCGATCT-3'). To conduct the ePCR, the PCR master mix was emulsified by vortexing with 220 µL Emulsion element 1, 20 µL Emulsion element 2 and 60 µL Emulsion element 3 per 50 µL PCR reaction at 4°C for 5 min at 3000 RPM. 100 µL of emulsion mixture was plated per well in 8 strip PCR tubes and cycled with the following conditions: 95°C for 2 min, 6 cycles of (95°C for 20 sec, 58°C for 20 sec, 72°C for 30 sec), 72°C for 5 min, hold at 4ºC. The amplified emulsion mixture was broken and purified by using Monarch PCR Purification Kit (NEB, Cat# T1030S) (hSTR1 and dpSTR) or purified by SpeedBeads (hSTR2) as above. Purified libraries were eluted in 30 µL of ultrapure water. Sequencing indexes were added by running a second ePCR which included 4 reactions, using the product from the first PCR, and the same reaction conditions as above for 8 cycles. The reaction was purified by gel extraction (hSTR1 and dpSTR) or by both Speedbeads purification and gel extraction (hSTR2). The products were sequenced by Illumina NextSeq 500/550 or Element Aviti to obtain barcode/oligo pairings.

Generating the MPRA expression plasmids: The MPRA expression plasmids for all three libraries (hSTR1, hSTR2, and dpSTR) were generated by inserting a minimal promoter attached to GFP (minP-GFP) amplicon into linearized plasmids (pGL4.23 plasmids with barcoded STR oligos) for each library. minP-GFP was ordered as a gBlock from IDT, amplified by PCR using the MPRA_GFP_Fwd_v1

(5’-CACTGCGGCTCCTGCGATCTAACTGGCCGGTACCTGAGCTCGCTA-3’) and MPRA_GFP_Rev_v1 (5’-TCTAGAGGTTCGTCGACGCGATTATTATCATTACTTGTACAGCTCGTCCATGC-3’) according to Abell et al.[3] and purified by 7% PEG SpeedBeads. The product was QCed by Nanodrop, agarose gel and Plasmidsaurus sequencing.

The plasmids were linearized by removal of the “filler” sequence via digestion with 100 units of AsiSI (NEB, R0630S) and 100 units of BsaI-HFv2 (NEB, R3733L) in 1x rCutSmart Buffer (NEB) in a 400 µL volume for 16 hours at 37°C. The reaction was then run on a 1% agarose gel and larger bands were purified by Zymoclean Gel DNA recovery kit (Cat# 11-301) and quantified by Nanodrop.

The minP-GFP amplicon was cloned to the plasmids by following the NEBuilder HiFi DNA assembly reaction protocol (NEB E2621s), with molar ratio of 1:2 (vector (118ng):insert (78ng)) in 20 µL of reaction volume for 60 minutes at 50°C. Insertion of the minP-GFP amplicon to the hSTR2 library, was conducted by following Abell’s Gibson assembly (NEB, E2621s) protocol, using 10 µg of the minP-GFP amplicon and 3.3 µg of the linearized plasmid in 300 µL of total volume for 90 minutes at 50°C, followed by a 6.45% PEG SpeedBeads purification. For the hSTR1 and dpSTR libraries, the resulting plasmid libraries were transformed by heat shock to NEB10-beta competent *E. coli* cells (NEB C3019H) in six parallel transformations. For the hSTR2 library, the resulting plasmid library was electroporated into 10-beta electrocompetent *E. coli* in six parallel transformations using a BioRad Gene Pulser as before except using 2.5 L LB media with 100 µg/mL carbenicillin for 12-13 hours, OD600=0.9-1. After cultures were pooled, plasmids were purified by the NucleoBond Xtra Midi kit (Item number: 740412.50) according to standard protocols, resuspended in EB and quantified by Nanodrop. The plasmid libraries were QCed by Plasmidsaurus sequencing and digested by restriction enzymes (XbaI, 3577bp; or HindIII with XbaI, 2755bp and 822bp; or NcoI and XbaI, 2829bp and 748bp; or a combination of the three enzymes, 2755bp, 748bp and 74bp) to verify the sizes of the fragments.

<u>Cell culture and transfection:</u> HEK293T cells were grown in DMEM, high glucose, GlutaMAX Supplement (ThermoFisher 10566024) supplemented with 10% FBS. HeLa S3 cells and the HeLa S3 RNase H1 KD stable cell lines were grown in DMEM, high glucose, pyruvate (ThermoFisher 11995073) supplemented with 10% FBS and 2 mM L-glutamine. Transfections were conducted in three replicates using either PDL (Poly-D-Lysine) coated 10 cm dishes or PDL-coated T225 flasks with cells at about 80% confluence. The MPRA expression plasmids

described above were transfected to the cells using Lipofectamine 3000 following the manufacturer's protocol. Each dish or flask of cells was transfected with 0.22-0.25 μg of plasmids per $cm^2$, each 1 μg of plasmid was mixed with 2.5 μL of Lipofectamine 3000 and 2 μL of p3000 (Life Technologies, L3000015) in 50 μL of Opti-MEM Reduced Serum Medium. Cells were incubated with transfection reagents for 16-18 hours prior to changing fresh medium. 40 to 42 hours post transfection, the cells on 10 cm dishes were washed with PBS followed by adding RLT buffer supplemented with beta-mercaptoethanol (beta-ME), to lyse the cells; the cells on the flask were washed with PBS followed by dissociation with 0.05% trypsin-EDTA (Life Technologies, 25300054). After the cells detached from the flask, 30 mL of medium was added, cell number was counted, and cells were centrifuged at 300 RCF for 5 min. After washing with PBS, a final collection at 300 RCF for 5 min was performed prior to storage at -80°C or addition of RLT plus beta-ME buffer.

RNA/DNA extraction and cDNA synthesis: Total RNA and DNA were extracted from cells using an AllPrep DNA/RNA mini Kit (Qiagen, 80204), following the manufacturer’s protocol including the on-column DNase digestion. For RNA, a second DNase treatment was performed using the RNeasy MinElute cleanup kit (Qiagen, 74204) following the manufacturer’s protocol, and DNA was cleaned up using a DNeasy PowerClean Cleanup kit (Qiagen, 12877-50) following the manufacturer’s protocol. Each RNA and DNA sample was quantified by Nanodrop for RNA and DNA quality and by Qubit to obtain RNA and DNA exact concentration.

First-strand cDNA was synthesized from 30 μg or 180 μg of total RNA with SuperScript II Reverse Transcriptase (ThermoFisher, 18064014 for hSTR1 or dpSTR) or Superscript III First-Strand Synthesis SuperMix (Thermo Fisher, 18080400 for hSTR2) and a specific primer MPRA_TruSeq_Universal_Adapter_REV (5’-ACACTCTTTCCCTACACGACGCTCTTCCGATCT-3’) following the protocol from Gordon et al.[4], except the total reaction volume was scaled to 50 μL for hSTR1 or dpSTR and 300 μL for hSTR2.

Generating the MPRA expression library for NGS: MPRA expression libraries were prepared from cDNA (RNA) and DNA by performing two rounds of PCR. For the hSTR1 and dpSTR expression libraries, 5 μL (3 μg of RNA) out of 50 μL of cDNA and 6 μg of DNA were used for the first PCR in a total reaction volume of 50 μL containing 25 μL 2X PrimeSTAR max (Takara, R045B), 0.5 μM of MPRA_Illumina_GFP_FWD

(5’-ACTGGAGTTCAGACGTGTGCTCTTCCGATCTCGCCCTGAGCAAAGACC-3’) and MPRA_TruSeq_Universal_Adapter_REV (5’-ACACTCTTTCCCTACACGACGCTCTTCCGATCT-3’). Samples were amplified with the following PCR conditions: 98°C for 1 min, 6 cycles (98°C for 10 sec, 58°C for 10 sec, 72°C for 20 sec), 72°C for 2 min and hold 4°C. Amplified cDNA and DNA was used for second round PCR without purification.

For hSTR2, all 300 µL (180 µg of the total RNA) of the cDNA and 36 µg of DNA were used for the first PCR. For hSTR2, the reaction volume (1200 µL) also included 600 µL 2X PCR NEBNext Ultra II Q5 Master Mix (NEB M0544) and 0.5 µM of MPRA_Illumina_GFP_FWD and MPRA_TruSeq_Universal_Adapter_REV. Each 50 µL reaction was loaded into a 96 well plate. Samples were amplified with the following conditions: 98°C for 30 seconds, 3 cycles (98°C for 10 sec, 65°C for 20 sec, 72°C for 30 sec), 72°C for 2 min and hold 4°C. Individual amplified cDNA and DNA were purified by 12% PEG Speedbeads and eluted in 170 µL of EB.

To estimate the cycle number for the second PCR, we used 5 µL of the unpurified (hSTR1 and dpSTR) or purified (hSTR2) first PCR product in a 20 µL qPCR reaction. The qPCR reaction contained: 10 µL 2X PrimeSTAR max (hSTR1 and dpSTR), or 2X PCR NEBNext Ultra II Q5 Master Mix (hSTR2), 2 µL 10X Sybr green I (Life Technologies, S-7567) and 0.5 µM of indices i5/i7. The conditions used: 98°C for 1 min, 40 cycles (98°C for 10 sec, 58°C for 10 sec, 72°C for 30 sec), 72°C for 5 min, 60-95 °C in 15 min and hold 4°C (hSTR1 or dpSTR); or 98°C for 1 min, 30 cycles (98°C for 10 sec, 60°C for 30 sec, 72°C for 1 min,) 72°C for 2 min, 60-95 °C in 15 min and hold 4°C (hSTR2). Based on the raw amplification curve of each sample, the number of cycles at which the amplification nearly plateaus for each sample were determined following Gordon *et al.*[4]. Since the numbers of cycles required for cDNA (RNA) and DNA samples were different, the second round PCR was run separately.

<u>Incorporation of sequencing indices:</u> 5 µL of the first PCR product from each sample was used in a PCR reaction with 2X PrimeSTAR max in 50 µL of PCR reaction volume for hSTR1 and dpSTR or for hSTR2, the entire 150 µL elution in a total reaction volume to 600 µL containing 300 µL 2X PCR NEBNext Ultra II Q5 Master Mix (NEB M0544). 0.5 µM of different indices i7/i5 were added for each sample and amplified with the same PCR conditions as used for the qPCR detailed above. Each reaction was loaded into a 96 well plate. For hSTR1 or dpSTR, each indexed library was run on a gel, then the correct size band (291bp) was cut and gel extracted

using a Zymoclean Gel DNA Recovery kit (11-301) and eluted with a 10 µL of Elution buffer or ultrapure water. For hSTR2, the 550 µL reaction was purified by 12% PEG Speedbeads, then eluted in 60 µL IDTE buffer. Three replicates of indexed cDNA (RNA) or DNA libraries were pooled separately using an equal amount (1 µg) for each replicate. The resulting pools were run on a 2% agarose gel. The correct size band was cut and gel extracted using a Zymoclean Gel DNA Recovery kit (11-301) and eluted with a 10 µL of Elution buffer or ultrapure water. Each mixed cDNA (RNA) and DNA indexed library was purified by 12% PEG Speedbeads, then eluted in a 20 µL IDTE buffer. Each indexed library was quantified by Nanodrop and Qubit was used to determine the indexed library concentration. The DNA size (291bp) was determined by running 15-30 ng of mixed cDNA and DNA on a 1.8-2% gel. DNA and RNA samples were pooled in a single LoBind tube with 1:1 ratio to obtain a 50 µL mixture at the final concentration of 10 nM.

#### *Library Sequencing on Illumina or Element Machines*

##### *1. Sequencing on the iSeq 100 System*

Prior to sequencing on the NovaSeq 6000 instrument and other instruments, the iSeq 100 system was used for library and pool QC. Thawed iSeq 100 kits (Illumina, 20021533), were used following Illumina instructions. Each library was diluted to a 1 nM concentration, and we pooled equal volumes of each 1 nM normalized library and determined the concentration of the total pool by Qubit. We then diluted the 1 nM library pool to 50 pM with 10 mM Tris-HCl, pH 8.5. PhiX was also diluted to 50pM. We then prepared 40 µL of 20% Phix and 80% pooled library and loaded 20 µL of the mixture to the sample reservoir of an iSeq100 cartridge and ran 2 x 150 paired-end cycles.

##### *2. Sequencing on the NextSeq 500/550 System*

We used NextSeq 500/550 High Output Kit v2.5 (75 or 150 Cycles) (Illumina, 20024907) for STR association libraries. We thawed HT1 at RT and stored at 4ºC until needed. We diluted each library and Phix to a 4 nM, with 10 mM Tris-HCl, pH 8.5 with 0.05% Tween 20, and pooled each library needed with 20% of PhiX for 1 nM. We measured the pooled library by using Qubit. We denatured the pooled library with fresh 0.2N NaOH, and diluted denatured libraries to 20 pM and made final dilution to loading concentration of 1.8 pM. We loaded 1300 µL of the diluted library at 1.8 pM to the sample reservoir of the cartridge and started a run on the NextSeq 500/550.

##### *3. NextSeq 2000 Systems*

We used NextSeq 2000 P3 Reagents (50 Cycles) (Illumina, 20046810) or NextSeq 2000 P4 Reagents powered by XLEAP-SBS™ chemistry (50 Cycles) (Illumina, ref# 2101842). We normalized and pooled all the libraries as we did above to 2 nM in a LoBind microcentrifuge tube, using the resuspension buffer (RSB) Tween. We quantified the pool on Qubit and calculated the volume of each library to generate a 2 nM library pool and added 2 nM PhiX to a final volume of 25 µL. That was diluted to 750 pM with RSB with Tween for the P3 kit and 560 pM with RSB with Tween for the P4 kit. We added 20 µL of the diluted library to the bottom of the sample reservoir.

##### *4. NovaSeq 6000 system*

The pool library was submitted to UCSD Institute of Genomic Medicine (IGM) center following their instruction and was run on Illumina NovaSeq 6000 system by UCSD IGM center using either PE100 or PE150 kits.

##### *5. Element sequencing*

The pool library was submitted to Element Bioscience and ran on Cloudbreak Freestyle kits with two indices of 8bp, a read1 length of 25bp and a read2 length of 135bp by using AVITI 2x75 Sequencing Kit Cloudbreak FS High Output (Element Bioscience 860-00015, 184 cycles and 1 billion reads). No PhiX was added to the pooled library. The lowest library concentration was 35 nM in 15 µL.

#### *RNase H1 Knockdown*

Generating gRNA donor backbone vector (PB_gRNA_S.pyogenes_scaffold_BB): The gRNA donor backbone vector was generated in our lab starting with the vector: PB-U6insert-EF1puro (Addgene plasmid #104537). Custom DNA Ultramer Oligos were ordered from IDT (4 nmole) containing BbsI recognition sites (underlined) upstream of the *S. Pyogenes* gRNA scaffold sequence (**bolded**), in order to generate a backbone vector where gRNA protospacers can be inserted directly upstream of the scaffold sequence through Golden Gate cloning. The top and bottom strand Ultramer Oligos were designed such that the double stranded annealed product would contain an AgeI restriction digest overhang on the 5' end of top strand (*italicized*) and an EcoRI restriction digest overhang on the 5' end of the bottom strand (*italicized*).

Top and bottom strand Ultramer Oligo sequences:

| | |
|---|---|
| top strand 5'-3' | *CCGG*GTCTTCATTAATGGAAGACAC**GTTTAAGAGCTAAGCTGGAAACAGCATAGCAAGTTTAAATAAGGCTAGTCCGTTATCAACTTGAAAAAGTGGCACCGAGTCGGTGCTTTTTTT** |
| bottom strand 5'-3' | *AATT***AAAAAAAGCACCGACTCGGTGCCACTTTTTCAAGTTGATAACGGACTAGCCTTATTTAAACTTGCTATGCTGTTTCCAGCTTAGCTCTTAAAC**GTGTCTTCCATTAATGAAGAC |

The PB-U6insert-EF1puro vector was digested with AgeI-HF and EcoRI-HF followed by gel extraction. The top and bottom Ultramer Oligos were annealed together and the product was ligated into the digested and gel extracted PB-U6insert-EF1puro vector using T4 DNA ligase (NEB cat. M0202L) (100ng of vector and 18.6ng of insert). The final gRNA donor backbone vector uses PiggyBac (PB) integration and puromycin selection to generate stable cell lines, and is called PB_gRNA_S.pyogenes_scaffold_BB (Addgene plasmid #226429).

Generating the RNase H1 CRISPRi gRNA Donor Vector Library (protocol adapted from Martin-Rufino et al.[5]): Six protospacer sequences targeting the RNase H1 promoter were chosen from Human CRISPR Inhibition Pooled Library (Dolcetto)[6]. Oligos containing the protospacer sequence (**bolded**) flanked by BbsI recognition sites (underlined) were ordered from IDT, along with a reverse primer for oligo amplification.

Protospacer Oligos and Oligo Amplification Primer:

| | |
|---|---|
| RNASEH1_1 | CTGAGCCgaagacCCCACCG**CCGGGAAGCATTTCGACTCC**GTTTTGgtcttcATGCTGCA |
| RNASEH1_2 | CTGAGCCgaagacCCCACCG**GCCAGCTCATCGCTCACTCC**GTTTTGgtcttcATGCTGCA |
| RNASEH1_3 | CTGAGCCgaagacCCCACCG**TTTCGACTCCCGGCCCAGCG**GTTTTGgtcttcATGCTGCA |
| RNASEH1_4 | CTGAGCCgaagacCCCACCG**GTCGAAATGCTTCCCGGTGC**GTTTTGgtcttcATGCTGCA |

| | |
|---|---|
| RNASEH1_5 | CTGAGCCgaagacCCCACCG**TCGAAATGCTTCCCGGTGCC**GTTTTGgtcttcATGCTGCA |
| RNASEH1_6 | CTGAGCCgaagacCCCACCG**TTCGACTCCCGGCCCAGCGT**GTTTTGgtcttcATGCTGCA |
| Oligo Amplification Primer | TGCAGCATgaagacCAAAAC |

Each protospacer oligo was resuspended to a 100 µM stock solution with IDTE. All six oligos were mixed together in equimolar concentration and diluted to a 10 µM solution of total oligo. PCR amplification of the oligos was performed with PrimeSTAR 2X mastermix (Takara Bio, R050A), using 2 µM final concentration of the mixed Protospacer Oligo pool and 4 µM final concentration of Oligo Amplification Primer. PCR cycling was done as follows: 2 mins at 98ºC, followed by 9 cycles of (30 sec at 64ºC then 20 sec at 70ºC) then 2 mins at 72ºC and hold at 4ºC. The 60bp Protospacer Oligo PCR product was gel extracted using Monarch® DNA Gel Extraction Kit (T1020S), and the cleaned product was used in a digestion and ligation cycling reaction. We used a 20:1 molar ratio of Protospacer Oligo PCR product to PB_gRNA_S.pyogenes_scaffold_BB.

Reaction set up:

| Reagent | 1 rxn (µL) |
|---|---|
| 10X T4 DNA ligase buffer (NEB, B0202S) | 1 |
| T7 DNA ligase (NEB, M0318) | 1 |
| BbsI-HF (R3539S) | 0.5 |
| PB_gRNA_S.pyogenes_scaffold_BB (35ng) | 1 |
| Protospacer Oligo PCR product (3.14ng) | 1 |
| UltraPure distilled water (Thermo Fisher, 10977015) to | 5.5 |
| Total | 10 |

Temperature cycling: 99 cycles of (2 mins at 37ºC then 5 mins at 16ºC) then 30 mins at 37ºC, then hold 4ºC until clean up. As soon as the reaction is finished cycling (to prevent re-ligation of backbone plasmid with its original cut out fragment) the digestion-ligation cycling reaction is cleaned using Zymo PCR Clean and Concentrate Kit (D4013). The cleaned RNase H1 CRISPRi gRNA Donor Vector Library product was transformed into NEB 10-beta Electrocompetent *E. Coli* (C3020) using heat shock transformation.

Transfecting HeLa S3 cell line expressing dCas9-tag-BFP-KRAB with RNase H1 CRISPRi gRNA Donor Vector Library: The HeLa S3 cell line expressing dCas9-tag-BFP-KRAB was a gift from Dr. Komor's lab at UC San Diego. The HeLa S3 cells were transfected with the RNase H1 CRISPRi gRNA Donor Vector Library as well a human PiggyBac transposase expressing vector pcsj533 (ref), using Lipofectamine 3000 Transfection Reagent (L3000015), at a 1:2.5 molar ratio of transposase vector to transposon vector (according to this user manual: https://www.systembio.com/wp/wp-content/uploads/2020/10/Manual_PiggyBac_System.pdf). One day after transfection, cells were passaged and 1µg/mL puromycin (Invitrogen cat. A1113803) was added to the media. Puromycin was replenished in cell media every other day until negative control (untransfected) cells were all dead (~5 days).

### Supplementary Figures

#### Supplementary Figure 1

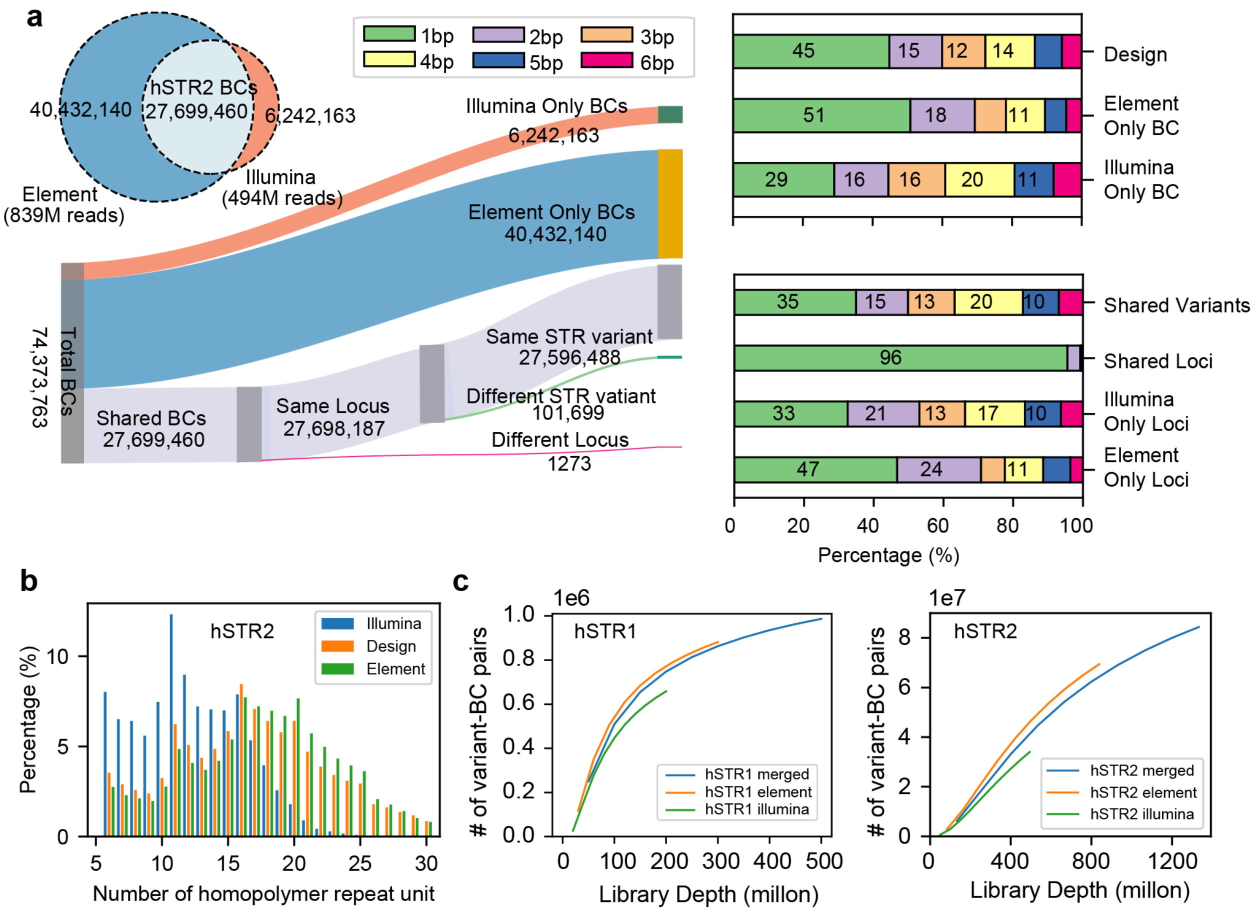


**Supplementary Figure 1 | Sequencing platform comparison and downsampling analysis of hSTR1 and hSTR2 libraries.**

**(a) Comparison of variant-BC pairs detected by NGS with Illumina and Element for hSTR2.** Left – a Venn diagram that depicts captured BCs using Illumina (orange) and Element (blue). The Sankey diagram expands the Venn diagram to show the concordance of BC assignments by each platform at the STR locus or variant level, since in some cases we observed agreement on the STR locus but disagreement on the copy number of the variant. Right – The percentages of all variants in hSTR2 by repeat unit length in the original oligonucleotide pool design (top). Subsequent rows show the breakdown by categories according to the Sankey diagram.

**(b) Percentage of homopolymers by copy number detected by each sequencing platform for hSTR2.** The *x*-axis shows the homopolymer copy number and the *y*-axis shows the percentage of all homopolymer variants with that length in the original design (light blue), vs. those captured by Illumina (orange) or Element (blue).

**(c) Downsampling analysis comparing detection rates.** The *x*-axes show the number of (million) reads and the *y*-axes show the number of variant-BC pairs for the moderate-complexity (hSTR1; left) and high-complexity (hSTR2; right) libraries, shown separately for Illumina-only (green), Element-only (orange), and merged (blue) sequencing data.

#### Supplementary Figure 2

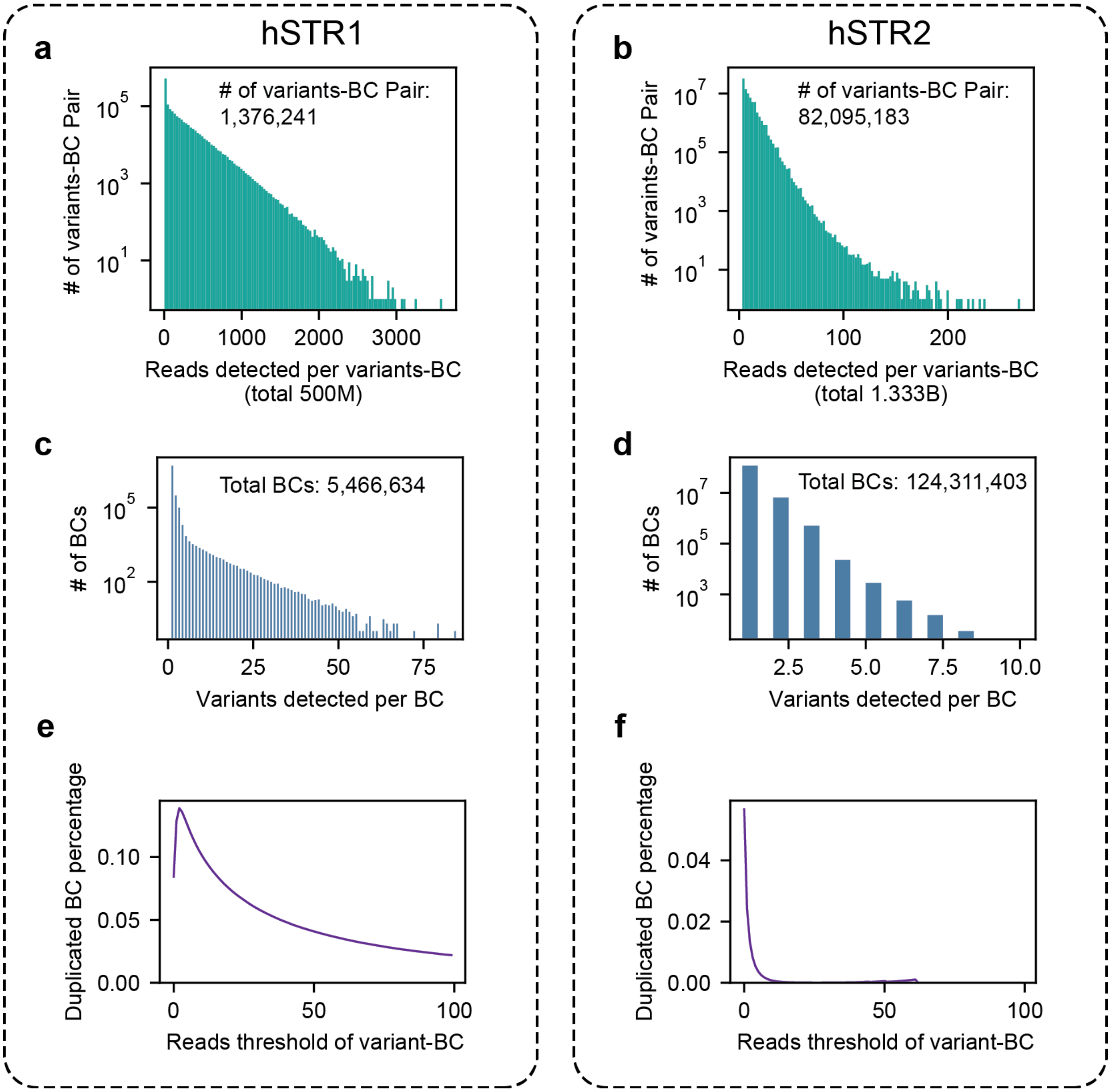


**Supplementary Figure 2 | Read count distribution and barcode duplication in hSTR libraries.**

**(a-b) Distribution of read counts per variant–barcode (BC) pair.** Data is shown for the moderate-complexity library (hSTR1, a) and the high-complexity library (hSTR2, b). Total read count annotated below x-axis.

**(c-d) Distribution of the number of variants detected per BC.** Plots show distributions for hSTR1 (c) and hSTR2 (d). Stutter error model applied when a single barcode was associated with variants of the same locus, otherwise barcodes associated with variants from multiple loci were filtered out (**Methods**).

**(e-f) Effect of read count thresholds on barcode duplication rates**. Plots show the distribution of the percentage of barcodes assigned to multiple variants in hSTR1 (e) and hSTR2 (f) as a function of the threshold used for minimum number of reads required for each variant-BC association.

#### Supplementary Figure 3

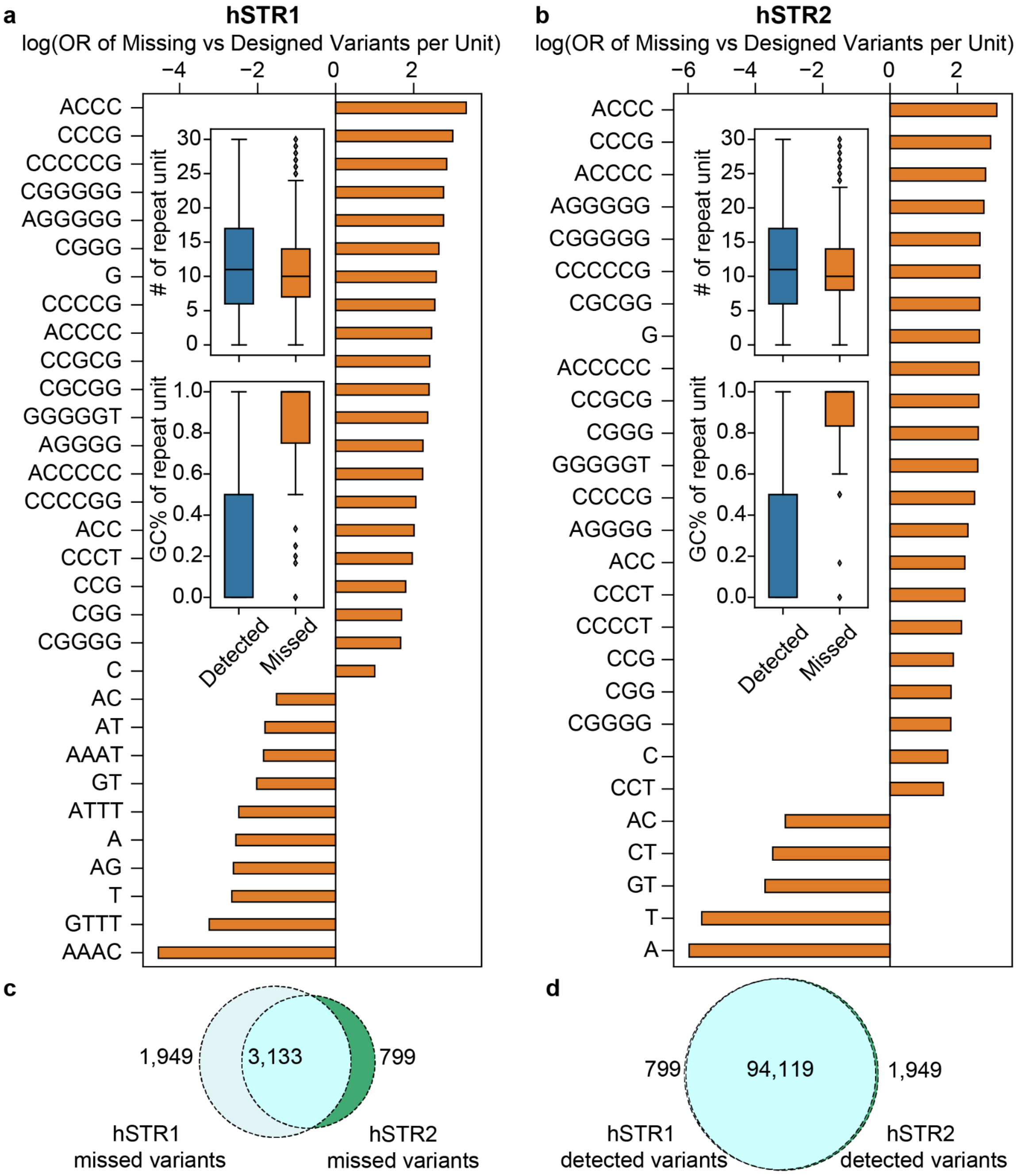


**Supplementary Figure 3 | Characterization of undetected variants and their features in MPRA libraries.** Bar plots display odds ratios (OR) for each repeat unit, comparing their representation among missing (undetected or filtered in our variant-BC association analysis) variants versus designed variants. Embedded above each bar is a box plot comparing the distributions of the number of repeat units (copy number) per variant between detected (blue) and missing (orange) variants; below each bar, a second box plot shows the distribution of GC percentage (GC%) of the repeat unit for detected versus missing variants. Data is shown for hSTR1 (a) and hSTR2 (b).

(c-d) Venn diagram depicts the overlap between variants in hSTR1 and hSTR2 that were missed (c) or detected (d). Light blue represents the variants that were only missed/detected in hSTR1, green represents variants only missed/detected in hSTR2, and cyan represents overlap between hSTR1 and hSTR2.

#### Supplementary Figure 4

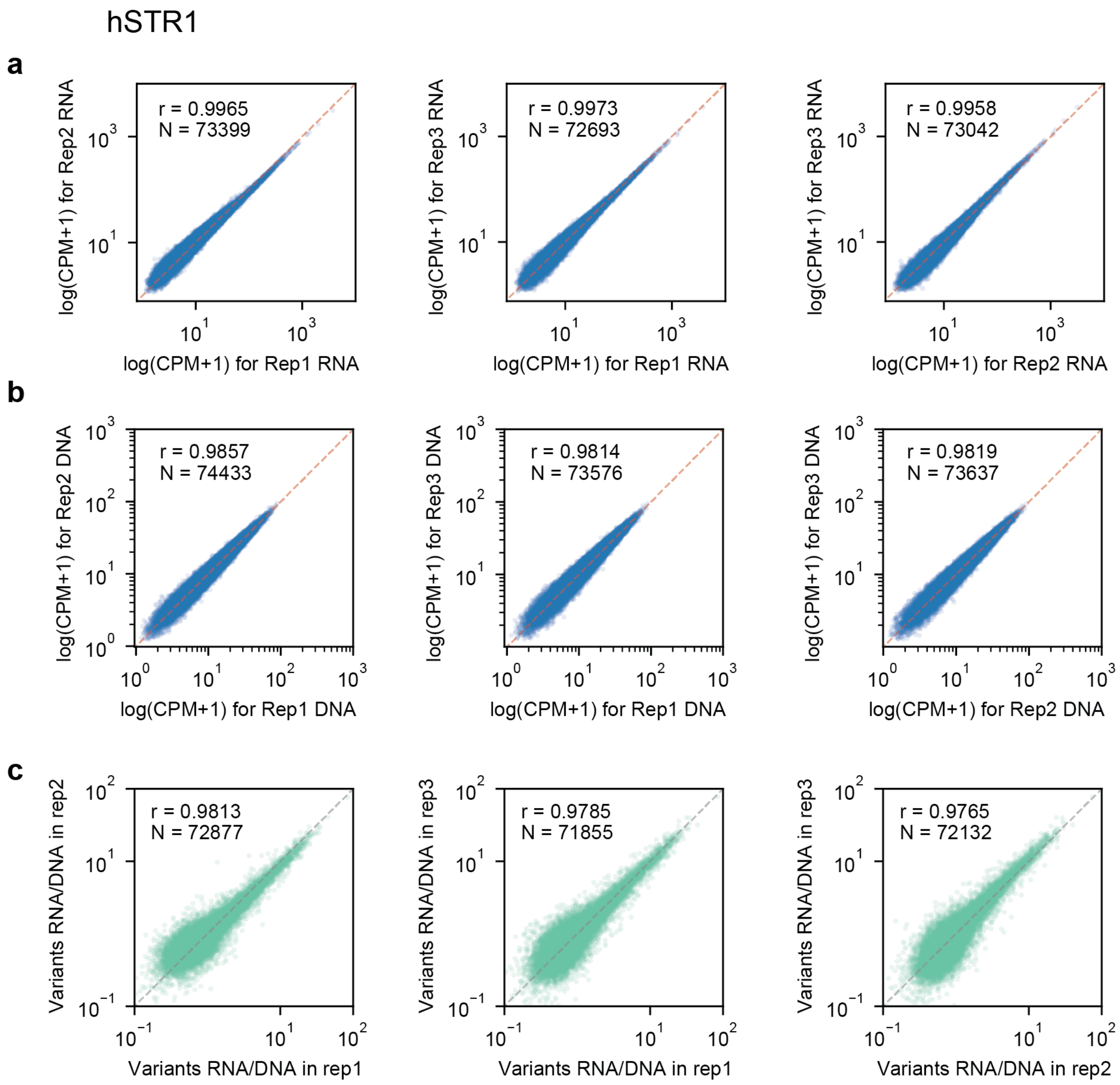


**Supplementary Figure 4 | Comparison of replicates for expression patterns in hSTR1.** Panels show the correlation of RNA counts per million (CPM) **(a)**, DNA CPM **(b)**, and RNA/DNA ratios **(c)** for each variant across replicates, based on RNA and/or DNA read counts summed across barcodes associated with each variant in each replicate. For all panels: left=rep1 vs. rep2, middle=rep1 vs. rep3, right=rep2 vs. rep3. Pearson correlation coefficient (r) and total variants detected across replicates (N) annotated. Dashed lines in each panel represent the $x=y$ line.

#### Supplementary Figure 5

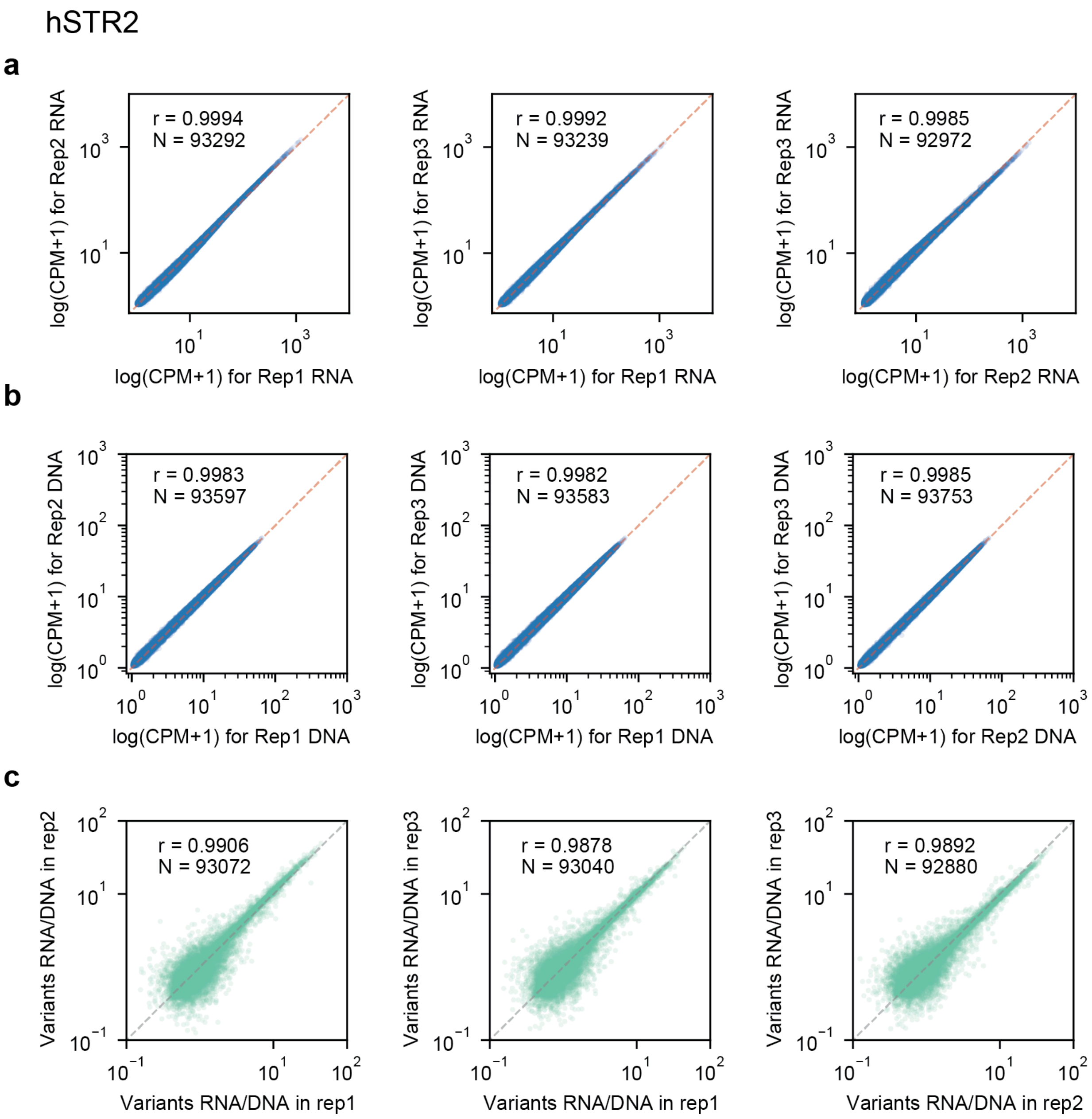


**Supplementary Figure 5 | Comparison of replicates for hSTR2 expression.** Panels show the correlation of RNA counts per million (CPM) **(a)**, DNA CPM **(b)**, and RNA/DNA ratios **(c)** for each variant across replicates, based on RNA and/or DNA read counts summed across barcodes associated with each variant in each replicate. For all panels: left=rep1 vs. rep2, middle=rep1 vs. rep3, right=rep2 vs. rep3. Pearson correlation coefficient (r) and total variants detected across replicates (N) annotated. Dashed lines in each panel represent the *x*=*y* line.

#### Supplementary Figure 6

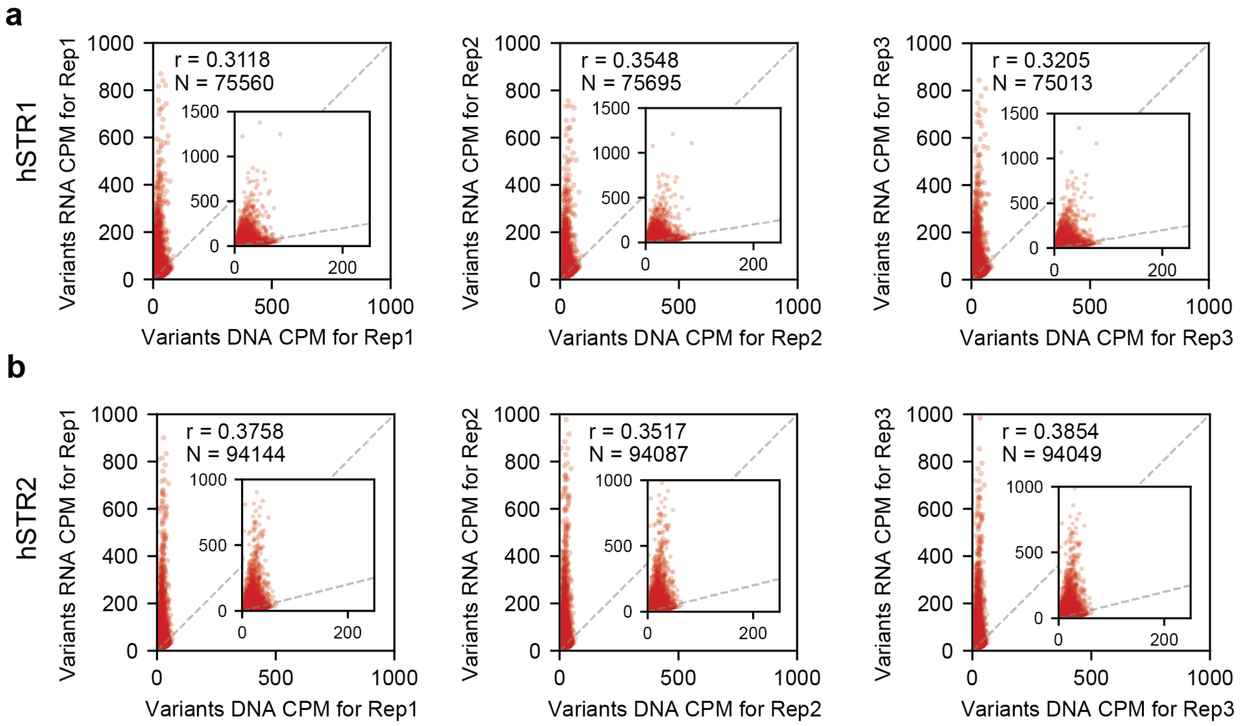


**Supplementary Figure 6 | Variant-level correspondence between DNA and RNA expression in hSTR libraries.** Each panel shows DNA vs. RNA counts per million (CPM) for each variant in each replicate, based on summing DNA and RNA reads across all barcodes associated with each variant. Data is shown for hSTR1 (top; **a**) and hSTR2 (bottom; **b**) for replicates 1-3 (left, middle, right). Inset plots show additional detail zoomed in to an x-axis range of 0-200. Dashed lines show the *x*=*y* diagonal.

### Supplementary Figure 7

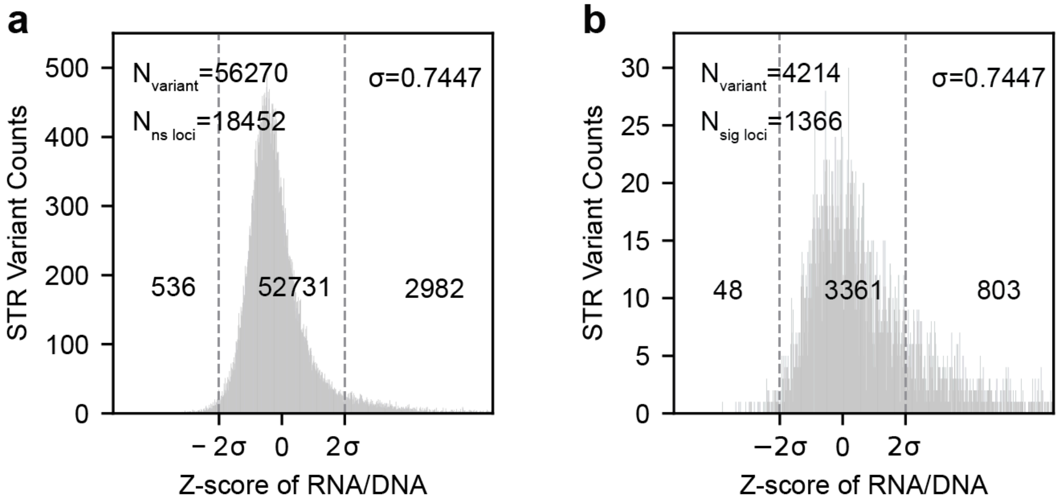


**Supplementary Figure 7 | Distribution of standardized RNA/DNA ratio (Z-scores) for variants grouped by regression locus significance.** Distributions are shown separately for all variants from loci with non-significant (**a**) and significant (**b**) associations between copy and expression.

### Supplementary Figure 8

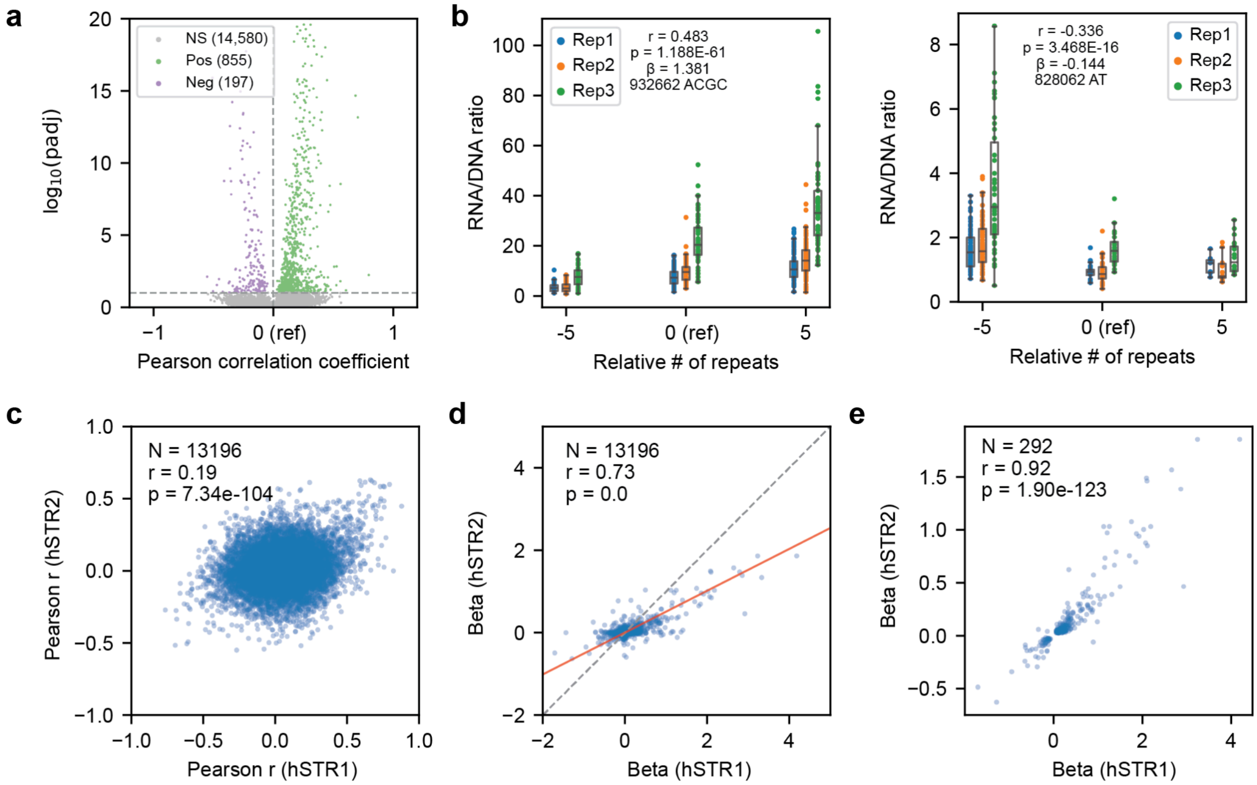


**Supplementary Figure 8 | Association of repeat length with reporter expression in hSTR2 and cross-library comparisons.**

**(a) Volcano plot summarizing associations between repeat copy number and reporter expression at each locus in hSTR2**. The *x*-axis shows the Pearson correlation between copy number and expression. The *y*-axis shows the log10 of the adjusted P-value. Purple=loci with a negative correlation between copy number and expression, green=loci with a positive association, grey=not significant.

**(b) Representative regression plots of examples with positive (left) and negative (right) associations.** Repeat copy number on the *x*-axes is shown relative to the reference genome (reference denoted by “0”). The *y*-axes denote RNA/DNA ratios. Each data point represents one barcode in one replicate and boxplots summarize distributions within each replicate. Regression slopes (β) and nominal p-values are annotated for each locus. Blue=replicate 1, orange=replicate 2, green=replicate 3.

**(c) Comparison of the correlation between copy number and expression across variants in hSTR1 vs. hSTR2.** Each data point represents one locus. Correlation is measured using Pearson’s r.

**(d) Comparison of regression effect sizes in hSTR1 vs. hSTR2 for all loci.** The diagonal gives the *x*=*y* diagonal. The red line gives the best fit line.

**(e) Comparison of regression effect sizes in hSTR1 vs. hSTR2 for significant loci.** The plot is the same as in **d** but only includes loci with significant associations in both hSTR1 and hSTR2.

### Supplementary Figure 9

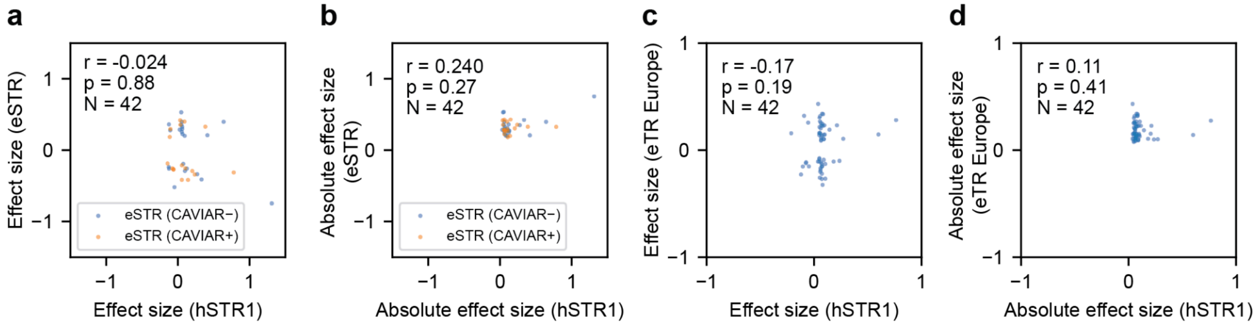


**Supplementary Figure 9 | Correlation of effect sizes between MPRA library hSTR1 and published expression STR (eSTR) population datasets.** Plots show comparisons of MPRA effect sizes for hSTR1 to those reported in GTEx by Fotsing *et al.*[7] (multiple tissues; **a-b**) or Europeans from the GEUVADIS cohort by Ziaei-Jam *et al.*[8] (lymphoblastoid cell lines; **c-d**). Only STRs with significant effects in both the MPRA and eSTR study are included. For the GTEx study (**a-b**), blue and orange represent STRs that failed and passed fine-mapping using CAVIAR, respectively. Panels **b** and **d** are the same as **a** and **c** but show absolute values of effect sizes.

#### Supplementary Figure 10

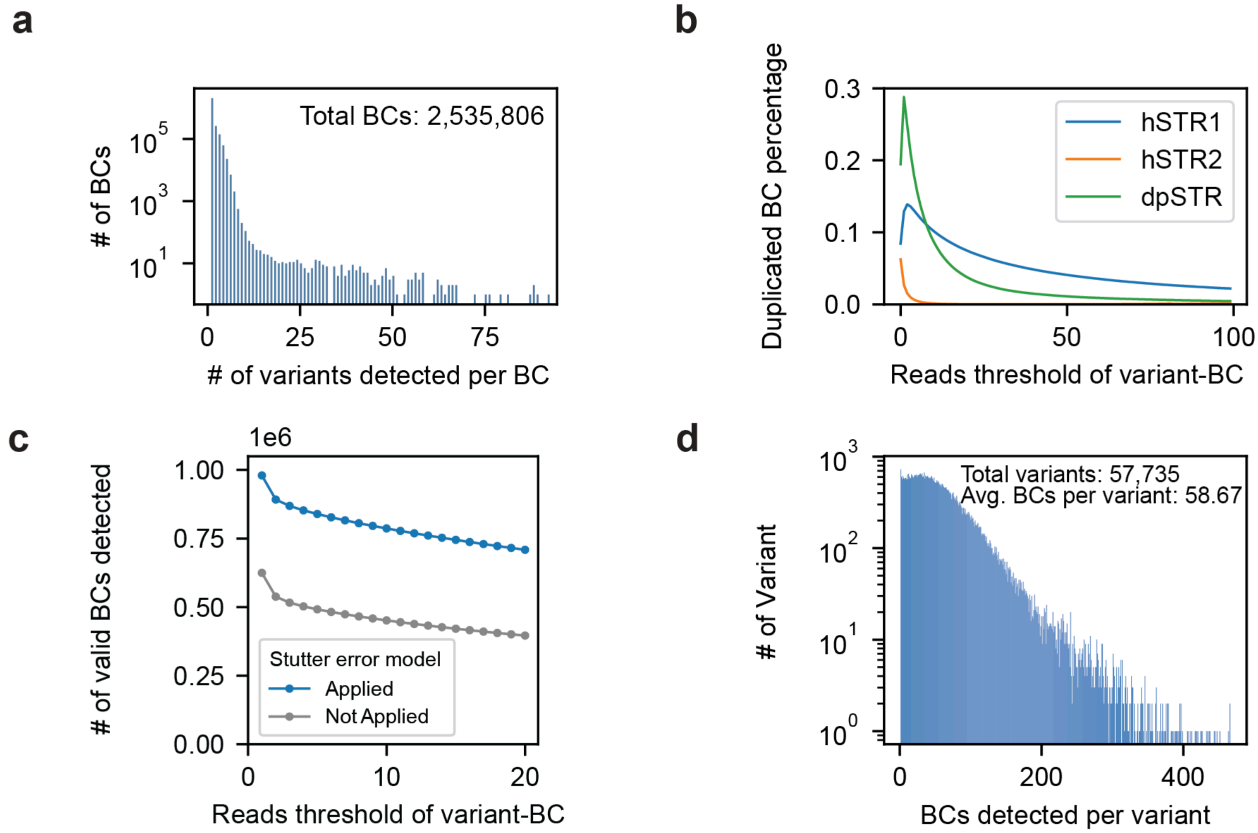


**Supplementary Figure 10 | Barcode detection and duplication analysis in the dpSTR library.**

**(a) Distribution of the number of barcodes (BCs) detected per variant in the dpSTR array.**

**(b) Duplicate BC percentage as a function of the threshold used to filter variant-BC pairs.** The *x*-axis shows the minimum number of supporting reads required to keep a variant-BC association. The *y*-axis shows the percent of passing BCs associated with more than one unique variant (duplicates). blue=hSTR1; orange=hSTR2; green=dpSTR.

**(c) Number of BCs detected at different read count thresholds in the dpSTR library, before and after application of the stutter correction model.** Gray=before stutter correction, blue=after stutter correction. The *x*-axis is the same as in **b.** The *y*-axis shows the number of passing BCs at each threshold. BCs are considering passing if they meet the minimum read support threshold and are not assigned to more than one variant.

**(d) Distribution of the number of BCs detected per variant in the dpSTR library after quality filtering.**

#### Supplementary Figure 11

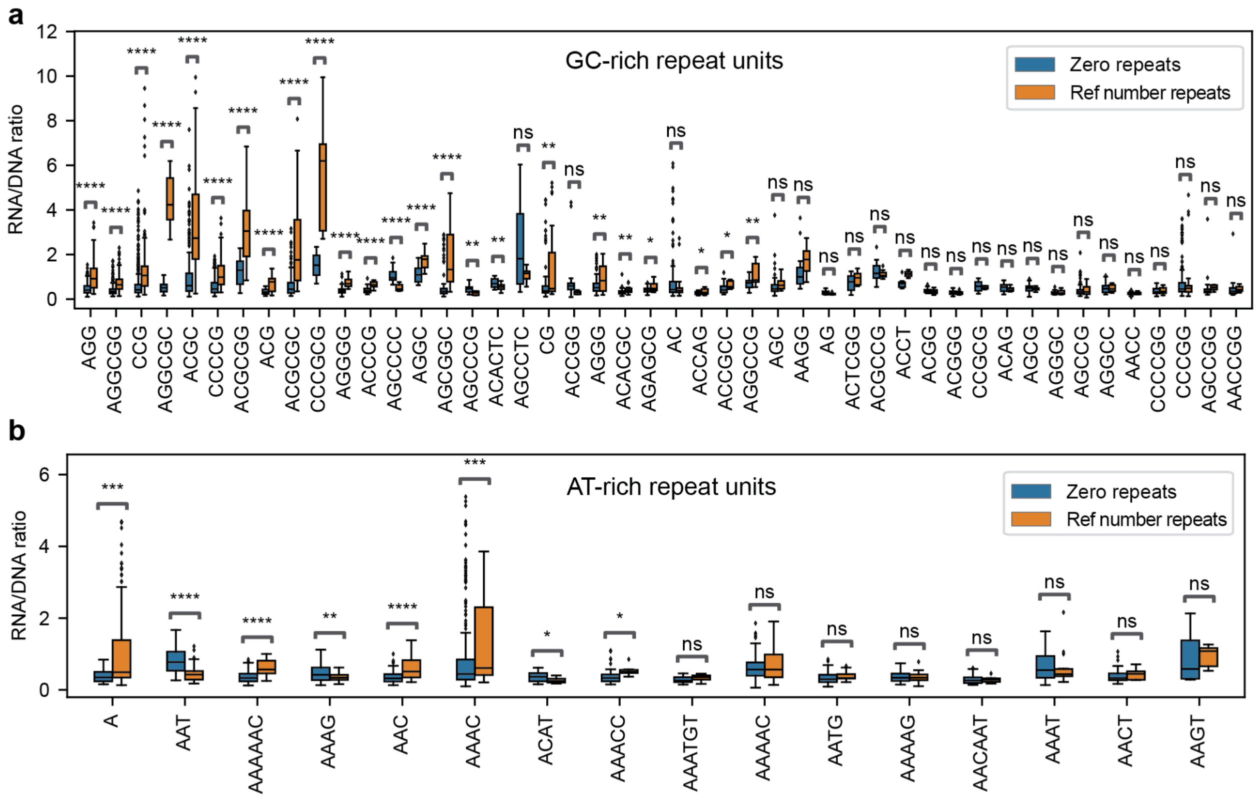


**Supplementary Figure 11 | RNA/DNA ratio comparison of the reference variant vs. deletion of the repeat (copy number=0) across repeat units in the dpSTR library.** Distributions are shown for GC-rich repeats (**a**) and AT-rich repeats (**b**). In each panel blue boxplots show distributions of RNA/DNA ratios across variants for which the repeat was deleted and orange shows distributions of ratios across variants with the reference sequence for each repeat unit. ns=not significant; *=1.00e-02 < p ≤ 5.00e-02; **=1.00e-03 < p ≤ 1.00e-02; ***=1.00e-04 < p ≤ 1.00e-03; ****=p ≤ 1.00e-04.

#### Supplementary Figure 12

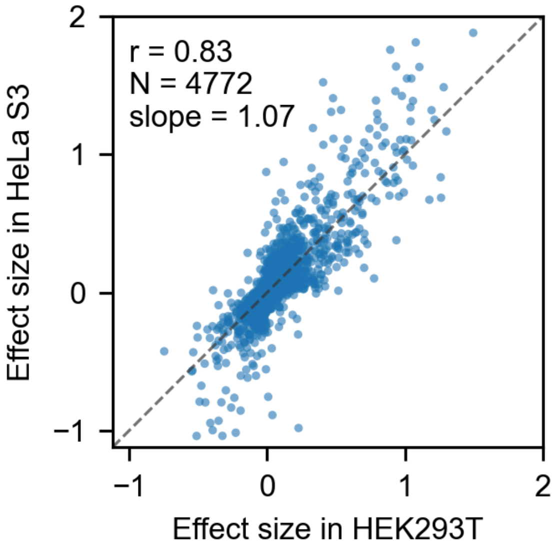


**Supplementary Figure 12 | Comparison of effect sizes measured for each locus/sequence pair in HEK293T and HeLa S3 wild-type cells for the dpSTR library.** P-value <1e-200. Dashed line gives the *x*=*y* diagonal.

### Supplementary Figure 13

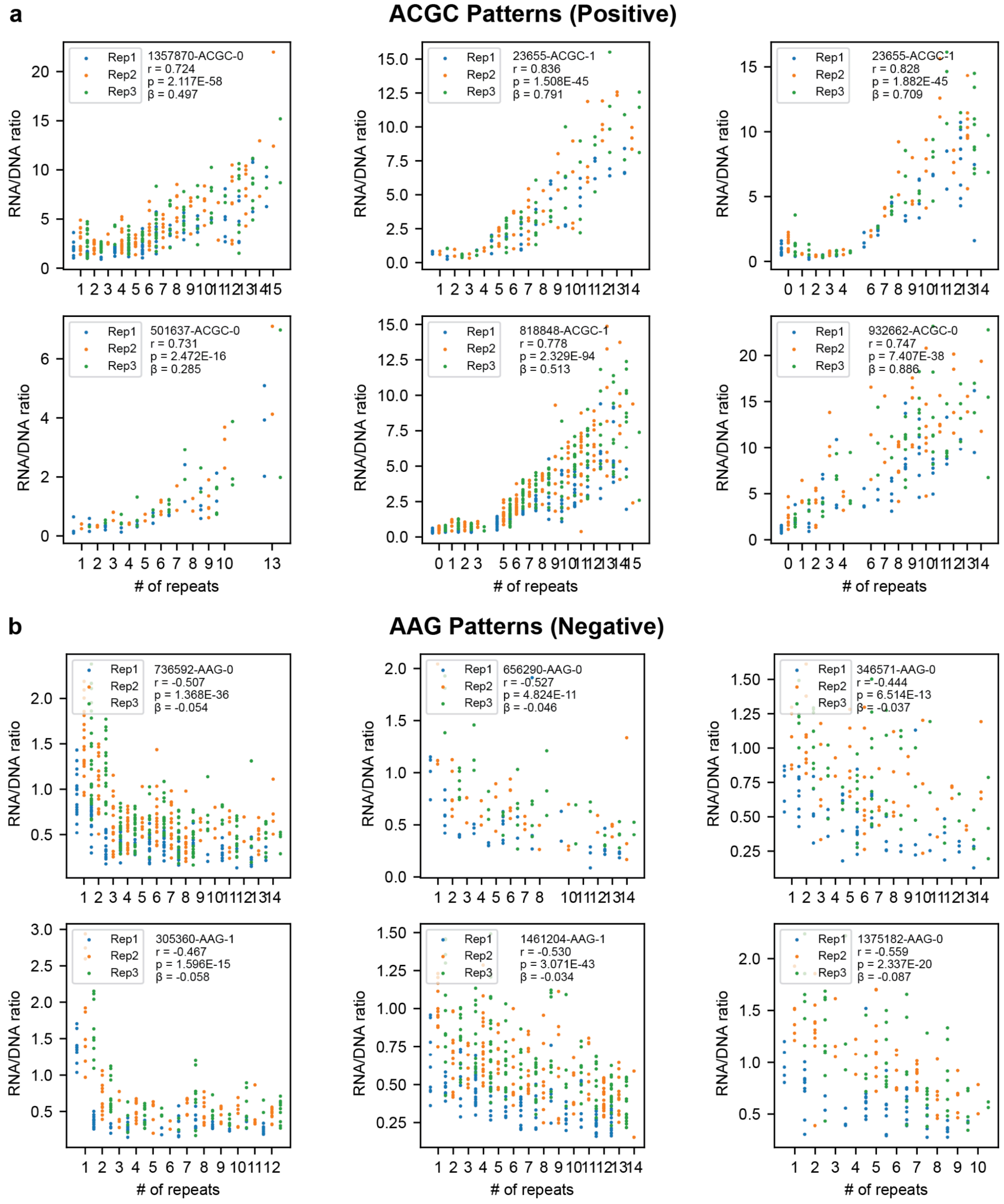

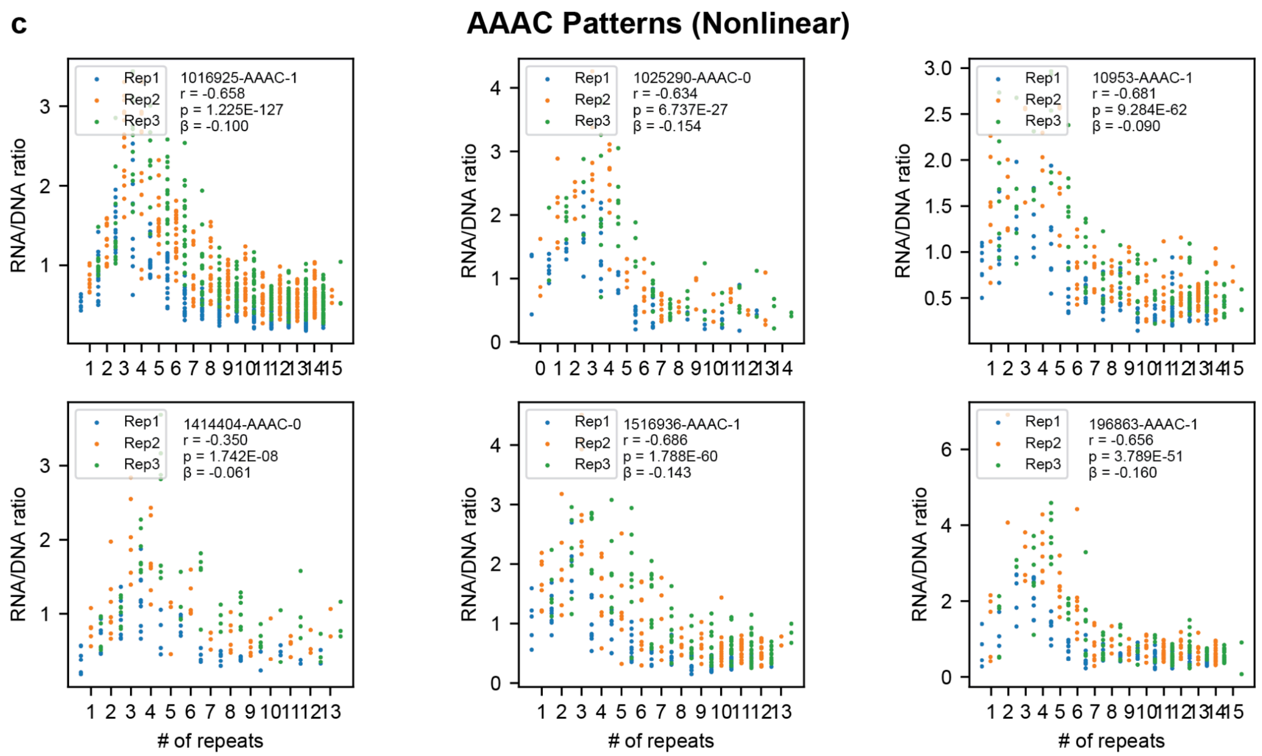


**Supplementary Figure 13 | Representative relationships between repeat copy number and RNA/DNA ratio for selected repeat units in the MPRA library.** Each panel shows data from a single repeat unit/locus pair in the dpSTR library. The *x*-axes give the total repeat copy number. The *y*-axes denote RNA/DNA ratios. Each data point represents one barcode in one replicate. Locus id, regression slope (β), nominal P-value, and Pearson correlation are annotated. Blue=replicate 1, orange=replicate 2, green=replicate 3. Panels in **(a-c)** show examples of ACGC, AAG, and AAAC repeats, respectively. Locus ids ending with "-0" indicate that the repeat was on the forward strand and those ending in "-1" indicate that the repeat was on the reverse strand (e.g. "AAAC-0" indicates AAAC whereas "AAAC-1" indicates TTTG repeats).

### Supplementary Figure 14

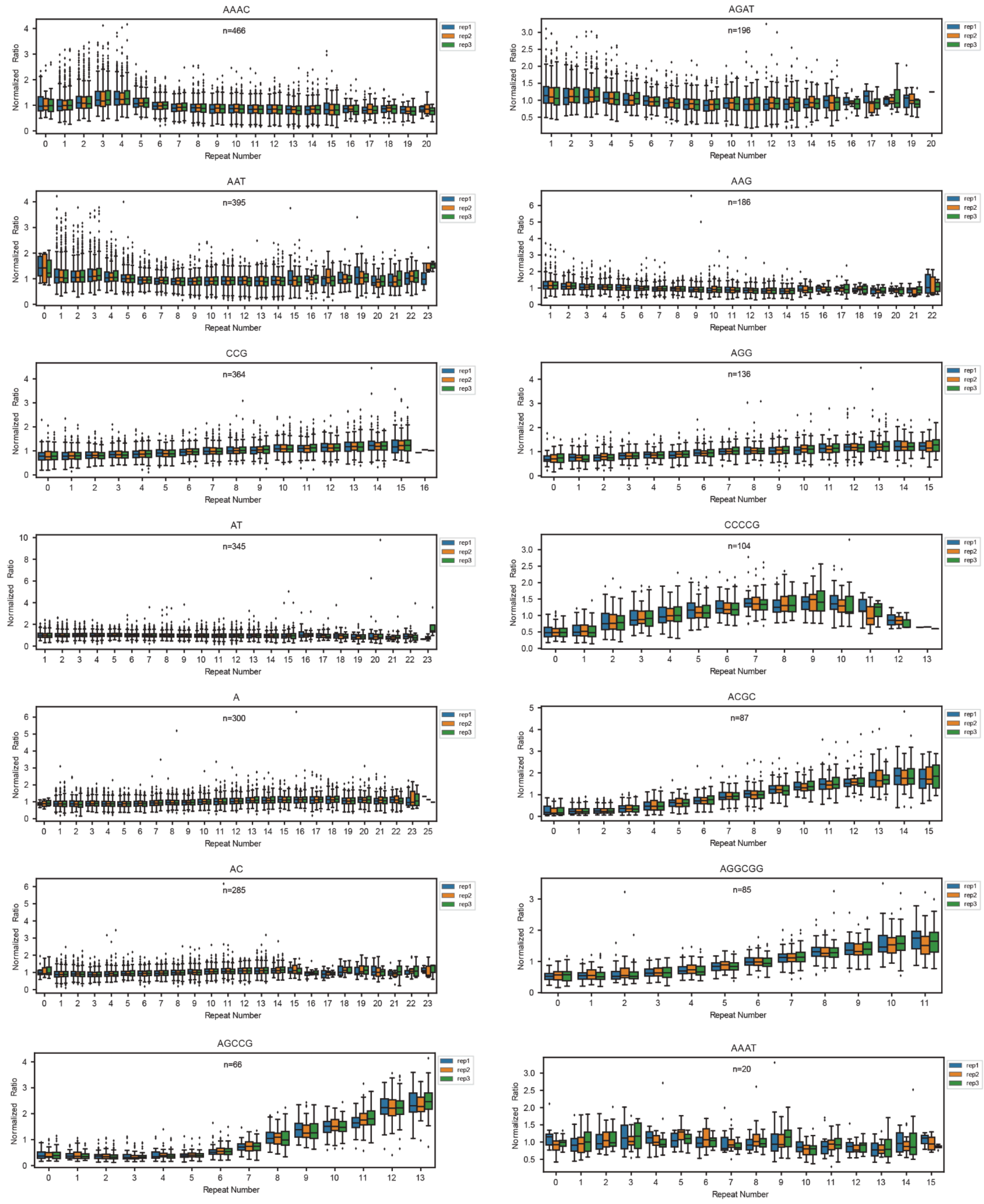

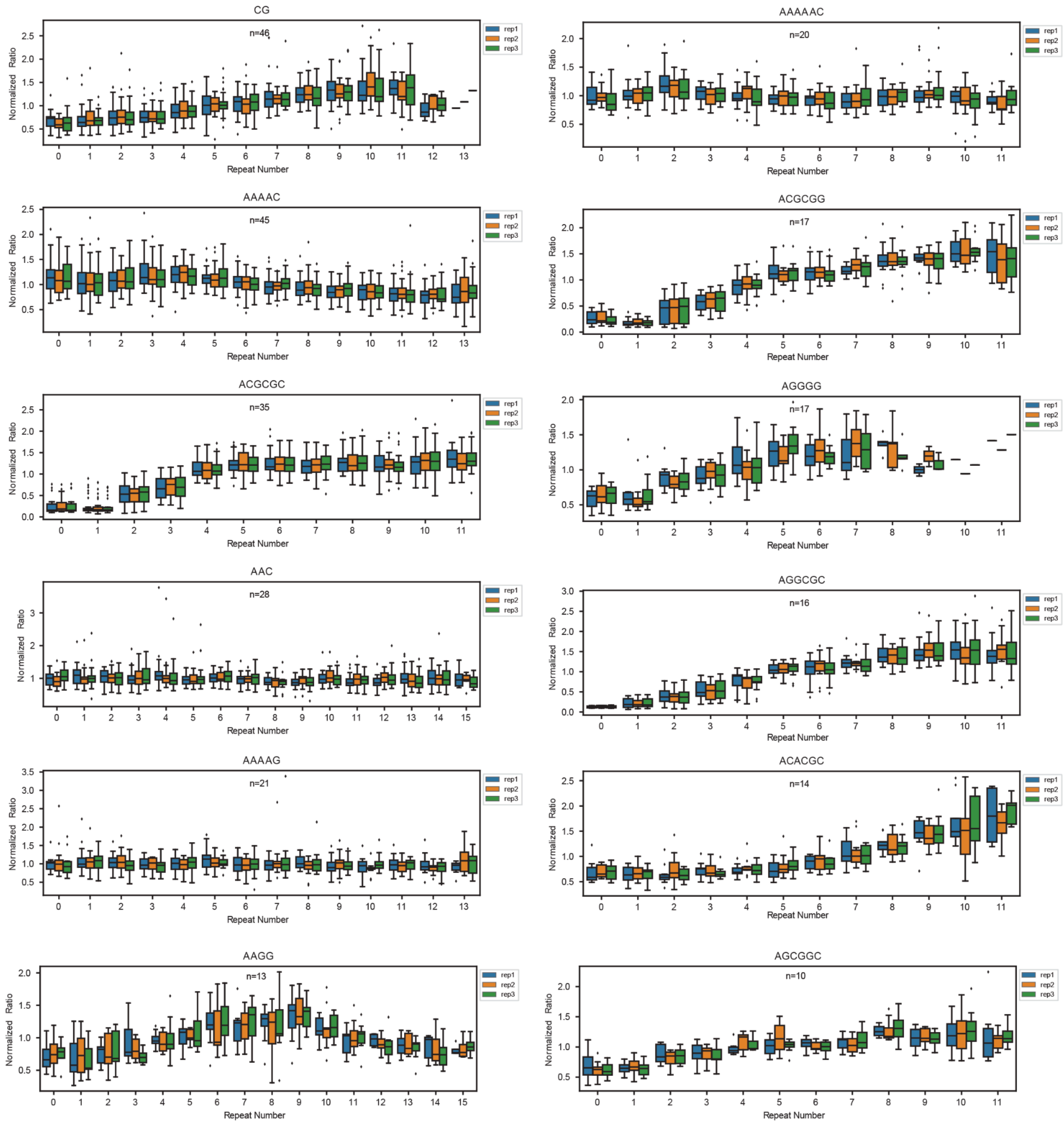


**Supplementary Figure 14 | Distribution of normalized expression (RNA/DNA ratios) by copy number across repeat units with ≥ 10 unique repeat unit/locus pairs.** All loci for which the indicated repeat unit was included as either the reference repeat unit or one of the perturbations are shown. Color indicates replicate number (blue=replicate 1, orange=replicate 2, green=replicate 3). The normalization procedure is described in **Methods.** Note, some loci might be tested with the same repeat unit twice if the reference contains that locus with imperfections, since we tested with and without the imperfection (**Methods**).

### Supplementary Figure 15

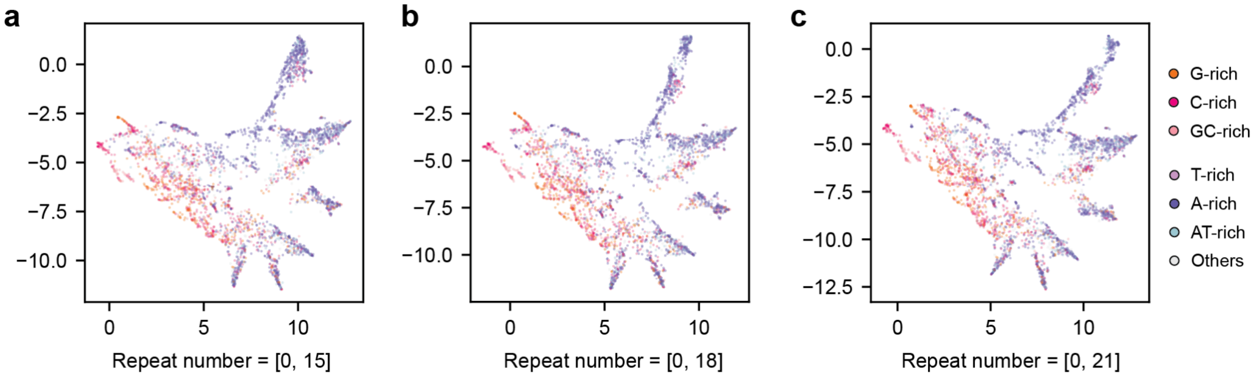


**Supplementary Figure 15 | Robustness of repeat unit pattern clustering visualized by UMAP with varying maximum repeat numbers.** The UMAP analysis shown in **Fig. 4f** was repeated including only copy numbers from 0-15 **(a)**, 0-18 **(b)**, or 0-21 **(c)** to ensure inferred patterns are robust to missing data due to the fact that not all repeat ranges were synthesized for all repeat units (**Methods**). Points are colored by repeat unit category as in **Fig. 4f**.

#### Supplementary Figure 16

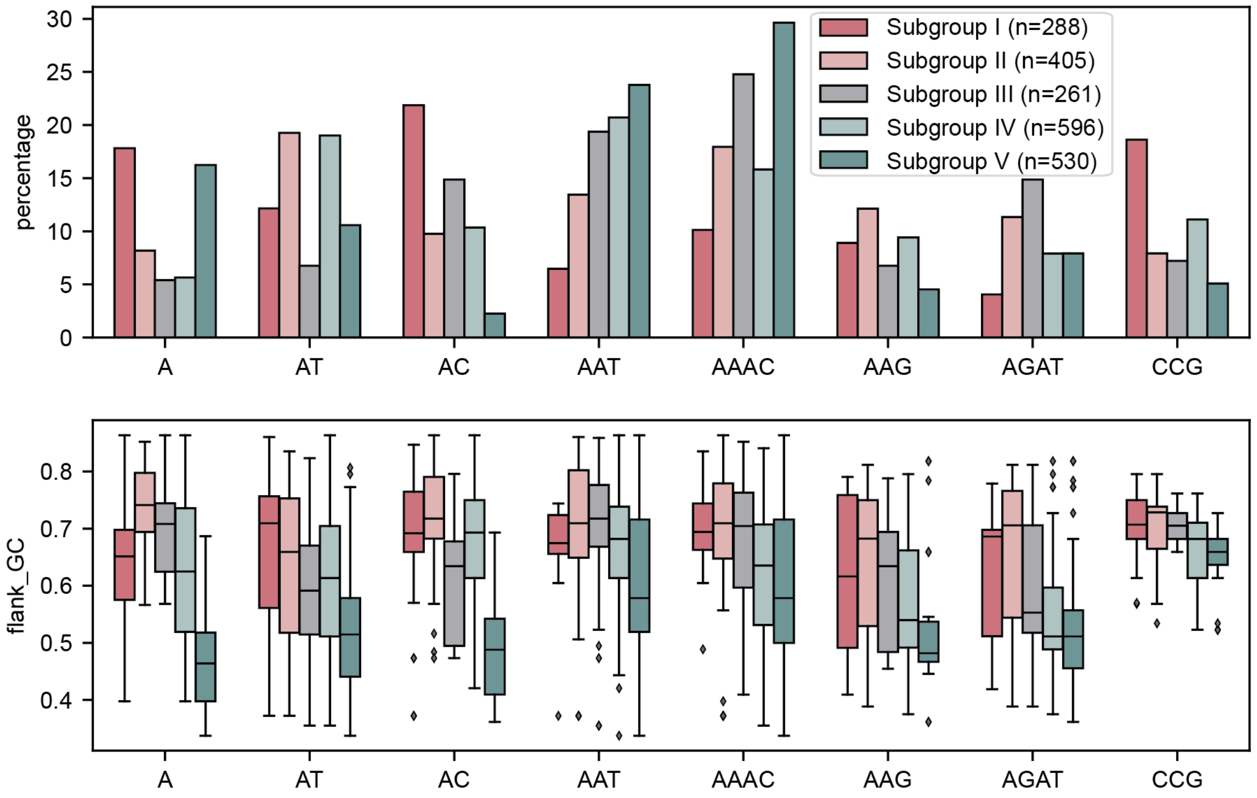


**Supplementary Figure 16 | Composition and flanking GC % distributions across repeat unit subgroups.** Top panel: Grouped bar plots showing the percentage of repeat unit/locus pairs in each subgroup with each repeat unit. Bottom panel: distribution of the flanking GC content for repeat unit/locus pairs in each subgroup grouped by repeat unit sequence. Subgroups divided from Group II in UMAP (Fig. 4f) by flanking GC% (Fig. 4i, j).

#### Supplementary Figure 17

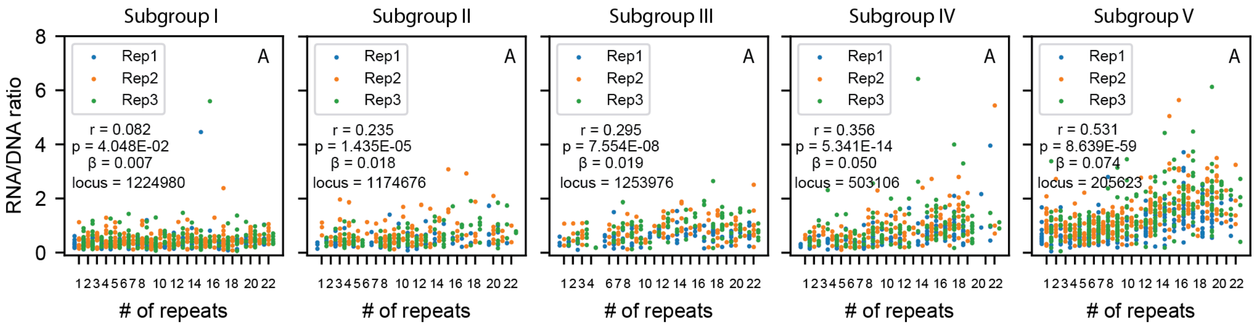


**Supplementary Figure 17 | Representative relationship between copy number and expression for examples with the repeat unit A across subgroups**. Subgroups are based on those defined in **Fig. 4f**. The *x*-axes give the total repeat copy number. The *y*-axes denote RNA/DNA ratio. Each data point represents one barcode in one replicate. Locus id, regression slope (β), nominal P-value, and Pearson correlation are annotated. Blue=replicate 1, orange=replicate 2, green=replicate 3.

### Supplementary Figure 18

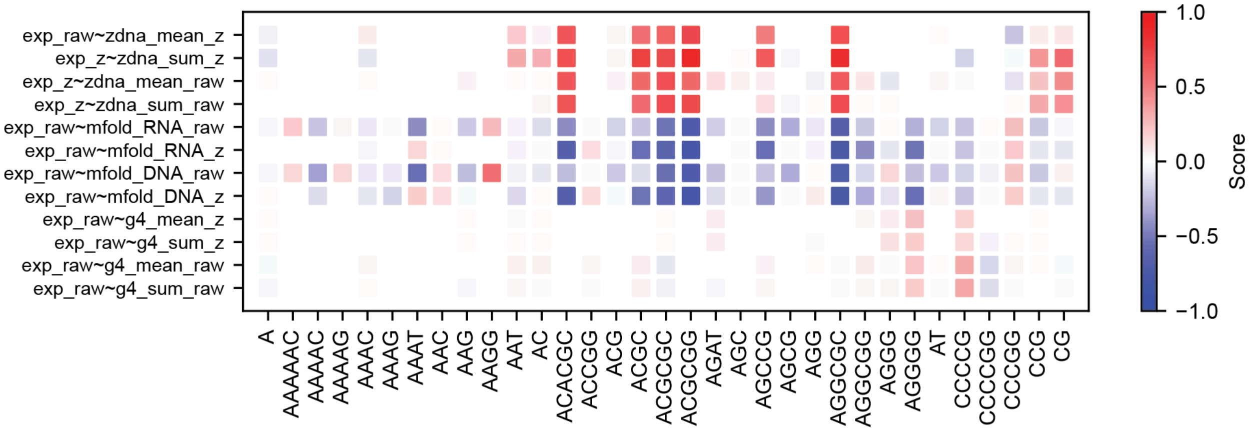


**Supplementary Figure 18 | Heatmap of correlations between RNA/DNA expression ratios and predicted structural scores across repeat units.** The *x*-axis shows repeat units. The *y*-axis labels indicate correlations between metrics based on RNA/DNA expression ratios and various predicted sequence-based structural features for variants associated with each repeat unit. Each cell in the heatmap shows the correlation coefficient between an expression metric and a structural score metric such as zDNA, mfold, or G4 predictions. Labels are in the format <expr_metric>~<structure_metric>, where expr_metric indicates whether we used raw or Z-normalized expression values and structure_metric indicates whether scores show predicted ZDNA, mfold energies based on either RNA or DNA sequences, or G4 scores. For example, "exp_raw ~ zDNA _sum_z" indicates a regression of raw expression ratios on the Z-score normalized sum of multiple zDNA scores per variant. Full details for different structure metrics computed are provided in **Methods**.

#### Supplementary Figure 19

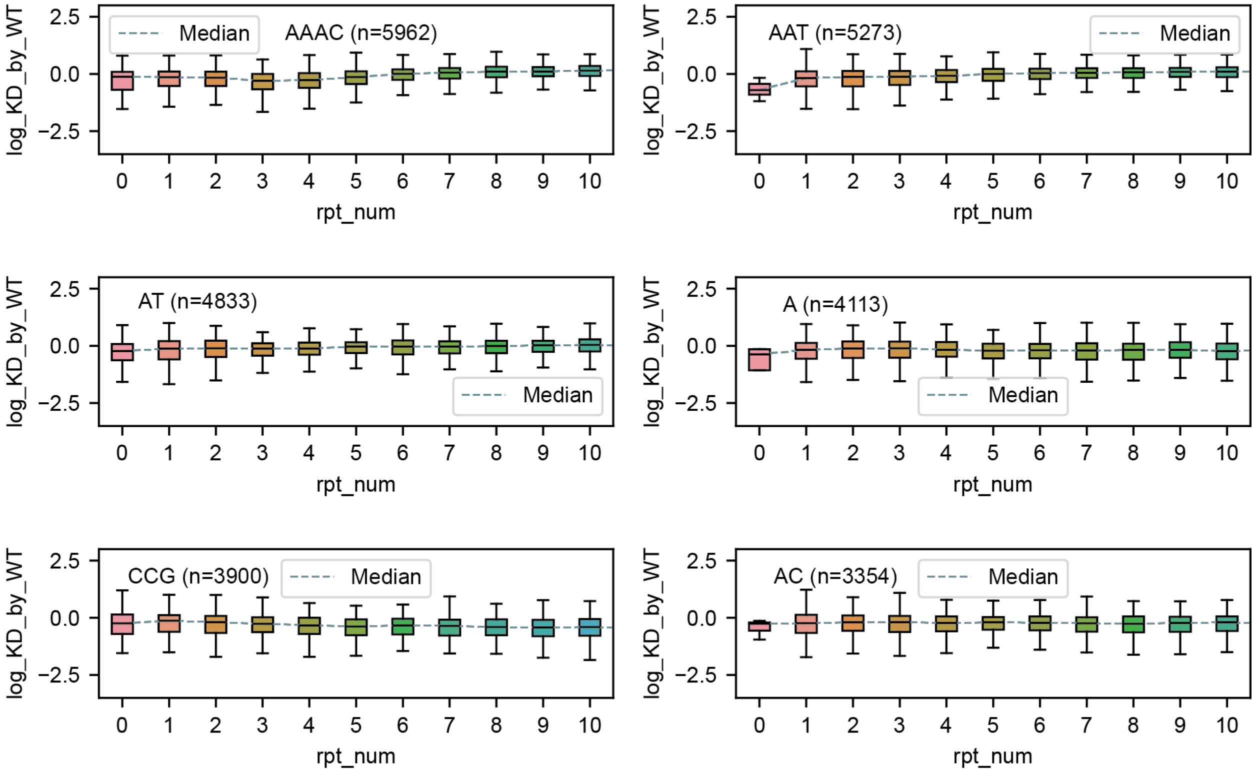


**Supplementary Figure 19 | Distribution of variant-level log(KD/WT) expression ratios stratified by repeat copy number for the six most abundant repeat units.** Panels display the distribution of log(KD ratio/WT ratio) values across repeat numbers for each of the six most abundant repeat units.

#### Supplementary Figure 20

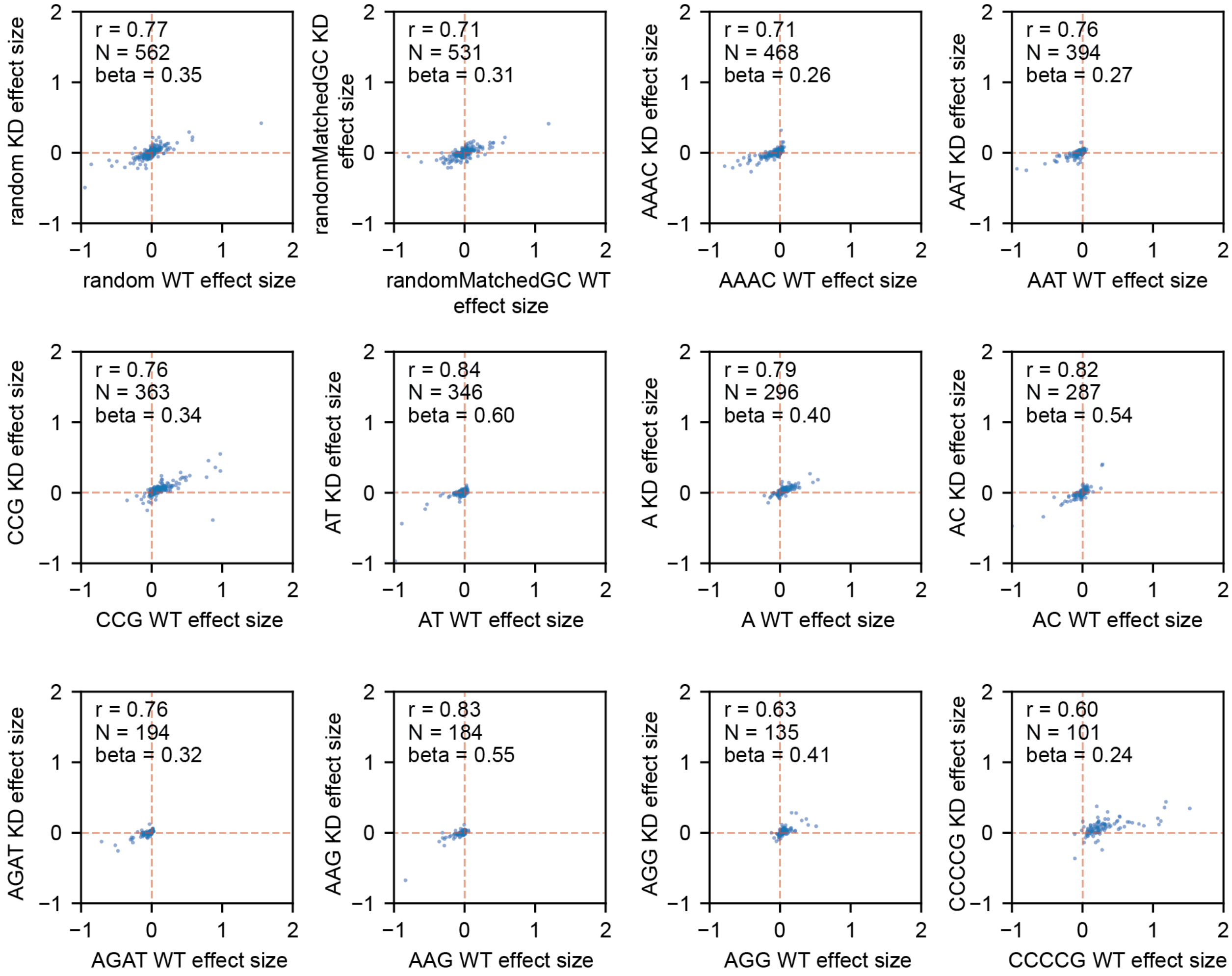


**Supplementary Figure 20 | Comparison of effect sizes obtained under WT and RNase H1 KD conditions for the most abundant repeat units.** Each plot compares effect sizes for a different repeat unit. All repeat units for which at least 100 repeat unit/loci pairs were included are shown.

#### Supplementary Figure 21

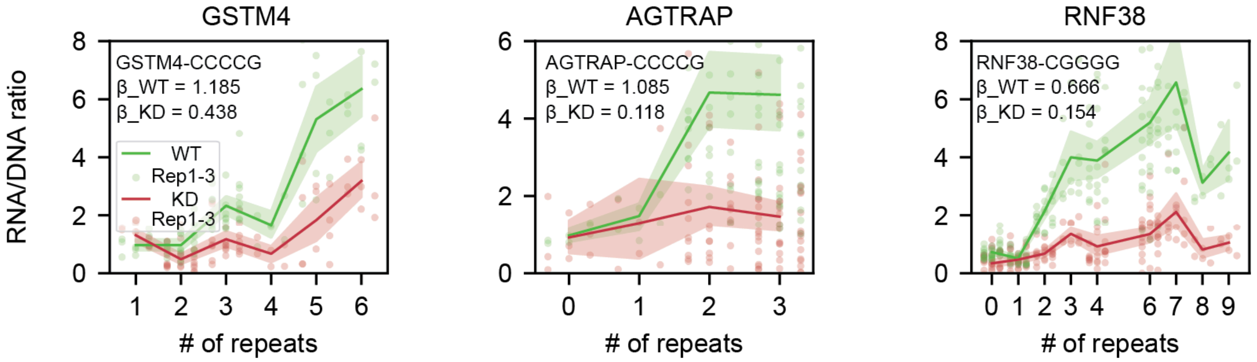


**Supplementary Figure 21 | Relationship between copy number and expression for examples with outlier changes in effect size in WT vs. KD conditions.** Plots are the same as in **Fig. 6f** but show the MPRA results for sequences with the reverse orientation (CCCCG while in **Fig. 6f** it was CGGGG). Green=WT, red=RNase-H1 KD, shaded regions indicate 95% confidence intervals.

### Supplementary Table Legends

**Table S1: Sequences used to generate STR MPRA libraries.** Sequences of oligonucleotide components, primers, and other sequences defined in **Methods** are provided.

**Table S2: hSTR1 and hSTR2 probe sequences.** This table contains all 100,000 oligonucleotide sequences with the following column definitions: variant=variant name; id=oligonucleotide ID; rpt_unit=repeat unit sequence; rpt_num_ref=number of repeats in hg38 reference; allele=copy number relative to reference (ref=reference allele, m5=reference minus 5 repeats, p5=reference plus 5 repeats, p3=reference plus 3 repeats); probe_seq=oligonucleotide sequence.

**Table S3: dpSTR sequences.** This table contains all 59,842 oligonucleotide sequences with the following column definitions: variant=variant name; Locus(id)=locus coordinates in hg38 (oligonucleotide ID); rpt_unit_original=repeat unit sequence in hg38; rpt_unit_current=repeat unit sequence in current variant; rpt_num=number of repeats in the variant; strand=orientation of current variant sequence relative to hg38; probe_seq=oligonucleotide sequence.

#### Supplementary References


1. Tewhey, R. *et al.* Direct Identification of Hundreds of Expression-Modulating Variants using a Multiplexed Reporter Assay. *Cell* **165**, 1519–1529 (2016).

2. Verma, V., Gupta, A. & Chaudhary, V. K. Emulsion PCR made easy. *BioTechniques* **69**, 421–426 (2020).

3. Abell, N. S. *et al.* Multiple causal variants underlie genetic associations in humans. *Science* **375**, 1247–1254 (2022).

4. Gordon, M. G. *et al.* lentiMPRA and MPRAflow for high-throughput functional characterization of gene regulatory elements. *Nat. Protoc.* **15**, 2387–2412 (2020).

5. Martin-Rufino, J. D. *et al.* Massively parallel base editing to map variant effects in human hematopoiesis. *Cell* **186**, 2456-2474.e24 (2023).

6. Sanson, K. R. *et al.* Optimized libraries for CRISPR-Cas9 genetic screens with multiple modalities. *Nat. Commun.* **9**, 5416 (2018).

7. Fotsing, S. F. *et al.* The impact of short tandem repeat variation on gene expression. *Nat. Genet.* **51**, 1652–1659 (2019).

8. Ziaei Jam, H. *et al.* A deep population reference panel of tandem repeat variation. *Nat. Commun.* **14**, 6711 (2023).
